# Unified nonparametric analysis of single-molecule spatial omics data using probabilistic indices

**DOI:** 10.1101/2025.05.20.654270

**Authors:** Stijn Hawinkel, Xilan Yang, Ward Poelmans, Hans Motte, Tom Beeckman, Steven Maere

## Abstract

Spatial omics technologies localize individual molecules at subcellular resolution, shedding light on the spatial micro-organisation of living organisms. Yet the development of analysis methods struggles to keep pace with growing numbers of molecules, features and replicates being measured, and with new scientific questions arising on single molecules’ localization patterns. To meet this need, we present *smoppix*, a nonparametric analysis method based on the probabilistic index, which unifies tests for several univariate and bivariate localization patterns, such as aggregation of transcripts or colocalization of transcript pairs, in a single framework. These tests can be performed across tissues as well as within cells, while accounting for nested design structures. The high-dimensionality of the data is exploited for variance weighting and for providing a meaningful background null distribution, unique for every individual molecule. *smoppix* sidesteps segmentation, warping, edge correction and density estimation, and scales to high numbers of molecules and replicates thanks to an exact permutation null distribution. We demonstrate its power by unearthing spatial patterns in four published datasets from different kingdoms, and validate some findings experimentally on *Selaginella moellendorfii* roots. Our method is available from Bioconductor as the R-package *smoppix*.

---

Advances in measurement technologies are driving a bonanza of spatial biology studies [1, 2, 3]. Biological tissues can be studied up to single-molecule resolution using multiple techniques. Sequencing technologies have been adapted to sequence separate libraries for a set of locations on the same tissue slice. Alternatively, individual RNA molecules and proteins can be tagged with fluorescent markers and localized; a technique called single-molecule localization microscopy (SMLM) [2, 4, 5]. In addition, cellular structures such as cell walls or nuclei can be stained and visualized on the same sections [2]. Finally, on a coarser scale, also different cell types can be localized in a tissue [1, 3]. Sequencing technologies are more scalable to whole-transcriptome profiling, whereas SMLM localizes a smaller set of pre-defined molecules with higher sensitivity and finer resolution [1]. As a result, SMLM is better at uncovering small-scale phenomena such as aggregation of transcripts within the cell or tissue, concentration gradients or co- or antilocalization of transcript pairs [3, 6]. Even though sequencing and SMLM study the same biological phenomena, the data types they generate are fundamentally different. Sequencing measurements consist of quantitative readouts (counts), whereas SMLM yields the locations of single molecules. Moreover, sequencing libraries are generated at a fixed grid of locations, whereas SMLM detects molecules at their natural, random positions. Because of these differences, custom analysis methods are needed for both data types. In this work we focus on the analysis of SMLM and single-cell localization data.

Spatial omics measurements are often replicated to account for technical and biological variability. This replication ranges from taking several images or fields of view (fov) from the same specimen, over cutting multiple sections of the same individual to measuring multiple individuals. A special form of replication arises when multiple cells are measured within the same image; if within-cell patterns are of interest, these cells serve as replicates. Given the complex makeup of biological tissues and irregular sectioning, rigorous statistical methods are needed to detect spatial patterns that persist across these replication levels. Apart from establishing if such patterns exist, researchers may also want to detect differences in localization patterns between conditions, e.g. study how a mutation or treatment affects the colocalization of a transcript pair. Although the human eye is good at detecting spatial patterns, visual inspection of images is labour-intensive and prone to cherry picking. Instead, many SMLM studies bin single molecules into artificial segments [4] or attempt cell segmentation [2, 5], and count the observed molecules within these segments to get quantitative readouts, reminiscent of sequencing datasets. These are then analysed using off-the-shelf statistical methods such as Pearson correlation. Yet this squanders the single-molecule resolution, and often the remaining location information of the segments is ignored in subsequent analyses [2, 4, 5]. Hence custom methods are needed to analyze collections of single-molecule locations, known in spatial statistics as *spatial point patterns*.

Historically, spatial statistics methods were developed for single point patterns and often revolve around testing the null hypothesis of complete spatial randomness (CSR), which implies that the molecules are distributed independently over the measurement area [7]. Biologically interesting departures from CSR, aggregation and regularity, as well as bivariate independence and its departures co- and antilocalization are shown in Figure 1a. In order to detect departures from CSR, a point pattern is commonly characterized through a summary function of intermolecule distances, calculated over a range of such distances. A popular choice is Ripley’s K-function, which is estimated for a single feature *g* as [7]

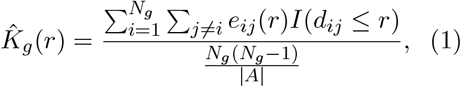

with *r >* 0 the range, *e*_*ij*_ an edge correction weight that accounts for censoring of points at the edge of the point pattern, *d*_*ij*_ the distance between molecules *i* and *j* with *i, j* = 1, …, *N*_*g*_, *N*_*g*_ the number of molecules of type *g* and *I*() the indicator function. 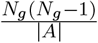 corrects for the baseline intensity of feature *g* in the point pattern on region A with area |A|. A bivariate equivalent of the K-function can be used to test for co- and antilocalization of transcript pairs [7]. Under CSR, *K*_0_(*r*) = *πr*^2^. To assess significance of the departure of 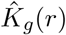 from *K*_0_(r), Monte-Carlo simulations are used to quantify the variability of 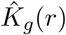 under CSR, e.g. through simulation envelopes [7] (see Figure 1b). Yet this approach is not scalable to spatial omics datasets with many features, and not generalisable to replicated experiments. A more automated approach is the Diggle-Cressie-Loosmore-Ford (DCLF) test with test statistic [7, 8]

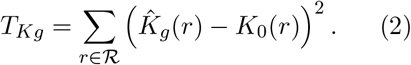

**Figure 1:**
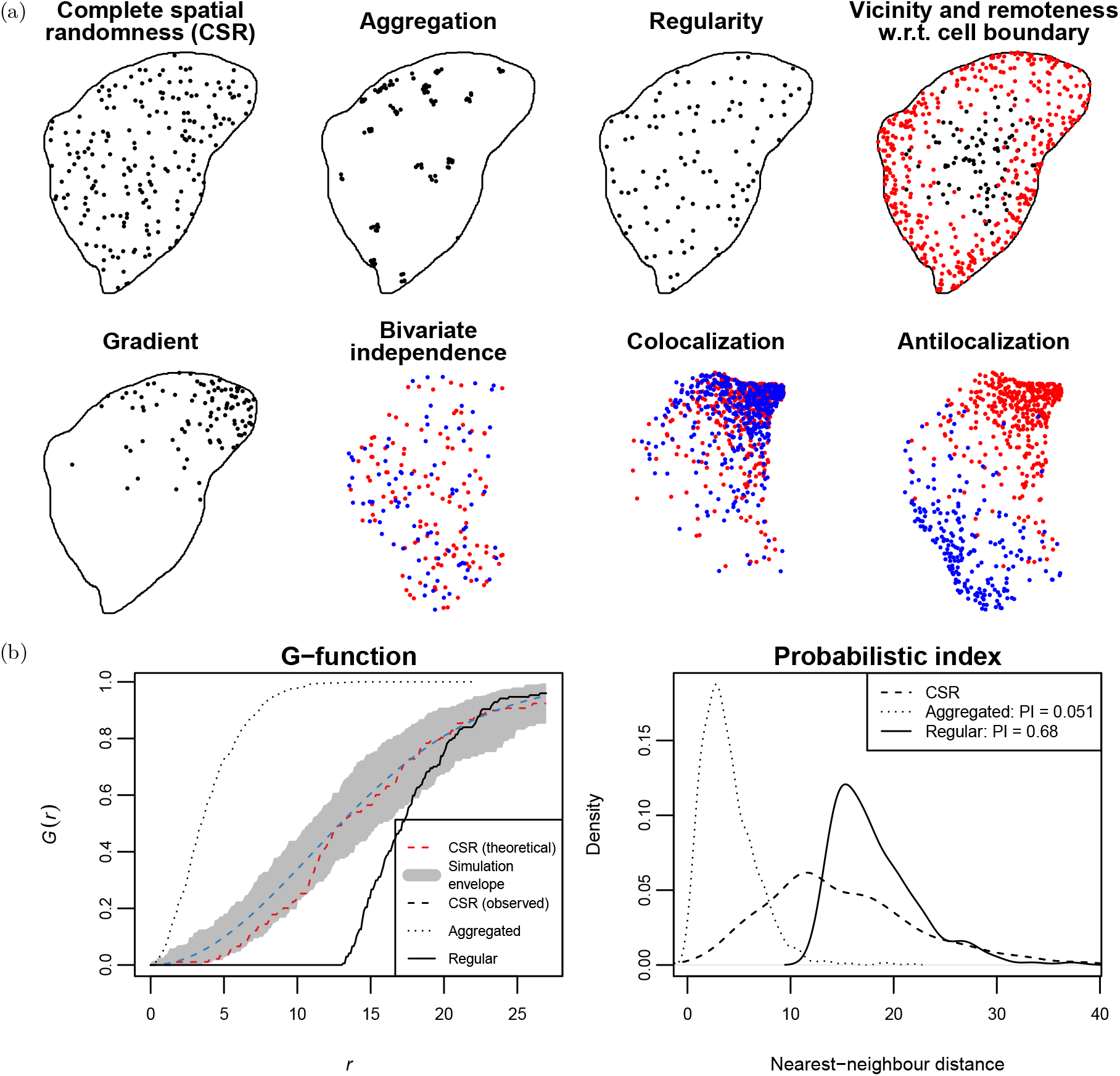
(a) Top row and left plot on second row: univariate CSR in a cell with different types of departure. The solid line indicates the cell boundary, the dots represent individual transcripts. In the top right plot, red and black dots represent two molecules close to and far from the cell boundary, respectively. The rightmost three plots on the second row depict bivariate independence and dependence types. Red and blue dots indicate different molecules. (b) Left: G-function (the nearest-neighbour distance distribution function, y-axis) as a function of range *r* (x-axis) for the first three point patterns in the top row (in black). The dashed red line is the theoretical G-function under CSR, the grey area represents a 95% simulation envelope. Right: Nearest-neighbour distance densities for the same three point patterns. The corresponding probabilistic index (PI) estimate with respect to CSR is shown in the legend.

*T*_*Kg*_ summarizes the departure from *K*_0_(*r*) over a range of distances ℛ in a single number; its null distribution is again approximated by Monte-Carlo simulation. For Monte-Carlo testing, it has been suggested to convert the K-function to the L-function 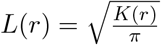 [9]. However, because of the square in the calculation of *T*_*Kg*_, information on the direction of the departure is lost, making it poorly suited to combine evidence across replicated point patterns. An alternative summary function is the cumulative distribution function of nearest-neighbour distances, the G-function, which is estimated as [7]

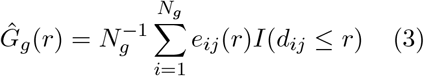

with *d*_*ij*_ the distance of point *i* to the nearest other molecule *j* of type *g*.

Analysis of replicated point patterns has been tackled by modelling the K- or G-function over different replicates in a parametric [10], nonparametric [11, 12, 13, 14] or functional [15] way. Many other methods use some scalar summary of such a function to simplify the analysis. Canete et al. [16] based their *spicyR* method to test for colocalization of cell types on an integral over the difference between the L-function and *r*. The resulting u-statistic has the desirable property of having expectation 0 under the null hypothesis, and is used as outcome in a linear model that accounts for the replication structure of the experiment. Similarly, Barua et al. [17] employ the area-under-the-curve (AUC) of the G-function as summary measure in subsequent analysis. The *DIMPLE* method estimates univariate molecule densities as a function of space and compares them across features using Jensen-Shannon distances, which are used as extracted features in follow-up analyses [18]. All these methods rely on feature-wise density estimation to account for density differences across the point pattern, for instance by plugging a location-dependent density estimate into the denominator of (1). Yet density estimation is difficult for ragged expression patterns, which occur naturally in tissues or result from tearing or degradation of tissue sections [19], or within cells where the number of molecules is low, resulting in low power and high computational demands [8, 19, 20]. Also edge correction can be problematic when no clear delineation of the area A is provided. The algorithm by Bauer et al. [21] quantifies the probability of the observed nearest-neighbour distance occurring by chance through permutations, somewhat similar to the method we propose below, although it relies on hard cut-offs for both the probability and the observed nearest-neighbour distance. Neighbourhood enrichment methods have also been proposed for the analysis of SMLM data [22, 23, 24], but they do not allow to analyze replicated point patterns.

To address these shortcomings, we present *smoppix* (for Single-MOlecule sPatial omics data analysed through the Probabilistic IndeX): a versatile nonparametric framework for hypothesis testing on replicated SMLM or cell type location data based on probabilistic indices (PIs). We apply *smoppix* to four datasets from different kingdoms to unearth previously undetected spatial patterns, and validate some of our findings experimentally. Our method is available from Bioconductor as the R-package *smoppix*.

## Results

### The probabilistic index summarizes differences between distance distributions

Our first objective is detecting vicinity or remoteness of molecules from fixed cellular structures, such as cell boundaries (see Figure 1a). Joyner et al. [25] introduced the B-function as the cumulative distribution function of distances of spiders to the nearest edge of their web. For spatial omics data, the B-function of feature *g* can be estimated as:

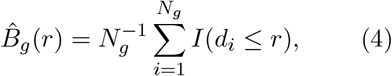

with *d*_*i*_ the distance of molecule *i* to e.g. the closest cell boundary. Joyner et al. [25] employ simulation envelopes for significance testing. Yet one can reduce computation time and gain power by observing that the B-function only considers one molecule at the time, such that the terms in (4) are independent. This observation unlocks classical goodness-of-fit tests to compare the observed and null distributions, which are invalid for intermolecule distances that have complex dependencies. Unlike for the DCLF test, the p-values of such tests are asymptotic and no longer discretized by the number of Monte-Carlo instances. Monte-Carlo simulation is still needed to approximate the B-function under the null hypothesis of CSR, 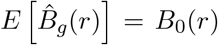, which is identical for all features (transcript types), but no longer to quantify the variability of 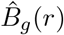. The goodness-of-fit tests most in line with (2) are the Anderson-Darling [26] and Cramér-von Mises [27, 28] tests, with test statistics of the form:

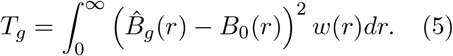

Setting *w*(*r*) = 1 yields the Cramér-von Mises test, while *w*(*r*) = [*B*_0_(*r*)(1 − *B*_0_(*r*))]^−1^ yields the Anderson-Darling test. No more arbitrary choice of integration limits as in (2) is needed, but the quadratic form of *T*_*g*_ still necessitates manual inspection of directionality for significant departures from the null.

As a remedy, our method *smoppix* employs the probabilistic index (PI) to contrast the null with the observed distribution. The PI is the probability that a randomly drawn observation from a population *k* is smaller than an observation from another population *l*, plus one half the probability that they are equal [29]:

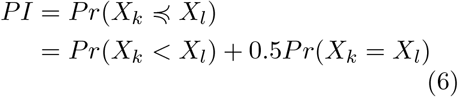

The PI is known in predictive modelling as the area-under-the-curve (AUC), and is the central quantity in the Wilcoxon-Mann-Whitney test [29]. We choose as population *k* the one under the null hypothesis, and population *l* as the observed one, such that the PI estimate of an observed distance *d*_*i*_ equals its evaluation in *B*_0_: *PI*_*i*_ = *B*_0_(*d*_*i*_). In analogy to (5), the PI can then be written as

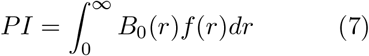

with *f* the density of the observed distances [29]. Values larger than 0.5 indicate remoteness from the cell boundary, whereas values smaller than 0.5 indicate proximity.

### Nearest-neighbour distances capture univariate and bivariate patterns

Next we aim to capture univariate and bivariate localization patterns. Nearest-neighbour distances are known to capture such patterns better than the collection of all intermolecule distances [7], so for this purpose we replace the B-function by the uni- or bivariate G-function in (7). PIs smaller than 0.5 indicate aggregation or colocalization, estimates larger than 0.5 indicate regularity or antilocalization (see Figure 1). Yet a complication arises, as nearest-neighbour distances are not independent, but rather have a complex dependence structure. As a solution, the *PI*_*i*_ estimates are averaged per feature *g* over all molecules *i* of a point pattern *t* to yield a single 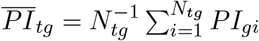 with *t* = 1, …, *n* indicating the point pattern (replicate) and *N*_*tg*_ the number of molecules of feature *g* in point pattern *t*. For the bivariate PIs, we average over both G-functions, i.e. for features *g* and *h*

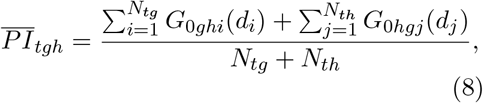

with *G*_0*ghi*_ the null distribution of nearest neighbour-distances of molecules of feature *h* to molecule *i* of feature *g* (see Methods for the rationale of the index *i*). These *PI*_*t*_ estimates are independent and can be used in downstream analyses. Aggregation patterns can be tested across tissues or within cells; in the latter case each cell is considered as an independent point pattern (replicate), and only distances within cells are taken into account.

### The choice between two null hypotheses

We distinguish two null distance distributions for the calculation of the PI. So far we have discussed CSR within the area A as null model, and have approximated the corresponding distance distributions *B*_0_ and *G*_0_ by Monte-Carlo simulation. Especially for intracellular patterns this may be the most useful null hypothesis to test, as the cells provide a natural choice for the area A. However, in many datasets, the molecules are inhomogeneously distributed across the section or tissue due to differences in cell density or metabolic activity or due to sample degradation [19], or no area is given at all [4, 5], making CSR an irrelevant null hypothesis. In such case we choose to exploit the high-dimensional nature of the data and let the collection of molecules of all features provide the null distance distribution. For fixed objects such as the cell boundary, *B*_0_ is then simply the observed distribution function of distances of all features’ molecules to the object. Nearest-neighbour distances are density-dependent, so to find their null distribution we could execute feature label permutations, similar to e.g. Bauer et al. [21] and Edsgärd et al. [35]. Yet this is computationally prohibitive for spatial omics datasets, and instead we propose an exact permutation null distribution based on the negative hypergeometric distribution (see Methods). Using the background null distribution rather than CSR changes the interpretation of the test. One is no longer assessing whether the molecules are aggregated or colocalized, but whether they are more aggregated or colocalized than the collection of all molecules, which is biologically often more meaningful.

### The PI as outcome in linear mixed models

So far, we have calculated the PI per molecule *i* (for the B-function) or point pattern *t* (for the G-function), but we want to analyze all point patterns jointly while accounting for the nested design structure. For this purpose, we propose a linear mixed model with the PI as outcome. For instance, for the B-function the model becomes

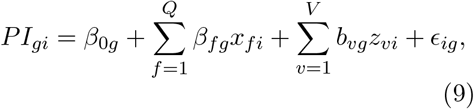

including *Q* random effects for cells, fields of view or individual organisms, and *V* fixed effects for covariates of interest such as treatments, and 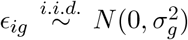. For PIs derived from the G-function, 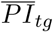 or 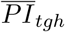 is used as outcome, replacing index *i* by *t*, and no random effects for cell or image are included. Significance tests on *β*_0*g*_ and *β*_*fg*_ than reveal baseline departures from the null distribution and differences between treatment groups, respectively (see Methods). Note that the permutation null distribution introduced above is used to estimate the PI but not for significance testing: asymptotic p-values per feature are derived from model (9), which quantifies variability over replicates. Point patterns with many molecules yield more precise estimates of the nearest-neighbour PI (8), even though the constituting entries are not independent. Hence we attach observation weights to the different PIs in the regression model (9), inspired on *spicyR* [16], but with some improvements (see Methods).

### *smoppix* provides calibrated tests and outperforms competitors computationally

To probe calibration of the statistical tests, we permuted the gene labels of two spatial transcriptomics datasets [4, 5] 100 times, and applied *smoppix* with background null and competitor methods *spicyR* and neighbourhood enrichment (NE, see Methods). *smoppix* yields mostly uniform p-value distributions, whereas *spicyR* and NE were found to have inflated type I error (Supplementary Figures S35-S Moreover, the PI is mostly invariant under subsampling of molecules (see Supplementary Section 2.2), indicating that it is insensitive to density changes, as is the NE log-ratio. *spicyR*’s u-statistic and the univariate nearest-neighbour PI, on the other hand, are sensitive to the molecule density, although the changes under subsampling are minor compared to overall variability of the test statistics. A computational benchmark (Supplementary Section 2.3) revealed that *smoppix*’ RAM memory usage and computation time scale well with increasing numbers of molecules and features. Most of its computing time is consumed by fitting the mixed models that account for the replicated design structure, rather than estimating the PIs. *spicyR* on the other hand, takes longer to compute and its memory usage increases quickly with the number of molecules. NE becomes very slow even for a moderate number of features and molecules.

### Real data analysis

We applied *smoppix* to four public datasets from different kingdoms. P-values were Benjamini-Hochberg adjusted [30] to control the false discovery rate at 0.05 over all features or feature pairs per type of PI. Where applicable, we also apply the alternative methods *spicyR* and NE.

### *smoppix* ‘ discoveries on *Selaginella moellen-dorfii* roots are confirmed by HCR RNA-FISH

Yang et al. [4] imaged transcripts of 97 genes through single-molecule fluorescent in situ hybridization (smFISH) in *Selaginella moellendorfii* roots after 0 and 3 days of growth after the first root bifurcation. Six plants were measured at each timepoint, with a variable number of sections per root. The original analysis binned the images into hexagons, calculated Pearson correlations between the resulting counts and clustered the genes hierarchically based on these correlations. Since most genes are mainly expressed in the central, vascular tissue of the roots (see Supplementary Figure S1), a *smoppix* analysis using CSR within the whole root as null model would yield trivial results, declaring transcripts of every gene aggregated and transcripts of every gene pair colocalized. Instead, we use the background null distribution, which automatically accounts for the differences in shape and transcript density between the roots without requiring density estimation. Our results on colocalization often differ from the ones in the original analysis and from those of *spicyR* (see Figure 2, Supplementary Table S1 and Figure S7). *smoppix*’ results overlap better with those of NE, although the latter makes fewer discoveries (Supplementary Section 1.1.2).

**Figure 2:**
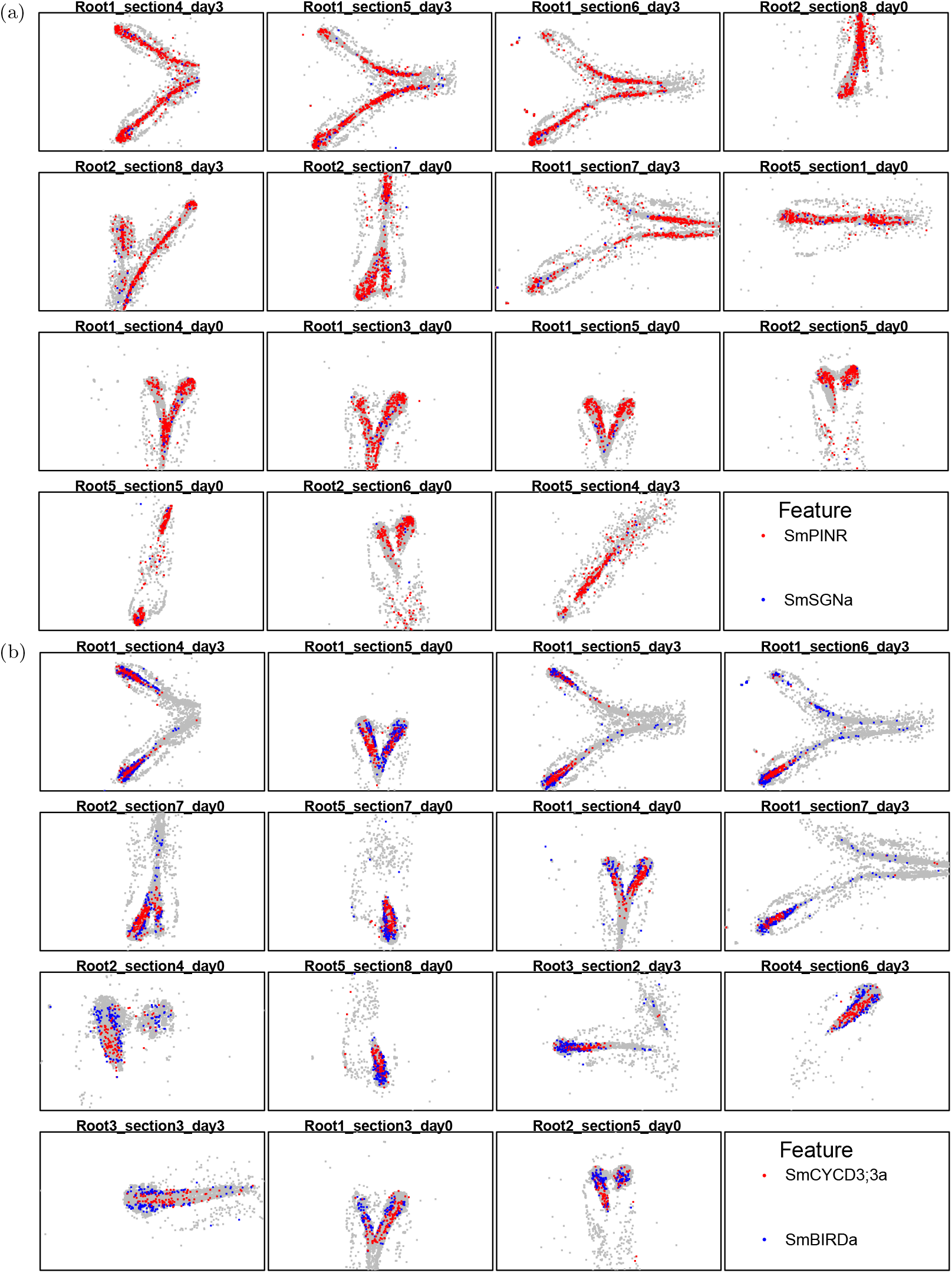
Gene pairs found colocalized (a) and antilocalized (b) in expression by *smoppix* in *S. moellendorfii* root dataset, but not in the original analysis [4]. The red and blue dots are the transcripts of the gene pairs, all other transcripts are shown in grey. The subset of 11 sections with highest expression of the gene pair per root and day is shown, as indicated in the plot titles.

To validate *smoppix*’ findings, three transcript pairs were visualized in *S. moellendor-fii* roots using hybridization chain reaction (HCR) RNA-FISH [31]. These pairs were selected for being significantly co- or antilocalized according to *smoppix*, and not significant or having an opposite pattern according to *spicyR*, NE and/or in the original analysis (see Supplementary Table S2). Multiple roots were measured for over a range of z-values. Example images are shown in Figure 3a, all images have been deposited to BioStudies with accession number S-BIAD1942. Colocalization was quantified through the pixel-by-pixel Pearson correlation of the intensities of the different dyes (see Supplementary Section 1.1.3). The results of the hypothesis test of the true correlation being zero are shown in Table 1 (top). This analysis confirms antilocalization of the SmBIRDa-SmCYCD3;3a pair, as picked up by *smoppix* only. It also calls antilocalization for the SmSGNa-SmPINR pair, contradicting the results from *smoppix, spicyR* and the original analysis. Yet the spatial distribution of the *SmSGNa* transcript looks very different in the two assays, being lowly expressed and aggregated in the smFISH data (Figure 2a) but expressed across the entire root in the HCR RNA-FISH data (Figure 3a), which may explain the discrepancy. The SmSGNb-SmPINS pair was found colocalized by *smoppix* and *spicyR*, but in the HCR RNA-FISH assay it was found negatively correlated with borderline significance. On visual inspection, both genes are solely expressed in the central vascular tissue, although their intensity measurements do not overlap exactly, leading to negative pixel-by-pixel correlations (see Figure 3a). This discrepancy in results may simply reflect the better resolution of the HCR RNA-FISH technology. Transcript pairs found antilocalized at coarse resolution by smFISH will still be found antilocalized at a finer resolution by HCR RNA-FISH (e.g. the SmBIRDa-SmCYCD3;3a pair), but transcript pairs found colocalized by smFISH may be revealed to be locally antilocalized by HCR RNA-FISH (e.g. the SmSGNa-SmPINR pair). To confirm this hypothesis, a second HCR RNA-FISH experiment was set up, including five additional transcript pairs that were declared antilocalized by *smoppix* based on the sm-FISH results, but were found colocalized in the original analysis. In addition, the three transcript pairs already tested were included with a dye swap. The results are shown in Figure 3b and Table 1 (bottom), and confirm the results from the first experiment for the three repeated transcript pairs. The other five transcript pairs were all found significantly antilocalized by HCR RNA-FISH. This demonstrates how *smoppix* maximally exploits the resolution of the smFISH data to detect antilocalization, where methods based on binning and *spicyR* falsely con-clude colocalization.

**Table 1:** Results for pixelwise correlation test between transcript pairs in two HCR RNA-FISH experiments, obtained from mixed models similar to (9) (see Supplementary Section 1.1.3). The dashed line separates repeated from new transcript pairs in the second experiment.

|  | Experiment | Estimate | Standard error | P-value | Adjusted p-value |
| --- | --- | --- | --- | --- | --- |
| SmCYCD3;3a–SmBIRDa | 1 | -0.430 | 0.045 | 4.5e-05 | 1.3e-04 |
| SmPINRa–SmSGNa | 1 | -0.308 | 0.029 | 1.5e-03 | 2.2e-03 |
| SmPINS–SmSGNb | 1 | -0.052 | 0.014 | 3.3e-02 | 3.3e-02 |
| SmCYCD3;3a–SmBIRDa | 2 | -0.368 | 0.012 | 9.4e-08 | 3.7e-07 |
| SmPINR–SmSGNa | 2 | -0.478 | 0.022 | 5.8e-07 | 9.3e-07 |
| SmPINS–SmSGNb | 2 | -0.030 | 0.034 | 4.0e-01 | 4.0e-01 |
| SmRBRa–SmSCRa | 2 | -0.545 | 0.026 | 1.5e-07 | 4.0e-07 |
| SmRBRa–SmSCRb | 2 | -0.409 | 0.028 | 6.9e-06 | 7.9e-06 |
| SmBIRDa–SmSCRb | 2 | -0.479 | 0.021 | 4.1e-07 | 8.2e-07 |
| SmSHRa–SmSCRb | 2 | -0.318 | 0.006 | 3.7e-09 | 2.9e-08 |
| SmCYCD3;3a–SmSHRa | 2 | -0.338 | 0.026 | 3.9e-06 | 5.2e-06 |

**Figure 3:**
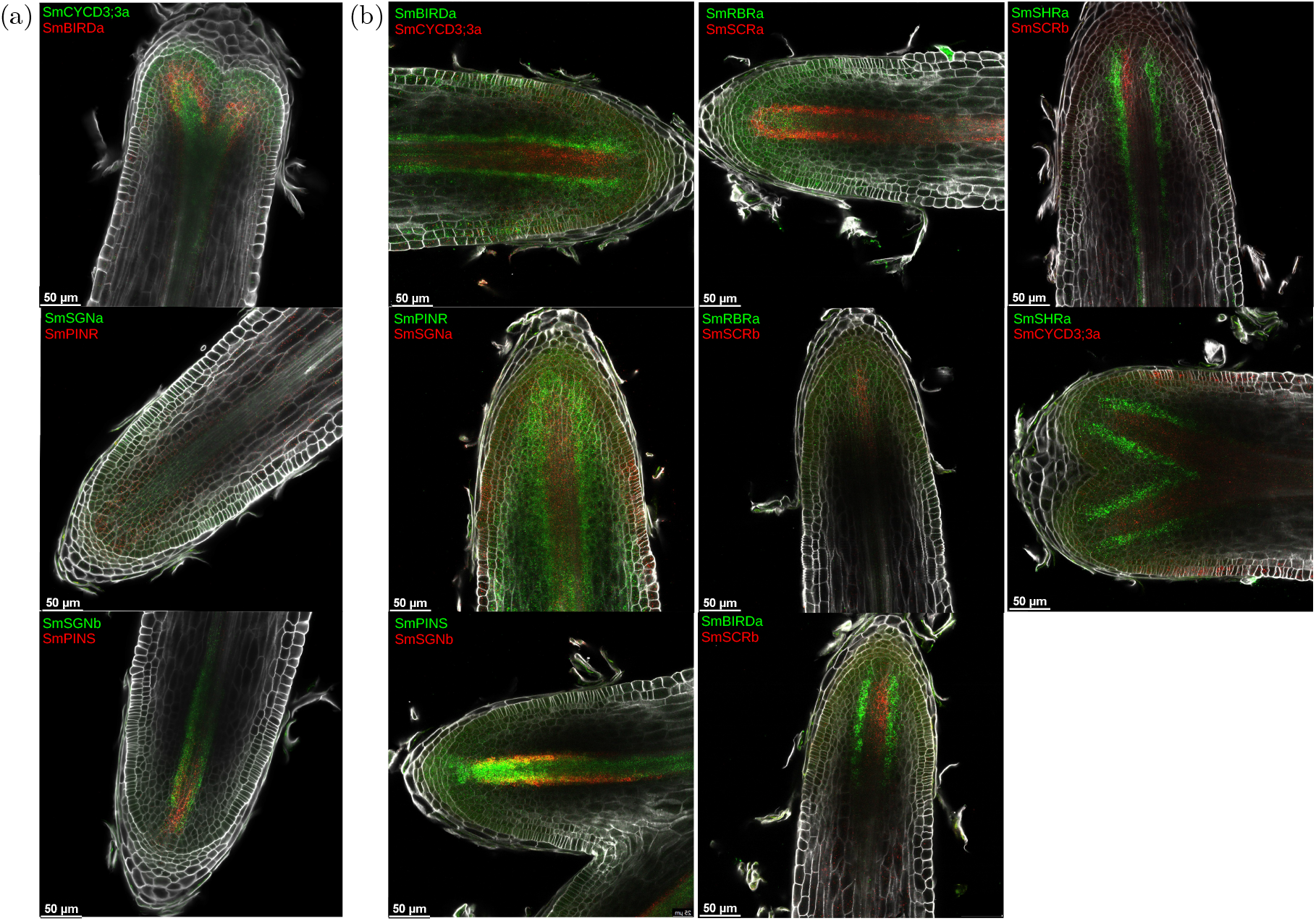
Selected images from a central z-coordinate for the eight *S. moellendorfii* gene pairs investigated through HCR RNA-FISH in the first experiment (a) and the second experiment (b). Transcripts are shown in green and red for the genes as indicated on the images, cell walls are shown in white.

### *smoppix* finds intracellular localization patterns in mouse fibroblast cells

Eng et al. [2] profiled 10,000 genes in mouse fibroblast cells through sequential FISH (seqFISH), with cell boundaries gated by hand based on DAPI staining. *smoppix* detects transcripts close to and far from the cell edge compared to CSR (Figure 4), as well as transcripts aggregated within the cell. No other analysis method allows to test for vicinity to fixed objects; testing for aggregation within the cell was found to be too time-consuming for NE and too memory-intensive for *spicyR. smoppix*’ results correspond to some extent to the authors’ original analysis based on clustering (see Supplementary Section 1.2), yet it makes many more discoveries and assigns significance to them. Moreover, *smoppix* is able to detect transcripts with unique spatial patterns that are not part of a cluster of spatially correlated genes.

**Figure 4:**
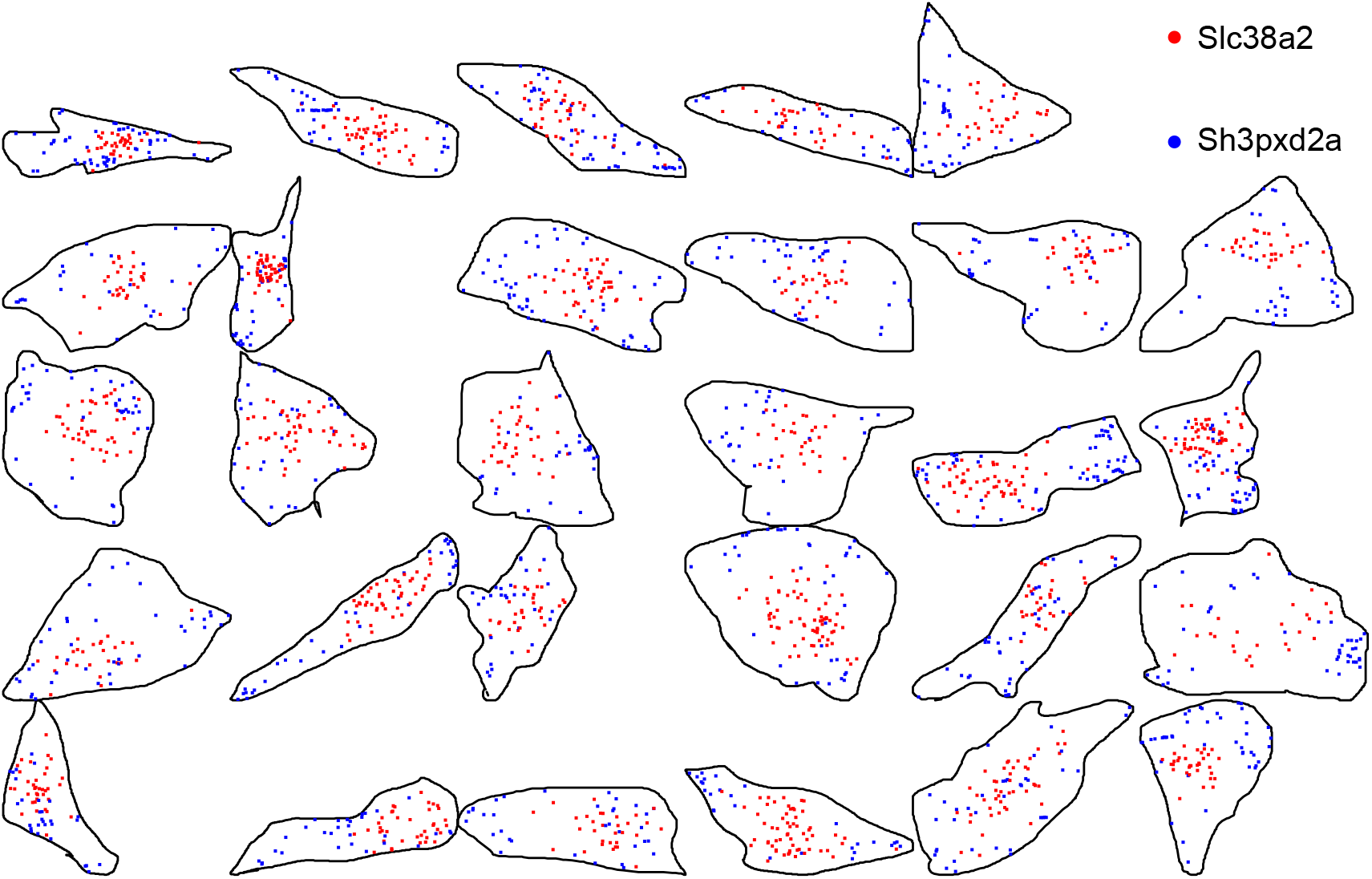
Transcripts most significantly close to (Sh3pxd2a) and far from (Slc38a2) the cell boundary according to *smoppix* in the mouse fibroblast dataset by Eng et al. [2]. A subset of 29 cells with highest expression of these two genes over all fields of view (fovs) is shown, not in their original position in the fov.

### *smoppix* detects shifts in localization patterns over time in bacterial biofilms

Dar et al. [5] profiled *Pseudomonas aeruginosa* biofilms after 10h and 35h of growth for spatial expression of 108 genes using seqFISH. A more pronounced spatial concentration of molecules after 35h is visible by the naked eye. The original analysis used Pearson correlation within segmented cells and visual inspection to look for spatial localization patterns and differences between timepoints. *smoppix* finds recurrent spatial patterns and differences in spatial patterns between 10h and 35h of growth, conditional on the overall differences in density. Figure 5 shows the transcript pair most significantly more colocalized after 35h of growth than after 10h, which was not discovered in the original analysis. More significant results can be found in Supplementary Section 1.3, demonstrating *smoppix*’ ability to detect differences in spatial organisation between conditions.

**Figure 5:**
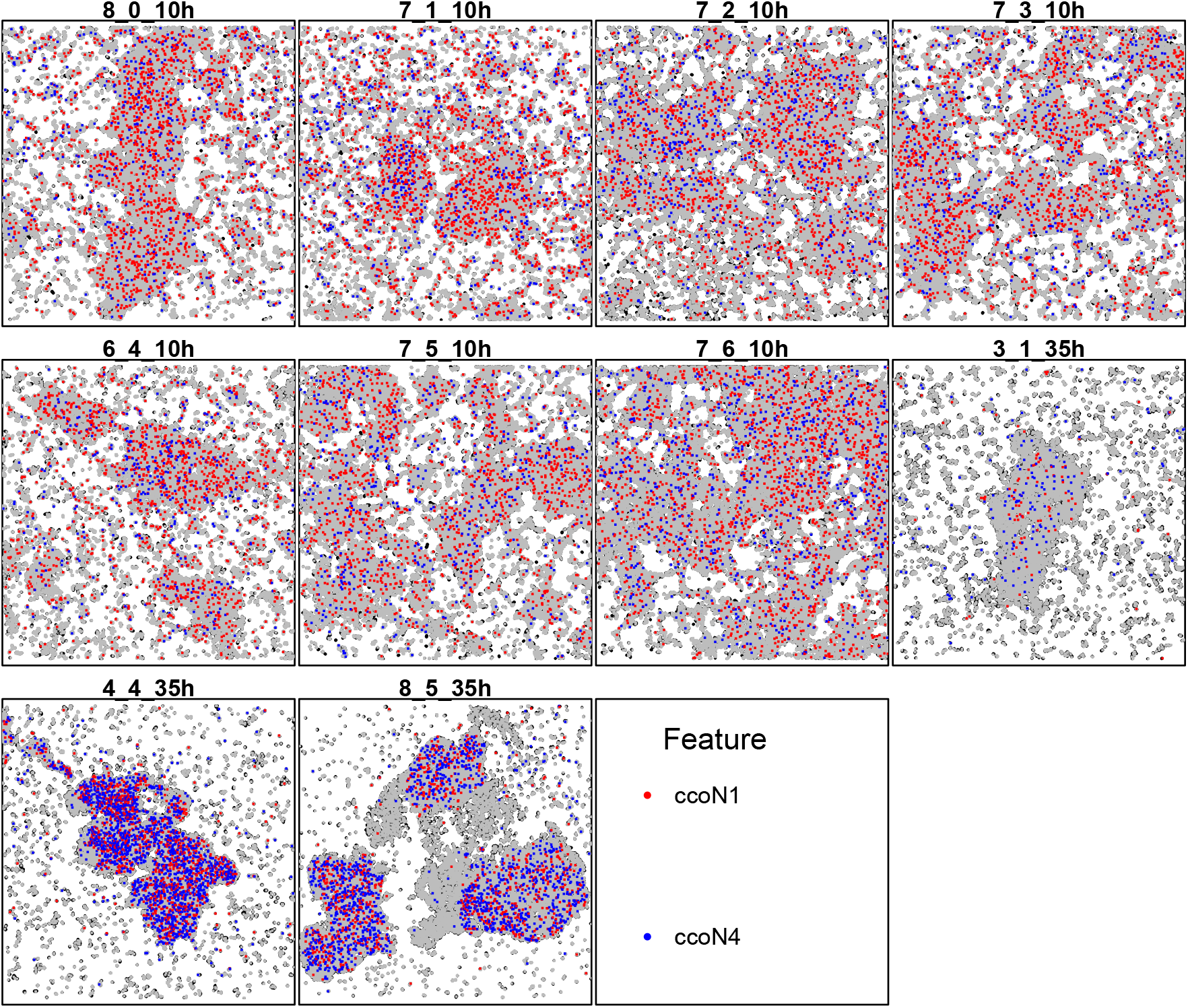
Transcript pair most significantly more colocalized after 35h than after 10h of growth according to *smoppix* in the bacterial biofilm dataset by Dar et al. [5]. The red and blue dots are the transcripts pair, all other transcripts are shown in grey. The titles show z-coordinate, replicate and timepoint, in that order. Per biofilm, the z-plane with highest overall expression is shown.

### Probabilistic indices serve as interpretable predictors in tumor classification

Keren et al. [32] localized different cell types in breast tumor sections, using the adjacency patterns of the tumor cells to classify the tumor type into either ‘compartimentalised’ or ‘mixed’. First we use *smoppix* with background null distribution to test for association between PIs and tumor type, conditional on patient age. Among univariate aggregation patterns, only that of tumor cells is found to be different between the tumor types (see Figure 6). This agrees with the tumor type definition, and corresponds to the results of NE. Remarkably, *spicyR* did not detect any differentially localized cell types, not even the tumor cells (see Supplementary Figure S34). Presumably, the u-statistic is affected by the difference in overall cell count between the two tumor types (Supplementary Figure S31) or by the empty spaces in the sections.

**Figure 6:**
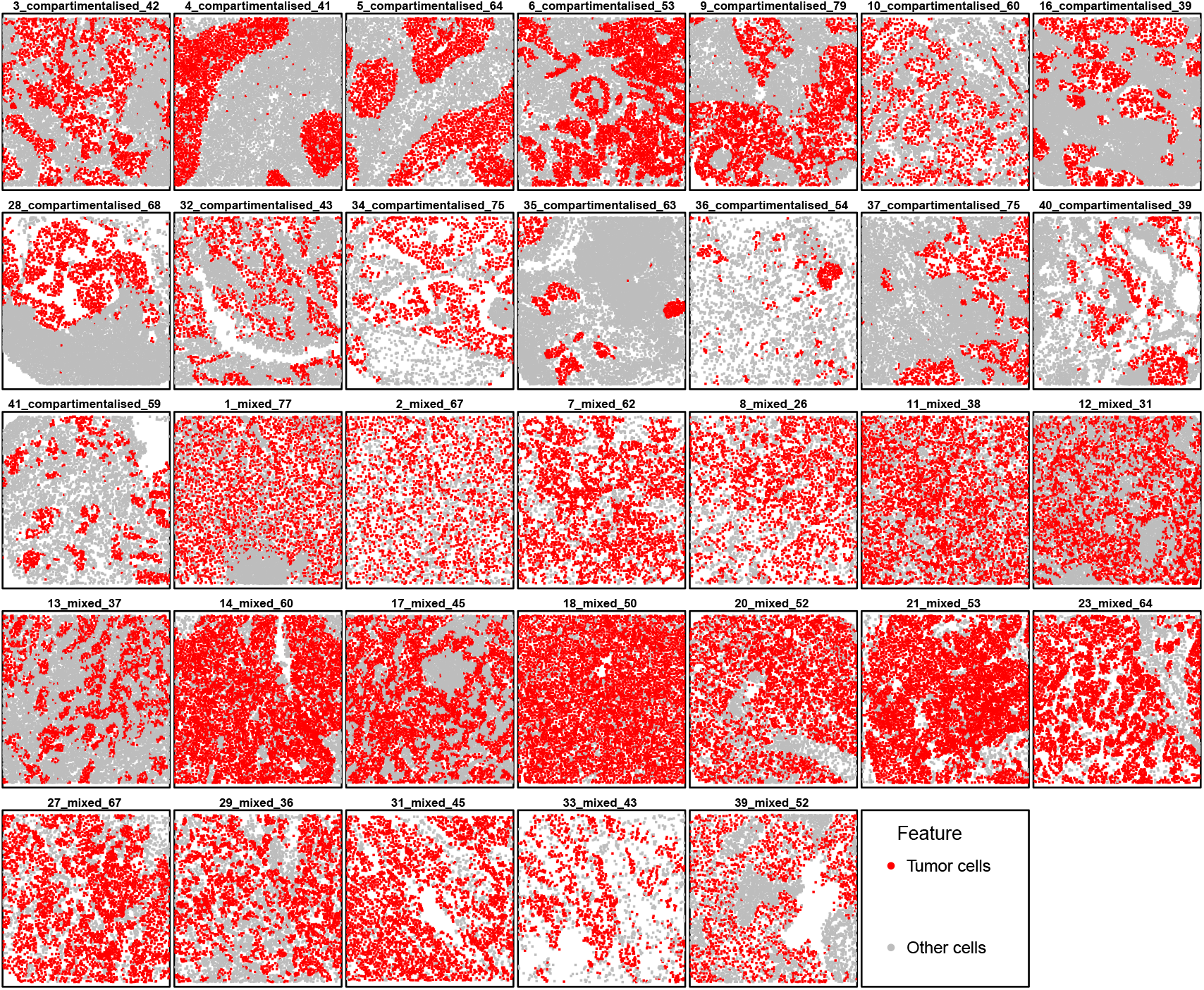
Breast cancer data by Keren et al. [32], with tumor cells shown in red and the other cell types in grey. The plot titles indicate patient id, tumor type and age, in that order.

Apart from hypothesis testing, PIs can serve as interpretable features for other statistical purposes too. Here we investigate their use as predictors in a regularized generalized linear model (LASSO [33]) to predict tumor type, see Supplementary Section 1.4.2 for details. Estimated accuracies in Table 2 suggest that the inclusion of the PIs as predictors improves predictive accuracy. Using the NE log-ratios as predictors yields a similar result. The *spicyR* u-statistic did not yield any improvement when included as predictor (see Supplementary Section 1.4.2).

**Table 2:** Estimated predictive accuracy and its standard error (SE) for LASSO regression models predicting tumor type on breast cancer data by Keren et al. [32] using different sets of predictors. The cell type count is the number of times each cell type was found in the section.

| Predictors | Accuracy | SE |
| --- | --- | --- |
| Age | 0.56 | 0.09 |
| Age, cell type count | 0.84 | 0.12 |
| Age, PI | 0.97 | 0.07 |
| Age, cell type count, PI | 0.91 | 0.05 |

## Discussion

Historically, analysis methods for spatial point patterns served to guide manual inspection of single, low-dimensional patterns, e.g. Ripley’s K-function. They reveal at which scale spatial patterns occur, but are time-consuming to interpret. Current summary measures of such functions are unadapted to the research questions and replicated nature of single-molecule spatial omics experiments, and computationally too demanding.

As a remedy, we present our *smoppix* method based on the probabilistic index (PI). The PI is a simple, interpretable measure, well-embedded in statistical theory. It detects unusual distance distributions with respect to fixed objects such as cell boundaries, as well as to the nearest other molecules to discover univariate and bivariate (co)localization patterns. It is a probability and thus unitless, rendering it comparable across replicated point patterns of varying densities and shapes. Calculating the PI does not require integrating a summary function, (pseudo)segmentation, density estimation, warping or edge correction. A custom C++ algorithm and an exact permutation null distribution based on the negative hypergeometric distribution ensure quick estimation of the PI of nearest-neighbour distances. Focussing on nearest-neighbour distances automatically adapts to the resolution of the measurement technique, unearthing patterns at the smallest scale measured. Crucially, *smoppix* not only detects spatial patterns but also allows for formal comparison of these patterns across conditions.

We demonstrated how *smoppix* picks up spatial patterns more accurately than other analysis methods through confirmatory experiments on *Selaginella moellendorfii* roots using HCR RNA-FISH. Outside hypothesis testing, PIs are interpretable features extracted from point patterns that can serve diverse statistical purposes, e.g. the prediction of tumor types from cell type localization data. The PI allows for easy extension to other applications that use distances, e.g. the colocalization of more than two molecules can be quantified through the distribution of the radius of the smallest circle or sphere encompassing them.

Clustering approaches are often used as a rough breakdown of the spatial organisation of molecules [2, 4, 34]. Yet they have the drawback of assigning all features to some cluster, even those that show no localization pattern at all. In addition, uncertainty on cluster membership or number of clusters is usually not quantified, leaving questions on reproducibility open. On the contrary, *smoppix* presents a gene-by-gene approach that can also discover features with unique spatial patterns, and provides measures of uncertainty for all such discoveries.

Neighbourhood enrichment (NE) presents an interesting alternative, fixing a neighbourhood size rather than a number of nearest neighbours. We suggest a way to adapt NE tests to replicated point patterns, which worked well for cell-type data. Still, for spatial omics data with many more molecules and features, NE is not scalable. NE methods are asymmetric by convention in that they yield different results for enrichment of transcripts of gene *a* close to transcripts of gene *b* than vice versa. On the other hand, summary functions of point patterns, and also *smoppix*, usually yield symmetric results, i.e. report a single result per gene pair. Yet both methods are amenable to either symmetric or asymmetric analysis. Asymmetric results can be interesting biologically, but they also increase the burden of multiple testing and interpretation in a setting where the number of bivariate combinations is already very high.

A main drawback of *smoppix* is that it remains a two-step procedure, like all existing methods that account for replication to our knowledge. First a summary measure (the PI) is calculated for every point pattern. Next, this summary measure is plugged into a linear model that accounts for the design factors and quantifies variability. Yet a one-step model, e.g. a probabilistic index model (PIM) that models all pairwise distance comparisons (pseudo-observations) [29] while accounting for a complicated design structure and dependencies between nearest-neighbour distances, may be programmatically unfeasible. Although it may be inefficient, the two-step procedure allows for easy combination with existing software implementations and has proven its worth in practice. Another liability are the increasing computation times as the number of features grows, especially for bivariate patterns due to combinatorial explosion. Still *smoppix* is much more scalable than competitor methods.

## Supporting information

Data used in the analysis from already existing publications

## Author contributions

S.H. conceived and developed the *smoppix* method, S.H. and S.M. wrote the manuscript. X.Y., H.M., T.B., S.H. and S.M. conceived the HCR RNA-FISH experiment, X.Y. and W.P. conducted it. S.M. supervised the study.

## Competing interests

No competing interest is declared.

## Data and code availability

The generated HCR RNA-FISH data was deposited to the BioStudies repository with accession number S-BIAD1942. The code for reproducing the results in the paper is available at https://github.com/maerelab/SmoppixPaper and the data can be found in the Supplementary material.

## Funding

The work of S.H. in the lab of S.M. was supported by Inari Agriculture NV, funded in part by VLAIO, grant HBC.2019.2814.

## Methods

### An exact permutation null distribution for fast calculation of the PI

We propose a fast, exact way to calculate the nearest-neighbour distance PI with background null distribution for a point *i* of feature *g*. We take univariate nearest-neighbour distances as an example, dropping point pattern subscripts *t*. Consider the set of all distances from molecules of all types to molecule *i* of type *g*, with size *N* − 1 (excluding only the molecule *i* itself) and ordered such that *d*_(1)_ ≤ *d*_(2)_ ≤ … ≤ *d*_(*N* − 1)_. A classical permutation procedure would repeatedly sample *N*_*g*_ − 1 molecules without replacement from this set (*N*_*g*_ is the number of molecules of type *g*), and record the shortest one. This collection of shortest distances then builds a unique null distribution *G*_0*i*_, which differs depending on whether *i* is located at the center or at the edge of the point cloud. *G*_0*i*_(*d*_*i*_) serves as PI estimate for molecule *i*, with *d*_*i*_ the observed nearest-neighbour distance. Conditioning on the observed point cloud eliminates the need for edge correction, density estimation or area delineation. Yet repeated permutations for every point separately are computationally prohibitive for realistically sized spatial omics datasets, especially since the complete distance matrix of size *N* (*N* − 1)*/*2 usually cannot be stored in working memory. As a solution, we observe that feature label permutation can be seen as sampling without replacement from an urn with *N*_*g*_ − 1 successes and *N* − *N*_*g*_ failures. A success is defined as the sampled distance being assigned to feature *g*. For any distance *d*_(*x*)_ with rank *x* in the sorted list, the probability of success equals 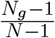. Yet for the nearest-neighbour distance PI, we need the probability that a distance *d*_(*x*)_ is assigned to feature *g and* that it is the shortest among all distances assigned to *g*. This probability decreases with the rank *x* and can be found with the help of the negative hypergeometric distribution, which models the number of attempts *x* needed to reach *q* successes when sampling without replacement from an urn [36]. Setting *q* = 1, this distribution assigns to each distance *d*_(*x*)_ a probability of being the smallest among the set of *N*_*g*_ − 1 selected distances. This yields an exact expression for *G*_0*i*_, to which a Monte-Carlo permutation procedure would only provide a time-consuming approximation (see Supplementary section 3.1 for details).

### The PI as outcome in mixed linear models

To maintain interpretability of the intercept *β*_0*g*_ in (9) as overall mean, sum coding is used for the categorical fixed effects. This means that a dummy variable is created for every level of the variable, and the coefficients are constrained to sum to zero, unlike the more common choice of having one reference level for which no dummy is included. Analogously, continuous variables are mean-centered; random effects have mean zero by definition. The null hypothesis *H*_0_ : *β*_0*g*_ = 0.5 then tests for overall departure from the null distribution, while the null hypothesis *H*_0_ : *β*_*fg*_ = 0 tests for differences in distance distribution associated to fixed effect *f*. Unlike e.g. an average distance, the PI is unitless and hence comparable across point patterns with different sizes and shapes and corresponding distance distributions under the null. The normal assumption for the error *ϵ*_*ig*_ or *ϵ*_*tg*_ may seem in contradiction with the PI being bounded between 0 and 1. Still, in practice most observed *PI*_*ig*_ and *PI*_*tg*_ values are sufficiently far away from these boundaries (see Supplementary Figure S2). And when they are close to these boundaries, they likely belong to features for which H_0_: *β*_0*g*_ = 0.5 does not hold, such that the null hypothesis will be rightly rejected despite violation of the normality assumption. If necessary, a logit transformation of the PI could be contemplated, although this complicates interpretation.

### Harnessing the high-dimensionality provides precise observation weights

Our procedure for attaching observations weights is based on *spicyR* [16], but with the following modifications. We take the PI of bivariate nearest-neighbour distance 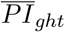 as an example. Consider the set 𝒯 of size |𝒯 | of point patterns with the same fixed and random effects in (9), and calculate the squared departure *m*_*ght*_ = (*PI*_*ght*_ −|𝒯 |^−1^∑_*τ*∈𝒯_ *PI*_*ghτ*_) ^2^ for every *t*. This quantity is comparable to 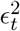, but is estimated for every *t* separately without fitting the mixed model. Next, call *l*_1*t*_ = min(*N*_*gt*_, *N*_*ht*_) and *l*_2*t*_ = max(*N*_*gt*_, *N*_*ht*_) such that *l*_1*t*_ ≤ *l*_2*t*_. log(*m*) is then modeled as a function of log(*l*_1_) and log(*l*_2_) through a decreasing bi-variate spline *s* over all feature pairs and point patterns: 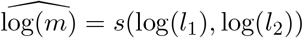. Finally, exp [*s*(log(*l*_1_), log(*l*_2_))]^−1^ is used as observation weight in (9). An example is shown in Supplementary Figure S4. As expected, the value of *l*_1_ has a much stronger influence on the variance than *l*_2_ as it is the limiting source of information on the PI. *spicyR* is different in that it treats *l*_1_ and *l*_2_ equally in its variance model, and calculates *m* with respect to the grand mean u-statistic over all point patterns 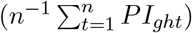. For univariate nearest neighbour PIs, a similar univariate spline as a function of *N*_*gt*_ is fitted (see Supplementary Figure S3).

### Log-ratios are portable measures for neighbourhood enrichment

Neighbourhood enrichment (NE) presents an interesting alternative for *smoppix* for analyzing spatial omics data on the single molecule scale [22, 23, 24]. For univariate enrichment analysis, the proportion of other molecules of feature *g* within a certain range *r* of point *i* is calculated as *p*_*gi*_ for each *i*. Throughout this work, we use as range *r* the median of one twentieth of the longest dimension of all point patterns in the dataset. The proportion *p*_*gi*_ is averaged over all *N*_*gt*_ molecules *i* in point pattern *t* to obtain *p*_*gt*_, and divided by the overall relative abundance of feature *g* in the point pattern: 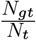. When this ratio exceeds 1, it indicates enrichment, and when it falls below 1 depletion. The methodology to account for replicated experiments is underdeveloped however. Schürch et al. [37] regress log(*p*_*gt*_) on 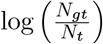and other covariates, but we used 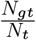 as an offset and calculate the log-ratio 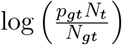 for use as outcome in a linear model like (9).

## Notes

https://www.ebi.ac.uk/biostudies/bioimages/studies/S-BIAD1942

### Competing Interest Statement

The authors have declared no competing interest.

### Summary of Updates

The author affiliations were updated and their order changed. Data files were added to the supplementary

