## Supplementary information for "Unified nonparametric analysis of single-molecule spatial omics data using probabilistic indices"

### Supplementary material for 'Unified nonparametric analysis of single-molecule spatial omics data using probabilistic indices'

Stijn Hawinkel, Xilan Yang, Ward Poelmans, Hans Motte, Tom Beeckman and Steven Maere

**4 Software versions** **45**

### 1 Real data analysis

#### 1.1 Discoveries on *S. moellendorfii* roots are confirmed by HCR RNA-FISH

In the initial *S. moellendorfii* root study by Yang et al. [1], all transcripts measured are localized towards the center of the roots (see Figure S1), since the genes analyzed are primarily active in the vasculature. Hence we use the background of all other transcripts as baseline to call co- or antilocalization using *smoppix*. The sections were taken at different planes of the root, but are considered here as repeated measures. All images from this initial study can be found at https://bioinformatics.psb.ugent.be/webtools/spatial-transcriptomics/selaginella/

The distributions of the estimated PIs per point pattern over all features are shown in Figure S2. The weighting functions for univariate and bivariate nearest-neighbour distance PIs are shown in Figures S3-S4.

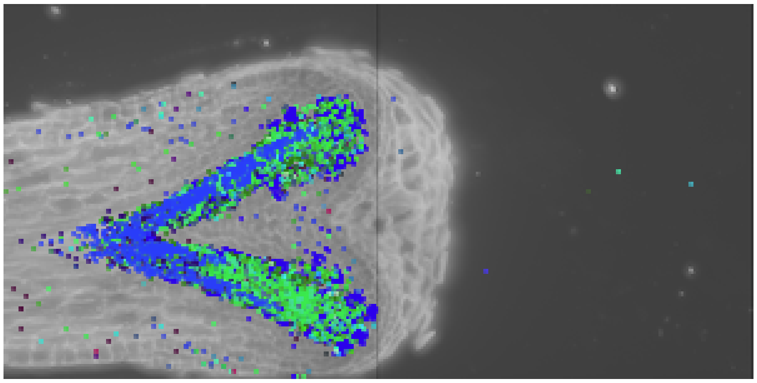

Figure S1: Calcofluor white staining of root 2, section 2 at day 0 of the *smFISH* experiment by Yang et al. [1] with molecules of two example transcripts shown as dots. The cell walls are visible in light grey. More images can be found at https://bioinformatics.psb.ugent.be/webtools/spatial-transcriptomics/selaginella/.

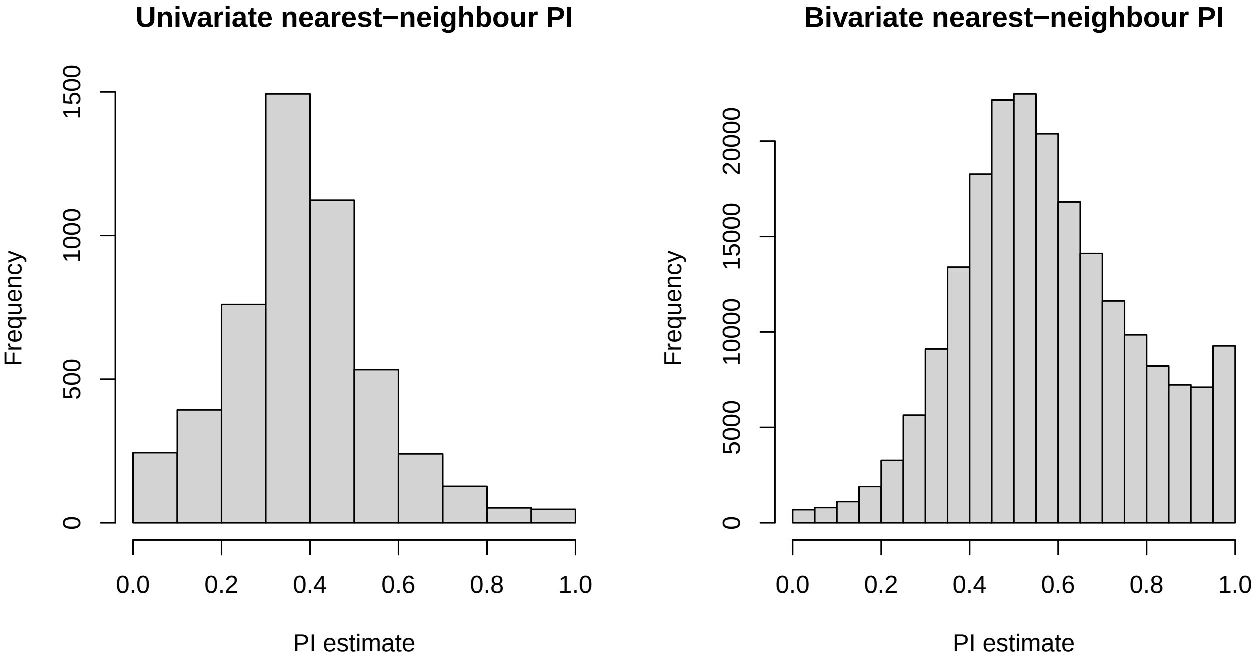

Figure S2: Histograms of estimated univariate (left) and bivariate (right) nearest-neighbour PIs for the Yang2023 *S. moellendorfii* root dataset.

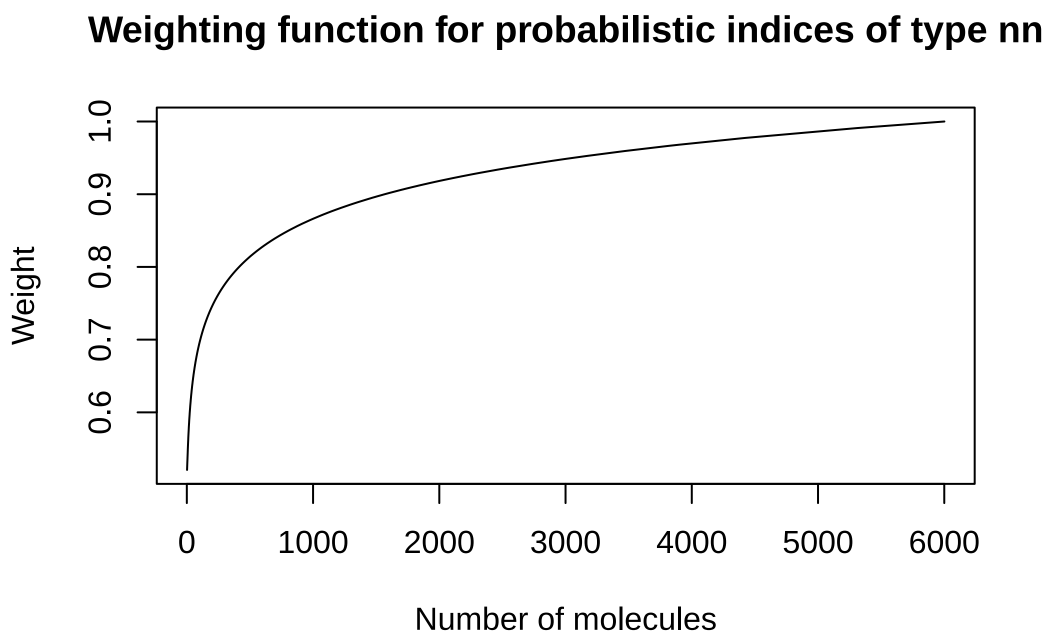

Figure S3: Illustration of the weighting function for univariate nearest-neighbour distances on the Yang2023 *S. moellendorfii* root dataset.

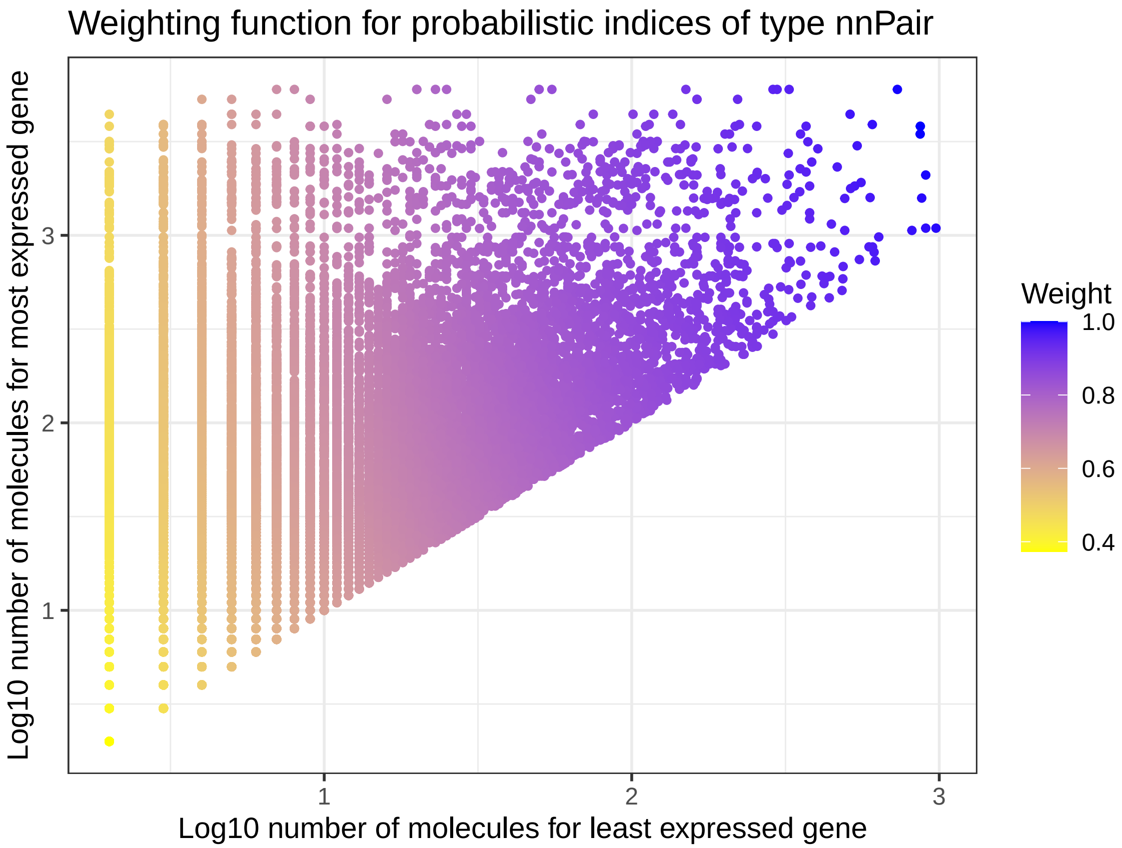

Figure S4: Illustration of the weighting function for the bivariate nearest-neighbour distance PI of the spatial transcriptomics dataset on *S. moellendorfii* roots. The modelled weight is shown in colour as a function of number of molecules in the least expressed (x-axis) and most expressed gene of the pair (y-axis).

We find 177 transcript pairs with significant colocalization and 1708 with antilocalization. The transcript pairs with the most significant colocalization and antilocalization are shown in Figures S5-S6.

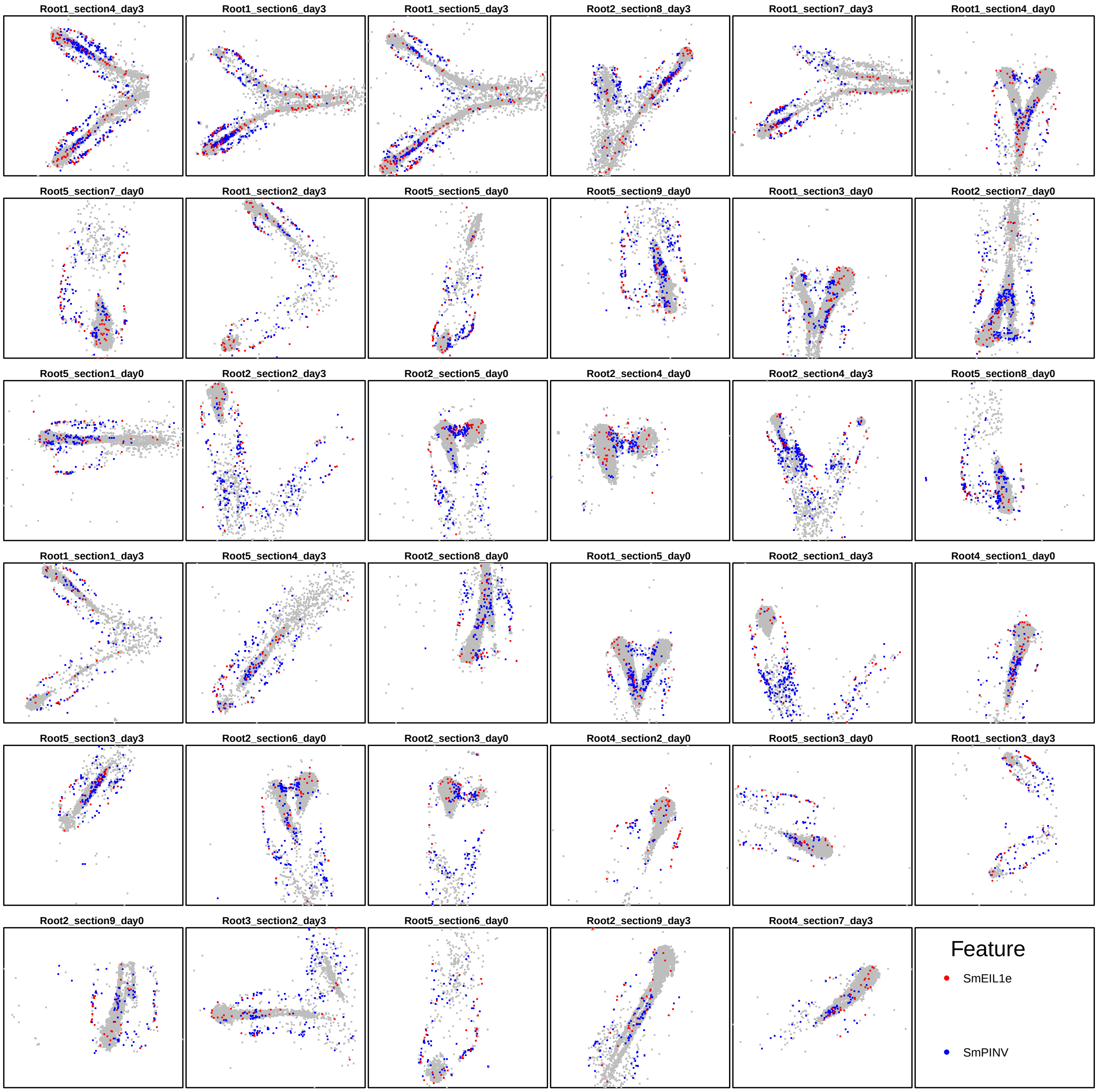

Figure S5: Most significantly colocalized gene pair in the *S. moellendoriff* root dataset according to *smoppix*. Despite its modest effect size (PI = 0.39), it owes its high significance to its consistency across replicates.

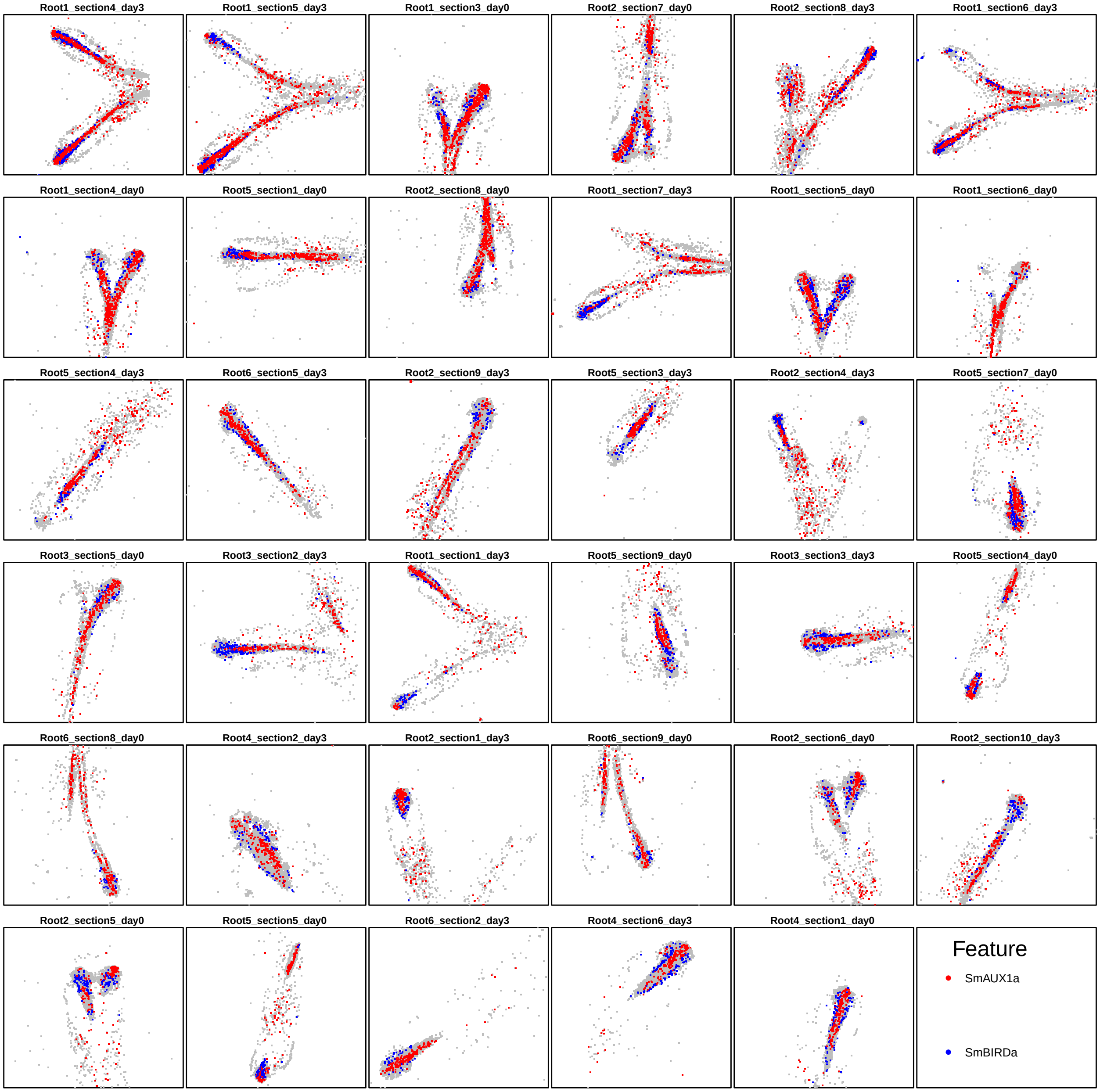

Figure S6: Gene pair with most significant antilocalization in the *S. moellendoriff* root dataset according to *smoppix*.

##### 1.1.1 Authors' findings

The authors of the original study came to the following conclusions based on pseudo-segmentation, calculation of Pearson correlation and hierarchical clustering [1]:

- SmTMO5b and SmVND1 are coexpressed
- SmWOX13a and SmCYCD3;3b are coexpressed (in day 3 root 1 sample)
- SmWOX13a and SmBRN are coexpressed
- SmSHRa, SmSCRa, SmSCRb, SmRBRa, SmBIRDa, and SmCYCD3;3a genes are coexpressed

We can confirm some these colocalizations using *smoppix* at a nominal FDR of 0.05, but not the pairs shown in Table S1. Plots for some of these pairs are shown in Figure 2 in the main text and Figure S7 here. SmBIRDa is often found in two strips at the outer edges of the point cloud, whereas SmCYCD3;3a is more localized towards the center, contradicting colocalization. Similarly, SmSHRa often lies closer to the root tip than SmSCRa. The original analysis based on pseudosegmentation disregards distances between pseudosegments, and is thus indifferent to whether molecules are in adjacent pseudosegments or on the other side of the root. In addition, part of the finer spatial patterns are lost by binning.

| | PI estimate | SE | p-value | Adjusted p-value |
|---:|---:|---:|---:|---:|
| SmWOX13a–SmBRN | 0.43 | 0.06 | 2.21e-01 | 3.30e-01 |
| SmSHRa–SmRBRa | 0.48 | 0.02 | 3.21e-01 | 4.42e-01 |
| SmRBRa–SmBIRDa | 0.50 | 0.01 | 7.66e-01 | 8.32e-01 |
| SmSHRa–SmSCRa | 0.50 | 0.02 | 8.10e-01 | 8.67e-01 |
| SmRBRa–SmCYCD3;3a | 0.51 | 0.01 | 4.30e-01 | 5.49e-01 |
| SmSCRb–SmCYCD3;3a | 0.52 | 0.02 | 3.19e-01 | 4.41e-01 |
| SmBIRDa–SmCYCD3;3a | 0.53 | 0.01 | 2.00e-03 | 7.00e-03 |
| SmSCRa–SmSCRb | 0.54 | 0.03 | 8.70e-02 | 1.59e-01 |
| SmSHRa–SmCYCD3;3a | 0.55 | 0.02 | 3.00e-03 | 9.00e-03 |
| SmSCRa–SmRBRa | 0.55 | 0.01 | 8.00e-03 | 2.20e-02 |
| SmSCRb–SmBIRDa | 0.58 | 0.02 | 1.00e-03 | 3.00e-03 |
| SmSHRa–SmSCRb | 0.58 | 0.03 | 4.00e-03 | 1.40e-02 |
| SmSCRb–SmRBRa | 0.58 | 0.03 | 2.00e-03 | 8.00e-03 |
| SmWOX13a–SmCYCD3;3b | 0.59 | 0.03 | 2.60e-02 | 5.90e-02 |

Table S1: P-values of nonsignificant (adjusted p-value > 0.05) or antilocalized (PI > 0.5) features according to *smoppix*, found colocalized by the authors. The table was sorted by PI estimate.

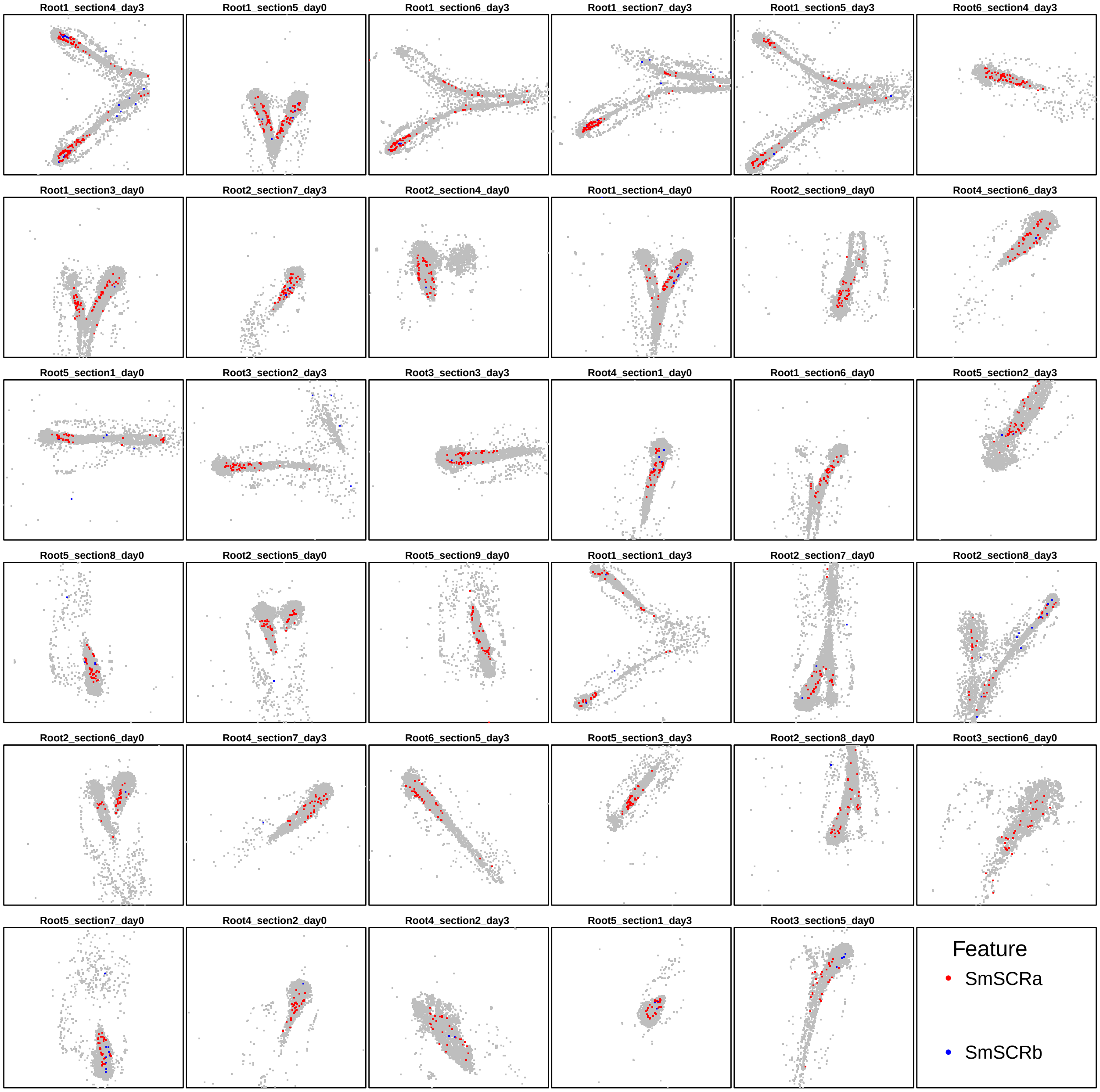

Figure S7: Transcript pair SmSCRa–SmSCRb which was found to be coexpressed by the authors but antilocalized in the *smoppix* analysis.

##### 1.1.2 Comparison with *spicyR* and neighbourhood enrichment

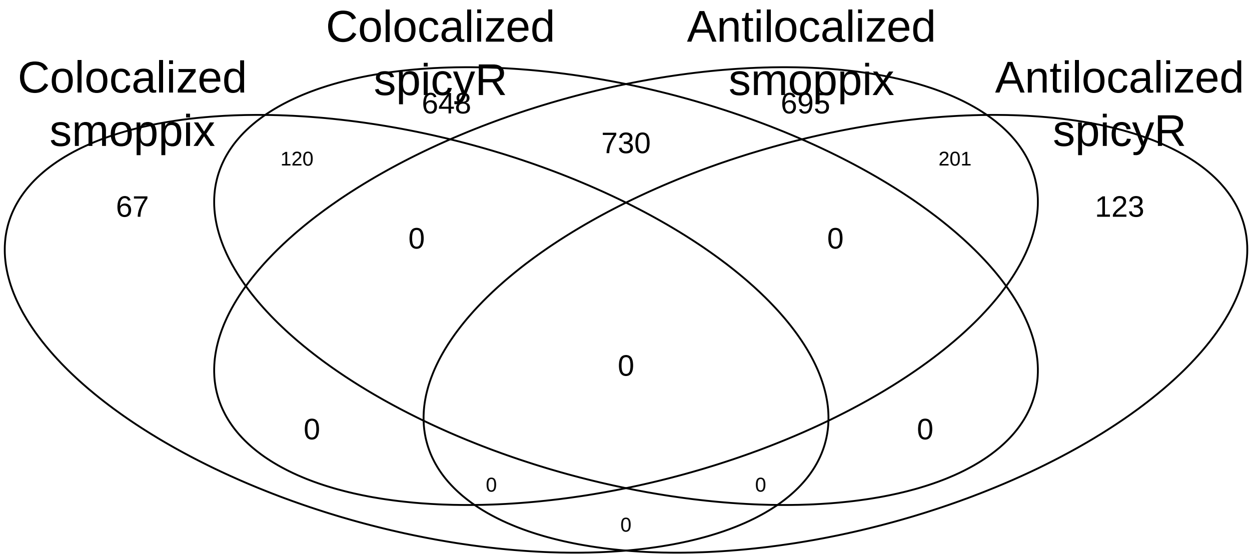

Figure S8: Venn diagram of co- and antilocalized transcripts in the dataset by Yang et al. [1] according to *smoppix* and *spicyR*.

The overlap between the results of *smoppix* and *spicyR* [2] is shown in Figure S8. As an example of different results, *smoppix* does not consider SmCASP1–SmIAA9b significant, whereas *spicyR* believes it is colocalized (see Figure S9). We also applied neighbourhood enrichment (NE), employing the *average_percentage_of_cells_within_radius* function from the *SPIAT* package [3] to calculate the neighbourhood proportions $p_{gi}$. The Venn diagram with overlap in results with *smoppix* is shown in Figure S10, revealing that NE makes fewer discoveries than *spicyR*, but they correspond better to those of *smoppix* than those of *spicyR*. The only gene pair with disagreement is shown in Figure S11.

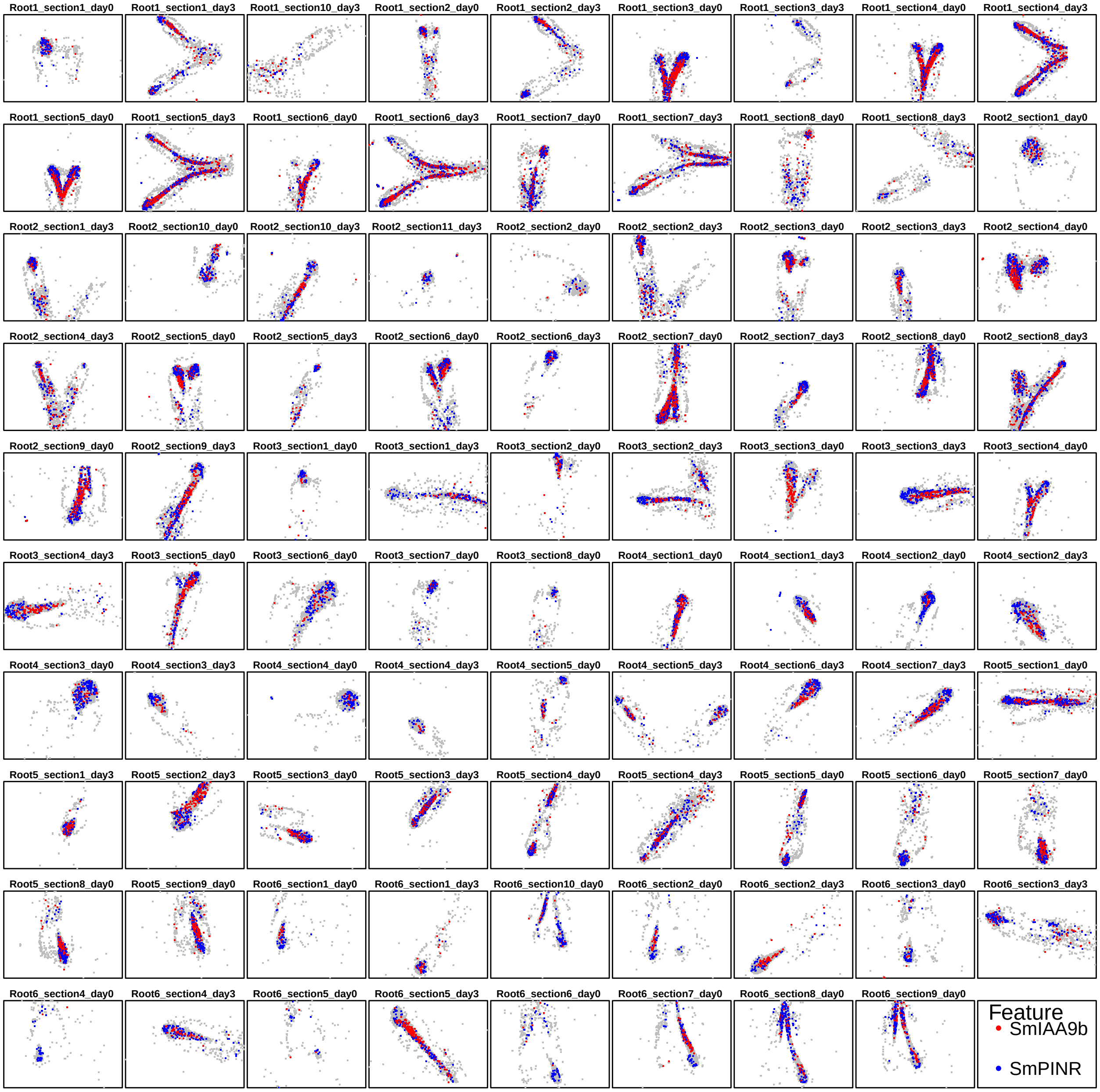

Figure S9: Transcript pair SmIAA9b–SmPINR which is considered colocalized by *spicyR* but not significant in the *smoppix* analysis.

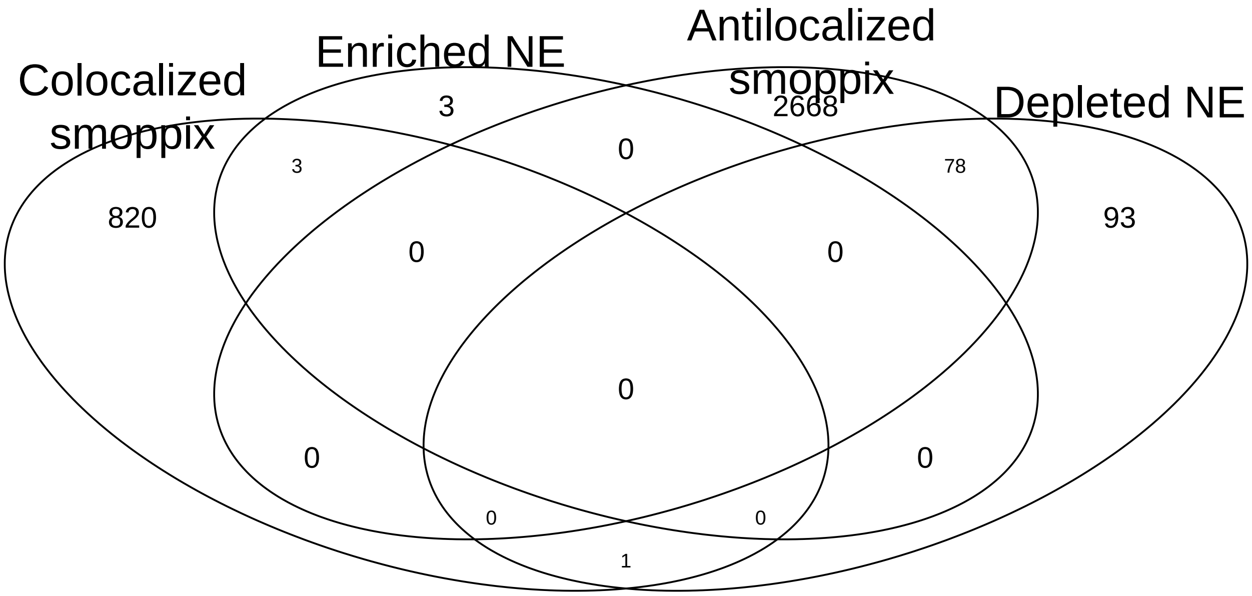

Figure S10: Venn diagram of co- and antilocalized transcripts according to *smoppix* and NE.

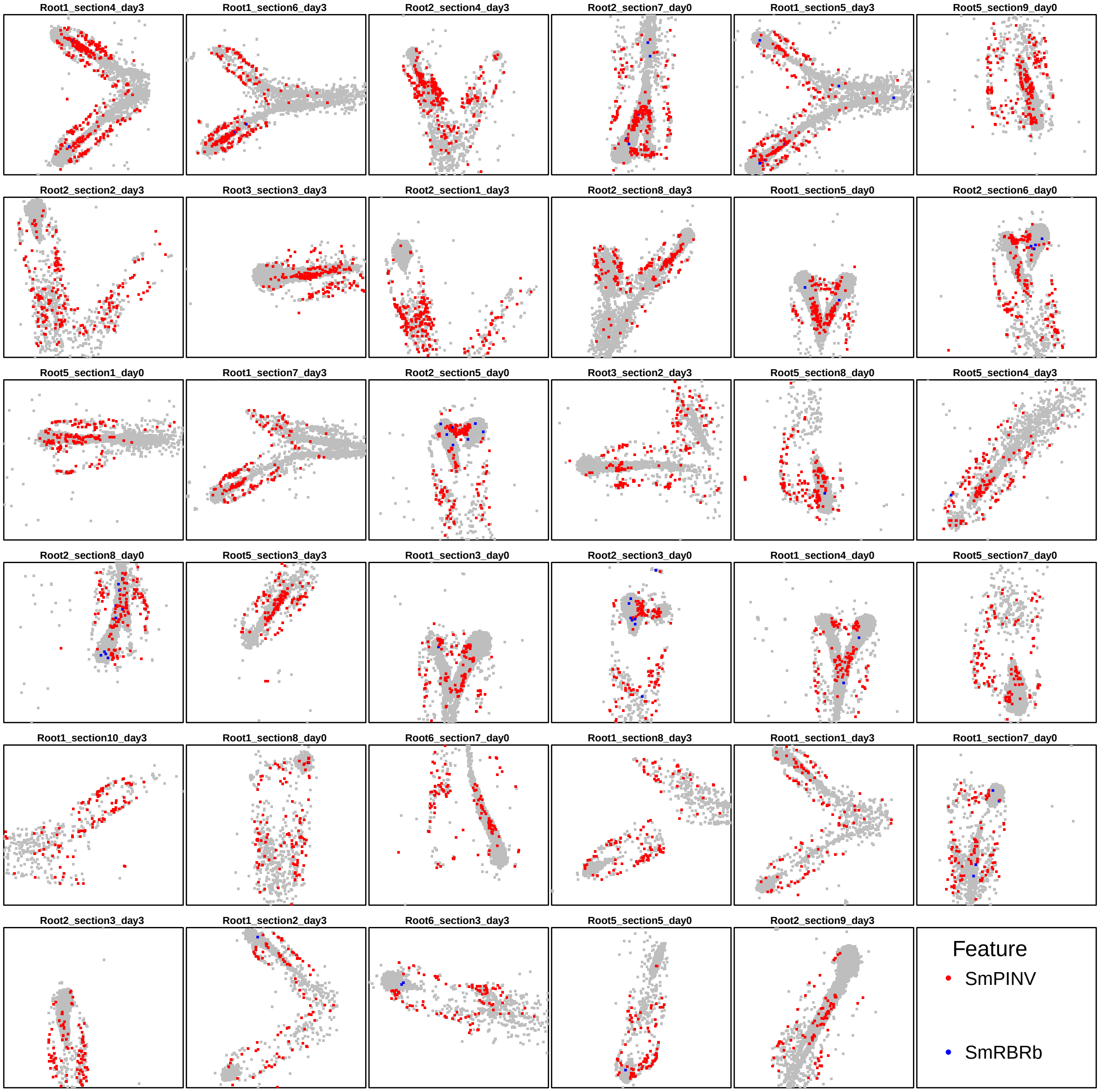

Figure S11: Example of gene pair found antilocalized by *smoppix* but neighbourhood enriched by NE. In this case, the enrichment found by NE is mutual, i.e of SmPINV in the neighbourhood of SmRBRb and vice versa.

##### 1.1.3 Confirmatory experiment with HCR RNA-FISH

The results on smFISH data by *smoppix*, *spicyR*, NE and the authors' original analysis for the eight genes chosen for HCR RNA-FISH confirmatory experiments for are shown in Table S2. *spicyR* and the original analysis declare almost all gene pairs colocalized, whereas NE finds no significant results. *smoppix* agrees with spicyR and the authors' analysis on colocalization of the pairs SmPINR–SmSGNa and SmPINS–SmSGNb (PI $< 0.5$ and adjusted p-value $< 0.05$), but declares the other pairs significantly antilocalized (PI $> 0.5$ and adjusted p $< 0.05$). The scatterplots of the smFISH experiment of the SmBIRDa–SmCYCD3;3a and SmPINR–SmSGNa are shown in Figure 2 in the main text.

| | Authors' analysis | Smoppix PI | Adj. p-value | spicyR u-statistic | Adj. p-value | NE | Adj. p-value |
|---|---|---|---|---|---|---|---|
| SmBIRDa–SmCYCD3;3a | Colocalized | 0.53 | 5.3e-03 | 529.2 | 5.5e-04 | 0.13 | 1.0e+00 |
| SmPINR–SmSGNa | Colocalized | 0.45 | 1.1e-03 | 442.8 | 1.5e-04 | 0.04 | 1.0e+00 |
| SmPINS–SmSGNb | Colocalized | 0.39 | 3.6e-06 | 831.8 | 8.9e-27 | 0.22 | 1.0e+00 |
| SmRBRa–SmSCRa | Colocalized | 0.55 | 1.8e-02 | 455.2 | 2.1e-03 | -0.28 | 1.0e+00 |
| SmRBRa–SmSCRb | Colocalized | 0.58 | 6.3e-03 | 302.8 | 2.3e-05 | -0.52 | 1.0e+00 |
| SmBIRDa–SmSCRb | Colocalized | 0.58 | 2.5e-03 | 235.0 | 5.5e-06 | -0.26 | 1.0e+00 |
| SmSHRa–SmSCRb | Colocalized | 0.58 | 1.1e-02 | 33.6 | 8.2e-01 | -0.24 | 1.0e+00 |
| SmCYCD3;3a–SmSHRa | Colocalized | 0.55 | 7.7e-03 | 411.2 | 1.4e-03 | -0.15 | 1.0e+00 |

Table S2: Table of original results and *smoppix*, *spicyR* and neighbourhood enrichment (NE) effect size estimates and adjusted p-values on smFISH data, for transcript pairs investigated further in HCR RNA-FISH experiments.

HCR RNA-FISH was conducted as described previously [4] and according to the manufacturer's guidelines (https://www.molecularinstruments.com/hcr-rnafish-protocols). Probes for genes were designed and synthesized by Molecular Instruments (www.molecularinstruments.com). The Selaginella roots were fixed by 4% formalin-aceto-alcohol (FAA) and permeabilized by series of ethanol and methanol [5]. Partial cell wall digestion was done with the enzyme mix described previously [6, 4, 7]. After enzyme digestion, roots were fixed in 10% (v/v) formaldehyde, Proteinase K treated and fixed in 10% (v/v) formaldehyde again. Cell wall staining occurred by 0.1% Calcofluor White for 30 min in ClearSee, after which the samples were stored in ClearSee. Confocal imaging of roots was performed using a Zeiss LSM710 confocal microscope for the first experiment and a Leica Stellaris 5 confocal microscope for the second experiment, imaging green fluorophores by 561 nm excitation and 580-650 nm wavelength detection, red fluorophores by 633 nm excitation and 650-755 nm wavelength detection, and Calcofluor White by 405 nm excitation and 410-525 nm wavelength detection.

Sample images per gene pair are shown in Figure 3 in the main text, the full stack of images is available from the BioStudies repository [8] with accession number S-BSST2022. As background correction, pixels falling below the average intensity plus two times the standard deviation of either colour channel were omitted before calculating Pearson correlations between pixels, similar to Choi, Beck, and Pierce [9]. The relationship between the estimated Pearson correlation, its standard error and the z-coordinate are explored in Figures S12-S13. Given its observed dependence on root and z-coordinate, we decided to model the Pearson correlation $\rho_{ig}$ in z-stack image $i$ of root $g$ using the following mixed-effects model:

$$\rho_{ig} = \beta_0 + \beta_z z_i + b_{gz} z_i + \sum_{r=1}^{R} b_{gr} x_{ir} + \epsilon_i, \tag{1}$$

with $z_i$ the mean-centered z-coordinates, $\beta_z$ the fixed effect slope, $b_{gz}$ the random slope of root $g$ and $b_{gr}$ the random intercept of root $r$ with $x_{ir}$ a dummy variable indicating root. Observation weights inversely proportional to the variances of the Pearson correlation estimates, as provided by the *cor.test* function in the *stats* R-package, are used in the mixed model. P-values were adjusted with Benjamini-Hochberg correction within each of the two HCR RNA-FISH experiments. The results of the test of the hypothesis $H_0 : \beta_0 = 0$ on the eight selected gene pairs are shown in Table 1 in the main text.

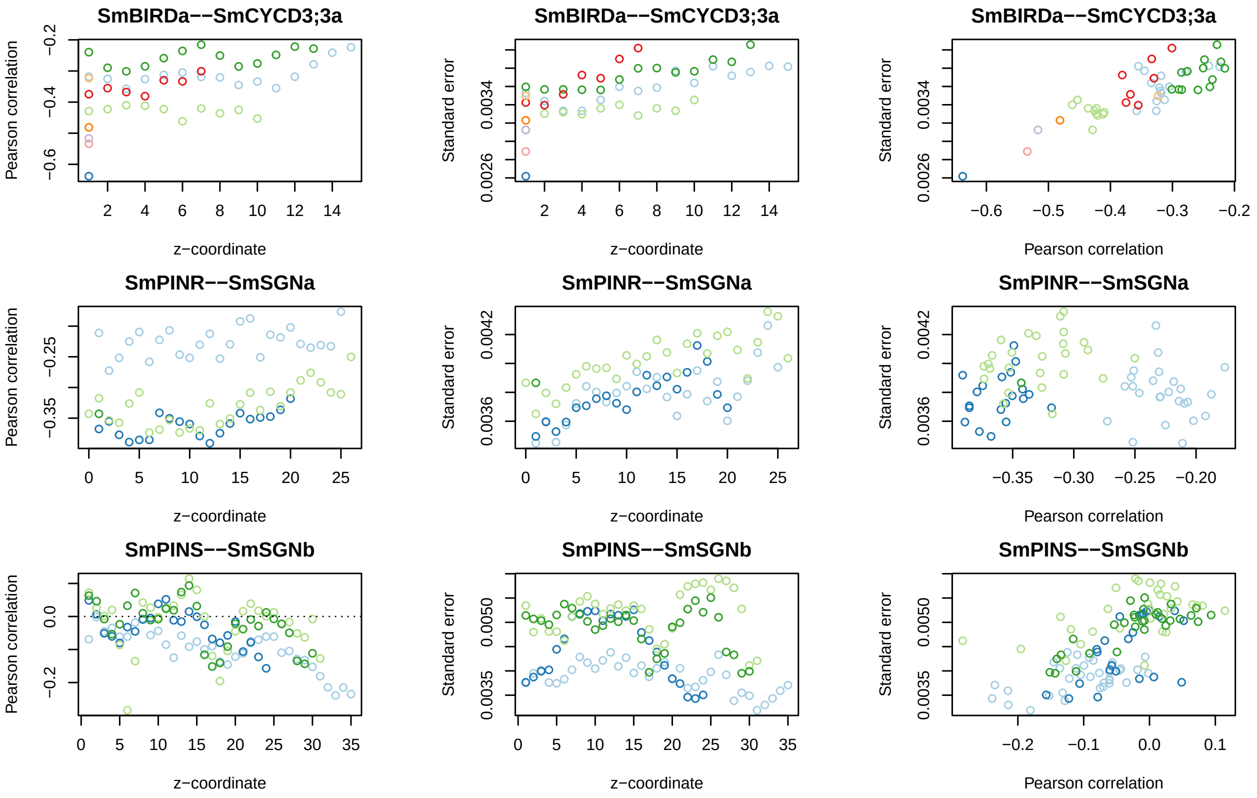

Figure S12: Scatterplots of the estimated Pearson correlation (left column) and its standard error (middle column) as a function of z-coordinate, and the standard error as a function of Pearson correlation (right colum), coloured by root for the three transcript pairs (plot titles) for the first experiment. In the left plots, the dotted horizontal line indicates zero correlation.

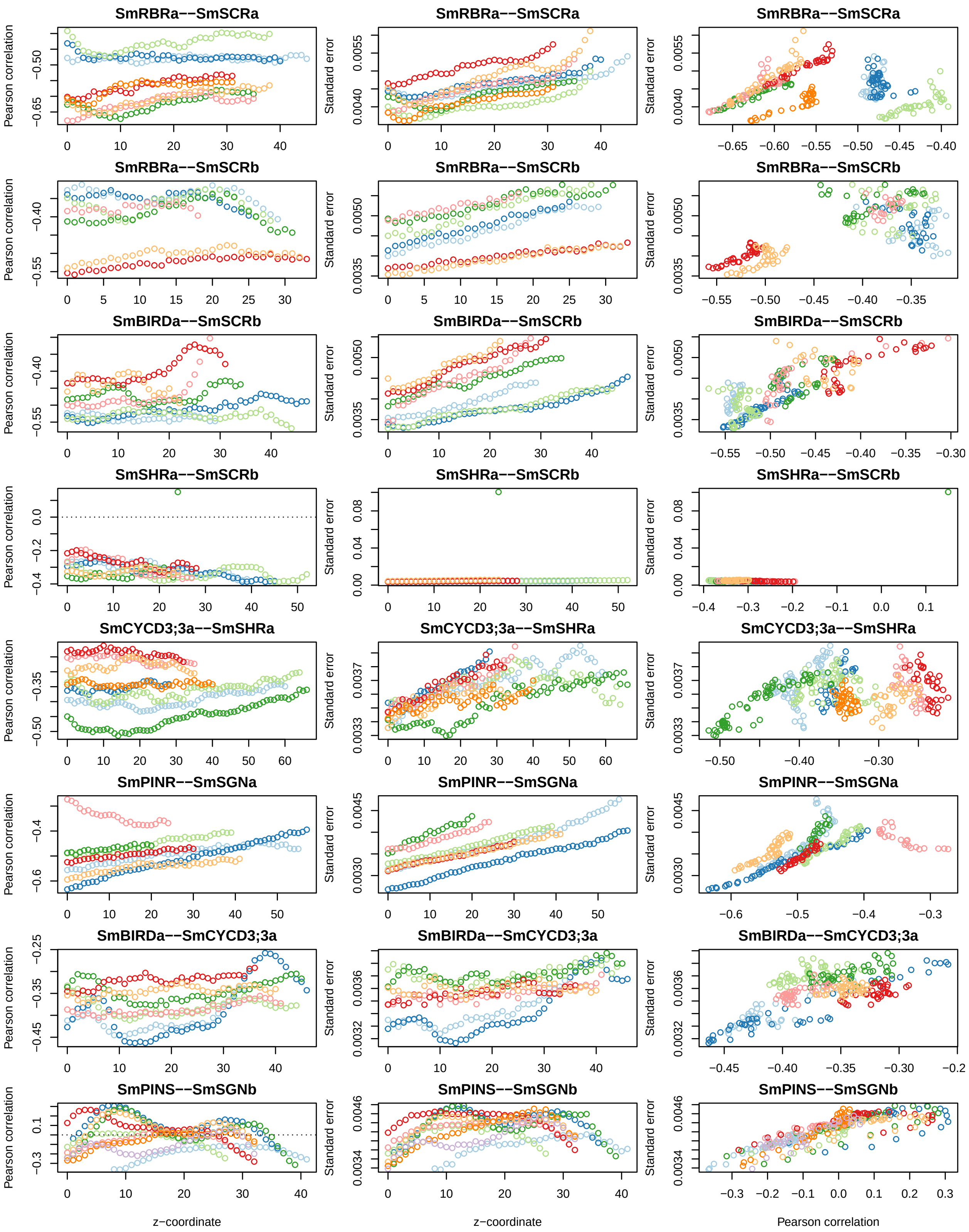

Figure S13: Scatterplots of the estimated Pearson correlation (left column) and its standard error (middle column) as a function of z-coordinate, and the standard error as a function of Pearson correlation (right column), coloured by root for the eight transcript pairs (plot titles) for the second experiment. In the left plots, the dotted horizontal line indicates zero correlation. The outlier for SmSHRa–SmSCRb corresponds to an image with little signal, which is downweighted in the analysis because of its high standard error.

#### 1.2 *smoppix* finds intracellular localization patterns in mouse fibroblast cells

We reanalysed a dataset by Eng et al. [10], containing measurements of 10,000 genes in NIH/3T3 mouse fibroblast cells. Five z-slices with z-steps of 1 $\mu$m were taken across multiple fields of view (FOVs). To demonstrate robustness, the experiment was performed twice on separate sections (the experiment variable). The cells were gated by hand based on DAPI staining. Here we test for vicinity to and remoteness from cell wall and cell centroid and for within-cell aggregation for one gene at the time. No bivariate analyses were performed as they would take too long. The bivariate analysis was restricted to the 500 most highly expressed genes for computational reasons, an analysis on all 10,000 genes would take roughly 20,000 cpu hours. *smoppix* found 14 genes significantly close to the cell edge and 4390 genes significantly close to the centroid compared to CSR within the cell; the most significant ones in each case are shown in Figure 4 in the main text. Also, *smoppix* found 4287 genes aggregated within the cell compared to CSR (see Figure S14).

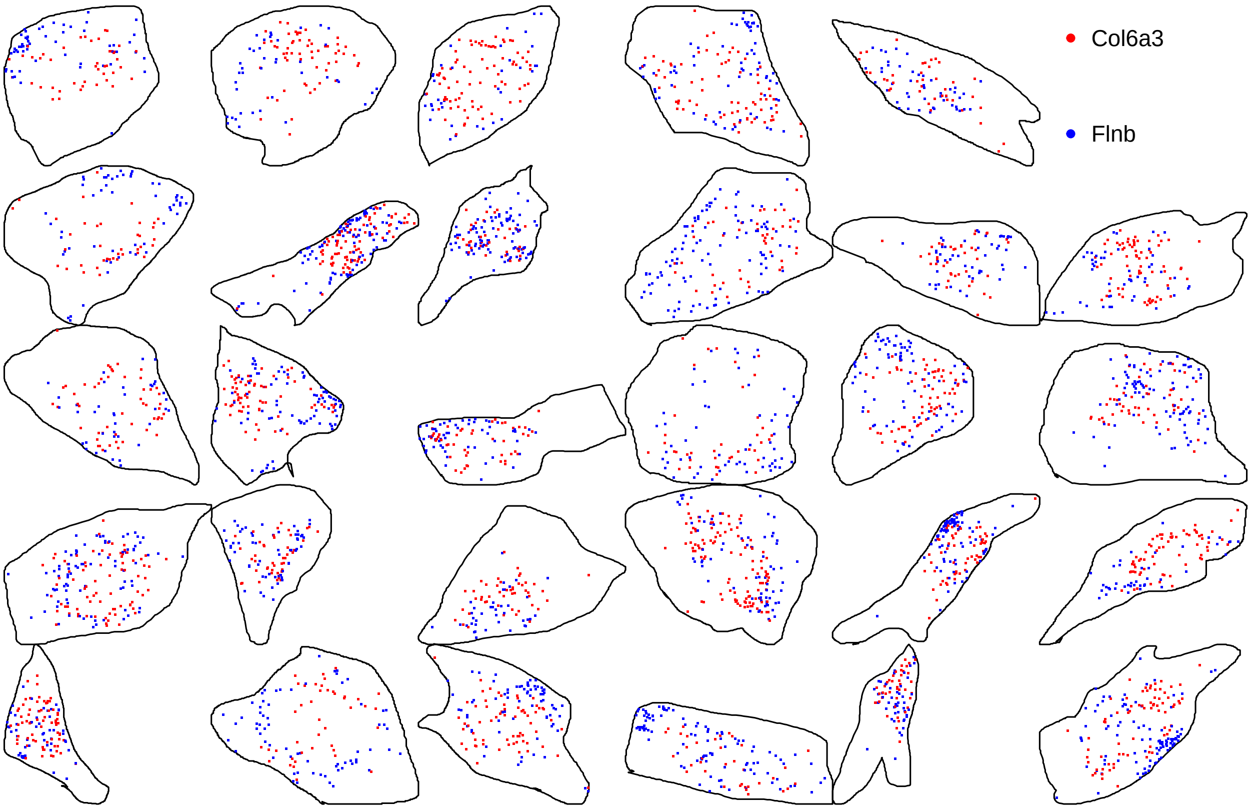

Figure S14: Two genes most significantly aggregated within the cell according to *smoppix*. The cells are not shown in their original location in the section but rather sorted by expression.

##### 1.2.1 Authors' findings

The spatial analysis in the original publication was based on pseudosegmentation into squares of 10 by 10 pixels, calculation of Pearson correlation on the resulting counts and hierarchical clustering. Three main clusters were thus identified: nuclear/perinuclear, cytoplasm and protrusions. An alternative analysis exploiting the full resolution of the data using latent factors was presented by [11], finding similar clusters of genes. There is some overlap between genes found close to the centroid by *smoppix* and in the nuclear/perinuclear region by the authors, and genes found close to the edge by *smoppix* and in protrusions by the authors, respectively (Figure S17). Yet there are also differences, e.g. genes found close to the centroid by *smoppix* but not in the nuclear/perinuclear region by the original analysis (Figure S15) or genes found enriched in the protrusions by the authors but not found close to the edge by *smoppix* (Figure S16). The latter tend to be lowly expressed genes, which may explain why *smoppix* does not find sufficient evidence to call spatial patterning. And of course, enrichment in protrusions not exactly the same as vicinity to cell edge, and the nucleus and the centroid of the cell may not colocalize. Also, the authors' analysis looks for clusters of genes with similar spatial distribution, whereas *smoppix* tests every gene independently.

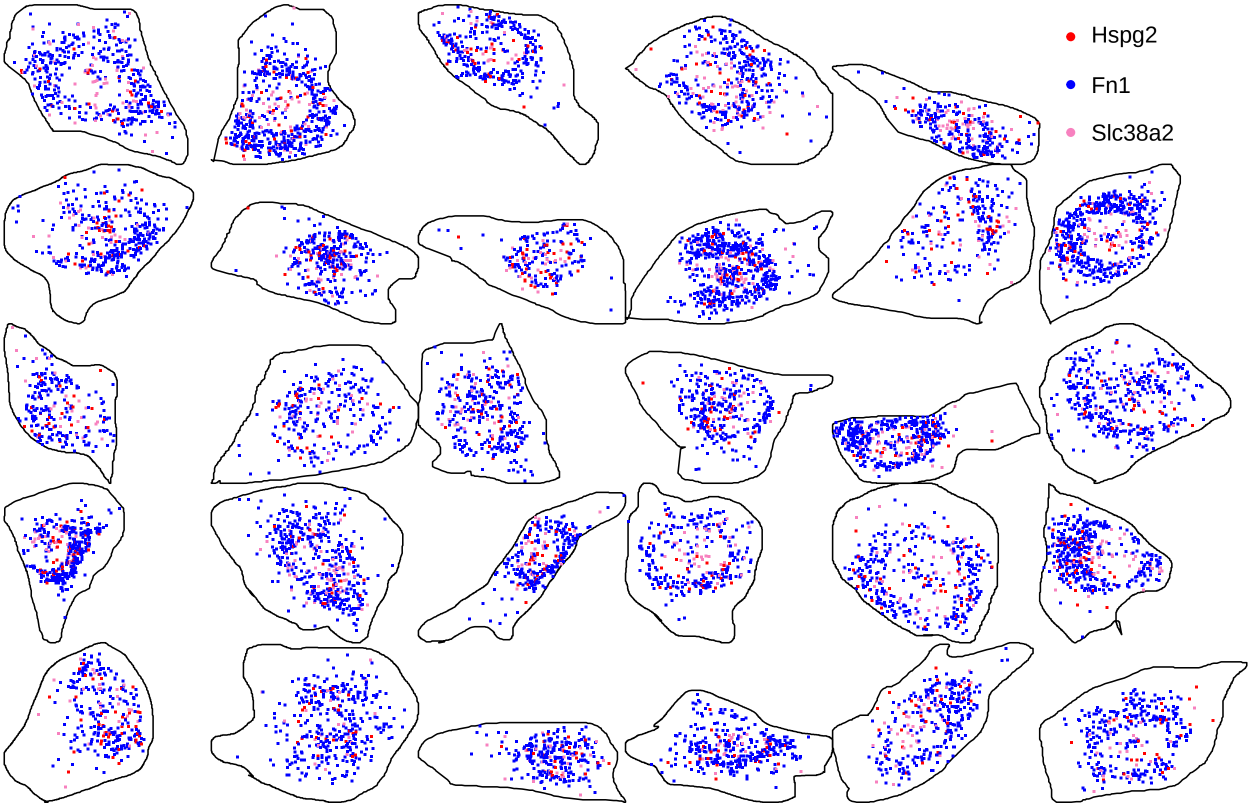

Figure S15: Example genes found close to the centroid by *smoppix* but not in nuclear/perinuclear region by the authors. The cells are not shown in their original location in the section but rather sorted by expression.

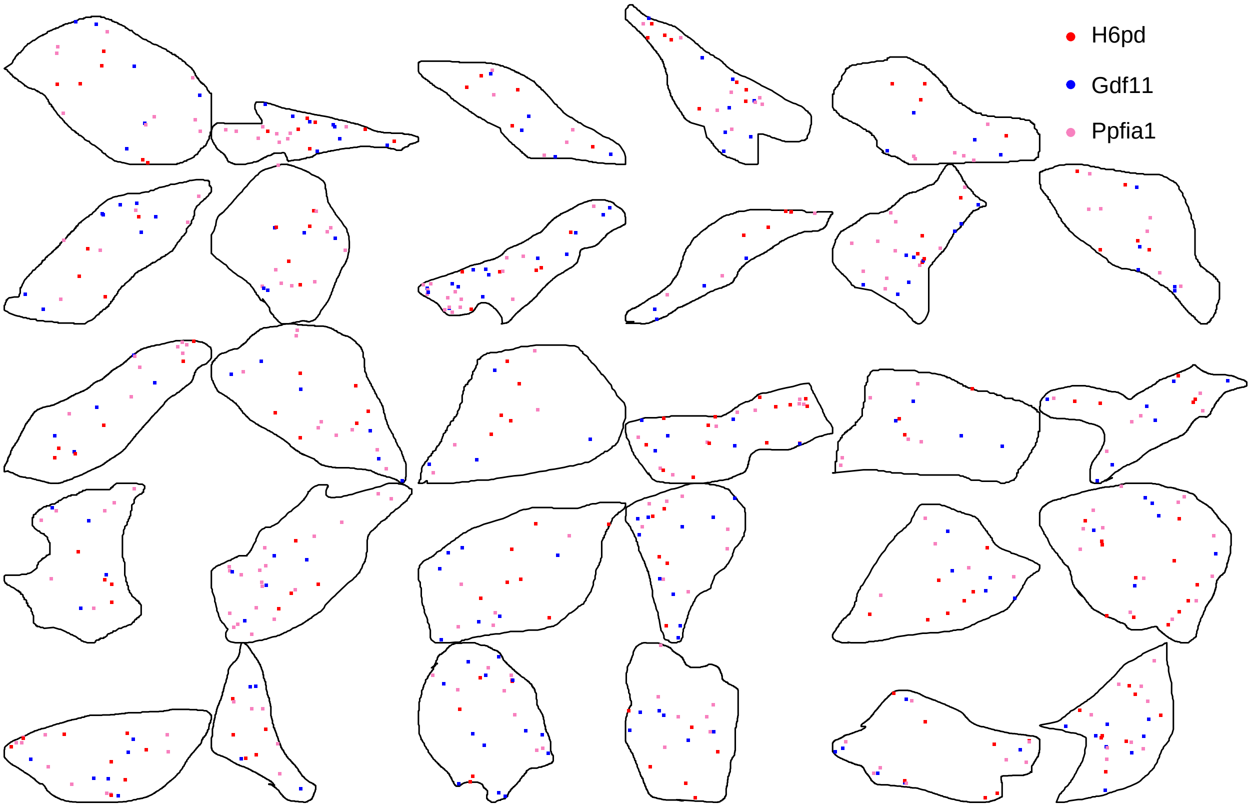

Figure S16: Example genes found in protrusions by the authors but not close to the edge by *smoppix*. The cells are not shown in their original location in the section but rather sorted by expression.

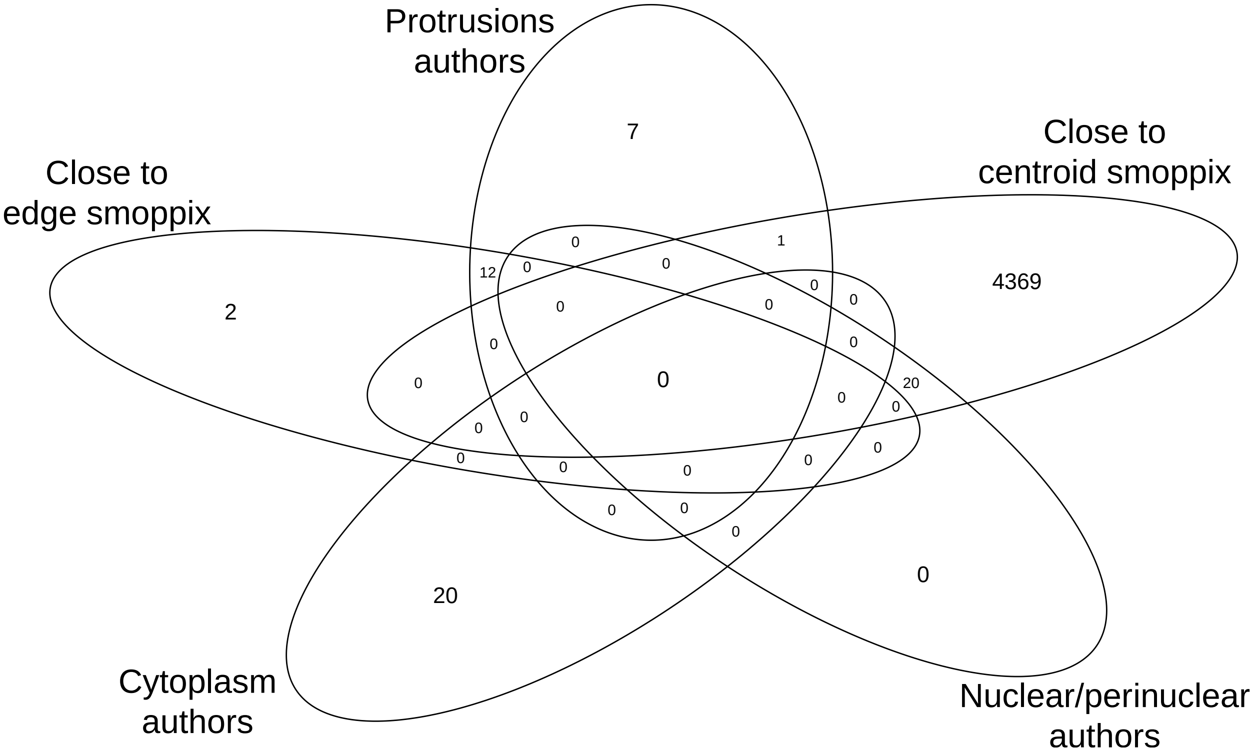

Figure S17: Venn diagram of overlap between results from the authors and from *smoppix*.

#### 1.3 *smoppix* detects shifts in localization patterns over time in bacterial biofilms

Dar et al. [12] profiled *Pseudomonas aeruginosa* biofilms for spatial expression of 108 genes after 10h of growth (7 replicates) and after 35h of growth (3 replicates). The biofilms were scanned at different depths (z-coordinates), but the lower and upper layers often yielded few molecules. The data were analysed in 2D here, so considering each z-plane as a separate replicate, and including a random effect for biofilm instance. We look for differences in spatial organisation between 10h and 35h. The transcript pair with most significant differential spatial localization between 10h and 35h is shown in Figure 5 in the main text. Further differences between both conditions are shown here in Figures S18-S19, demonstrating *smoppix*' ability to detect differences in spatial localization patterns between conditions.

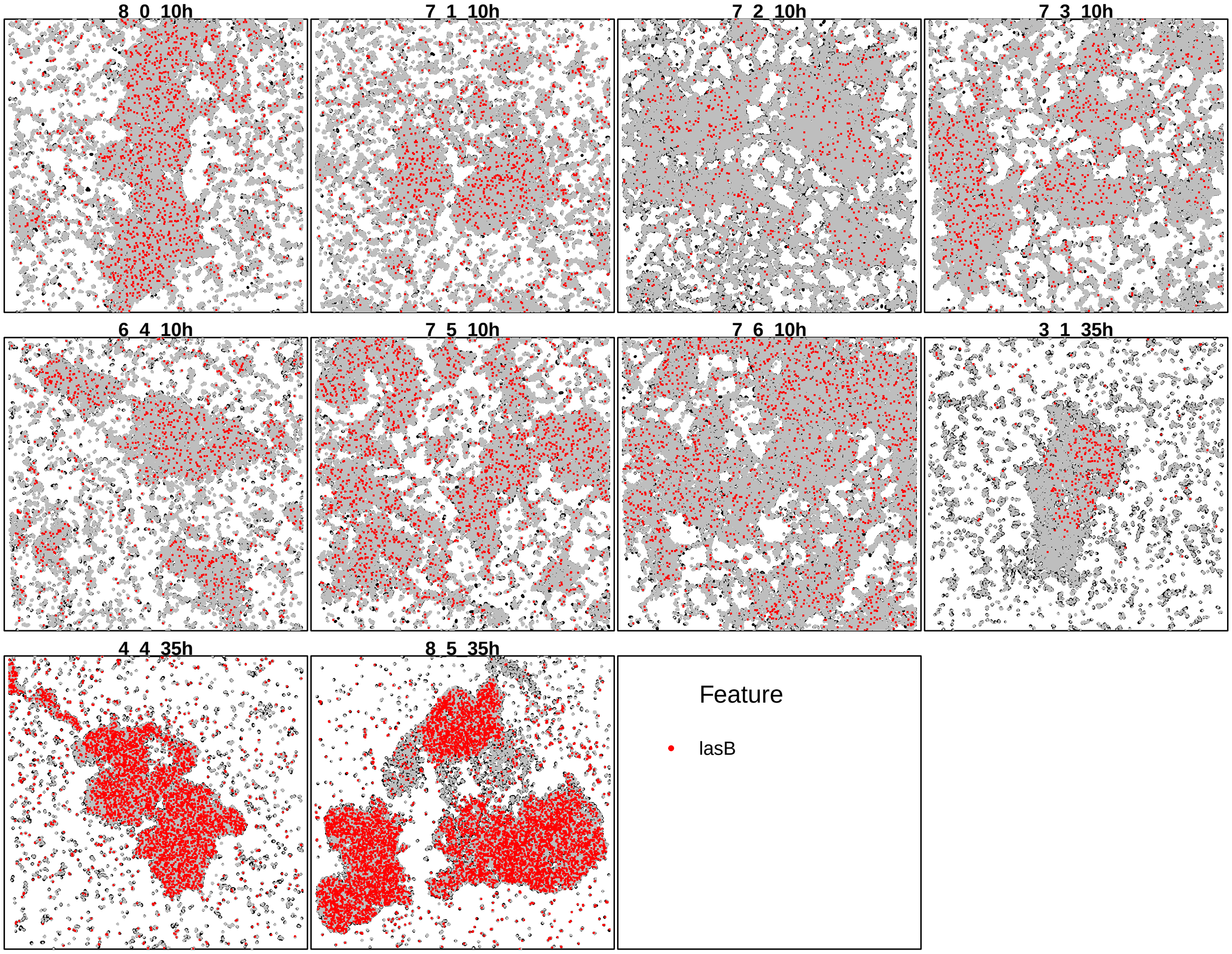

Figure S18: Gene most significantly more regularly spaced at time 35h than 10h according to *smoppix*.

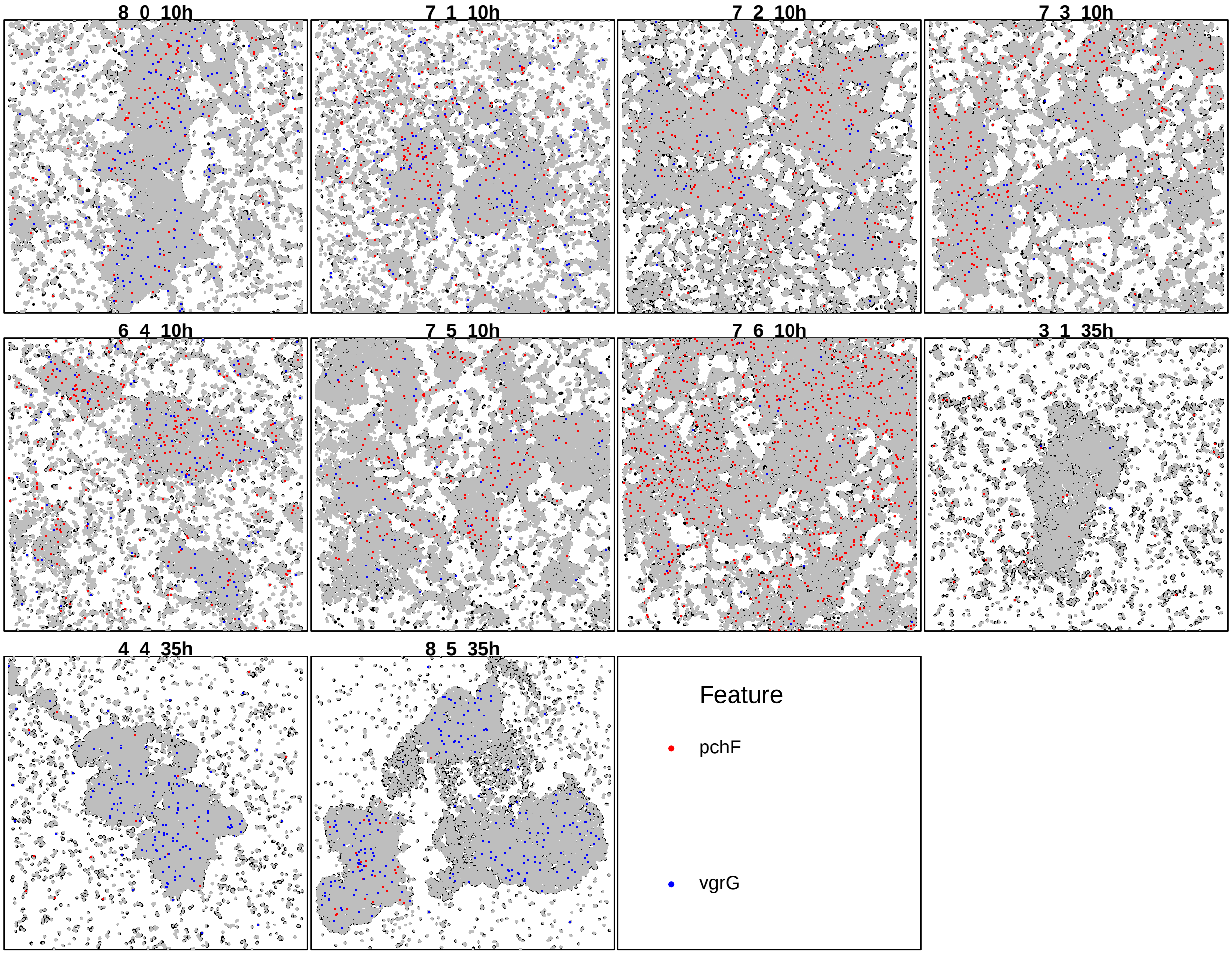

Figure S19: Gene pair most significantly more antilocalized at time 35h than 10h according to *smoppix*.

##### 1.3.1 Authors' findings

Here we compare *smoppix*' results with the author's findings, obtained by pseudosegmentation and Pearson correlation analysis [12]. The authors found the *napA* and *uspL* genes to be expressed in a regularly spaced manner but at a low level in the 35h aggregates, yet we find no evidence for any spatial patterning (p-values 0.174 and 0.276, respectively). Also the *cdrA* transcripts were found to be regularly distributed, we indeed find evidence for regularity (p-value = 0.00636). The authors found the *fliC* transcripts to be more regularly spaced after 35h. We find it to be regularly distributed at both timepoints, but more regular at 10h. The authors found anti-correlation between *fliC* and *pilA* (checkerboard pattern) at 10h, we can confirm this at both timepoints (p-value 0.0316). Moreover, the original publication also found spatial correlation between anaerobic metabolism genes' transcripts, such as those in the denitrification pathway (narG-nirS-norB-nosZ), and the oxidative stress response genes katA, katB, and sodM [12]. *smoppix* confirms colocalization patterns for most denitrification transcripts (see Figure S20 for an example), and between *katB* and *katA*, but not for the other oxidative stress transcripts. *smoppix* also finds associations between oxidative stress and heat-shock protease expression (Figure S21). Yet *smoppix* finds no evidence of colocalization between the denitrification and oxidative stress transcripts. Moreover the authors found that the stress response pattern was also spatially correlated with heat-shock protease expression, including the membrane protease ftsH. *smoppix* does not detect such patterns.

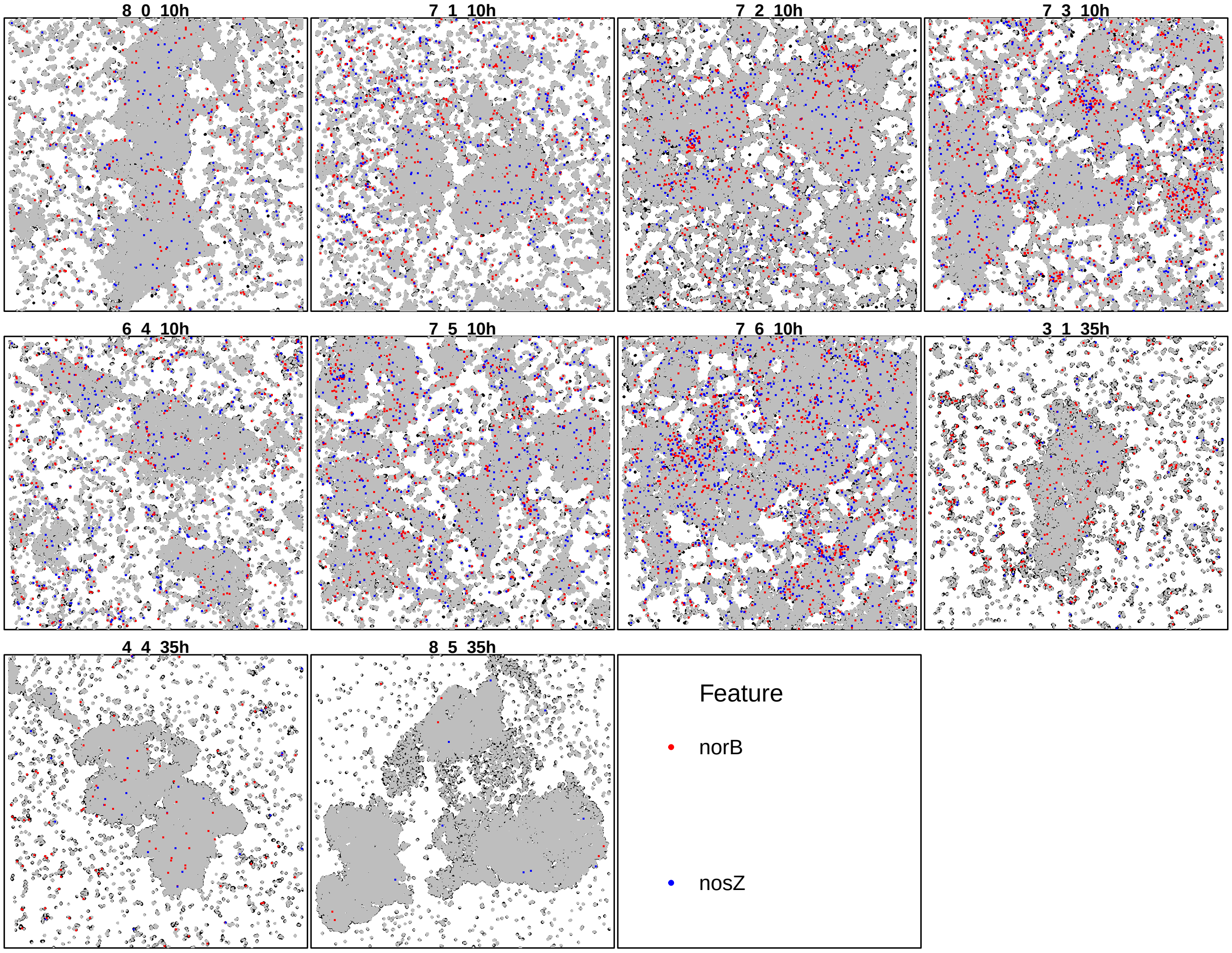

Figure S20: Denitrification gene pair with most significant colocalization according to *smoppix*.

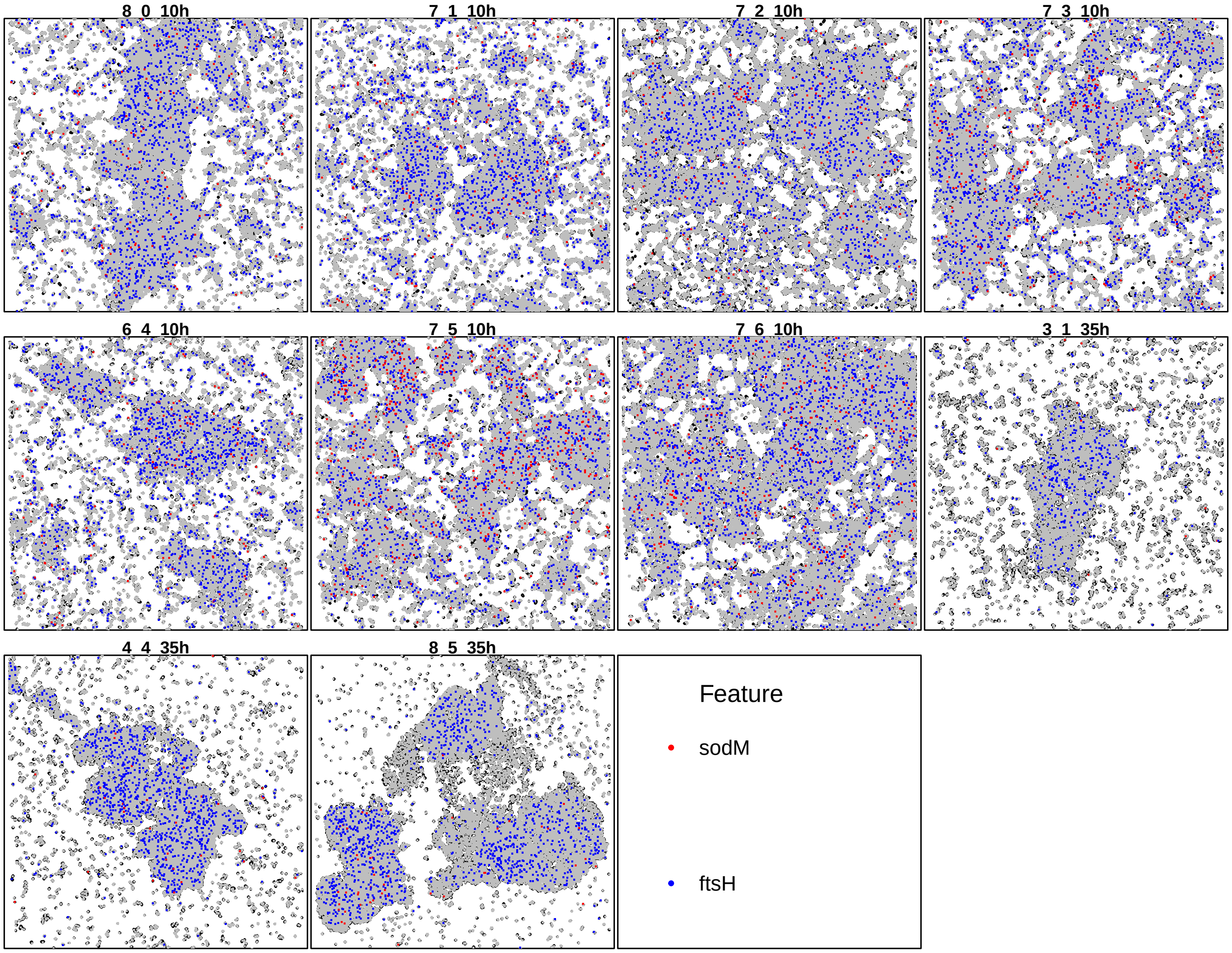

Figure S21: Oxidate stress gene (sodM) and heat shock gene (ftsH): the gene pair with most significant trancript colocalization according to *smoppix*.

##### 1.3.2 Comparison with *spicyR*

The overlap of findings of the univariate methods between *smoppix* and *spicyR* is shown in Figure S22; NE could not be applied to this dataset for computational reasons. *spicyR*'s most significant findings are plotted in Figures S23-S24.

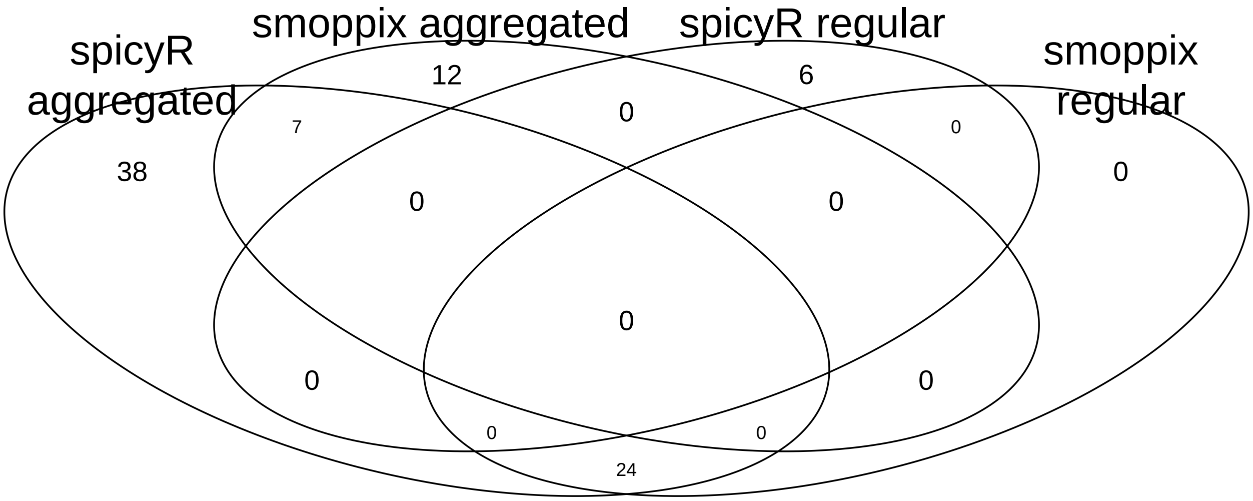

Figure S22: Venn diagram of overlap between *smoppix* and *spicyR* results for univariate analysis.

24 genes were found to be aggregated by *spicyR* but regularly spaced according to smoppix, one example is shown in Figure S25. These results also reflect the difference in the null hypothesis tested. *spicyR* considers only the densities of the genes themselves for estimating a baseline, whereas *smoppix* considers the background of all other molecules as null. The Venn diagram for the bivariate analysis is shown in Figure S26, with *spicyR* finding a lot of colocalization (see Figures S27 and S28 for most significant results), and smoppix more antilocalization. The gene found most significantly antilocalized by smoppix among the ones found colocalized by *spicyR* is shown in Figure S29; especially at time 10h there are indeed regions with only one of both genes expressed.

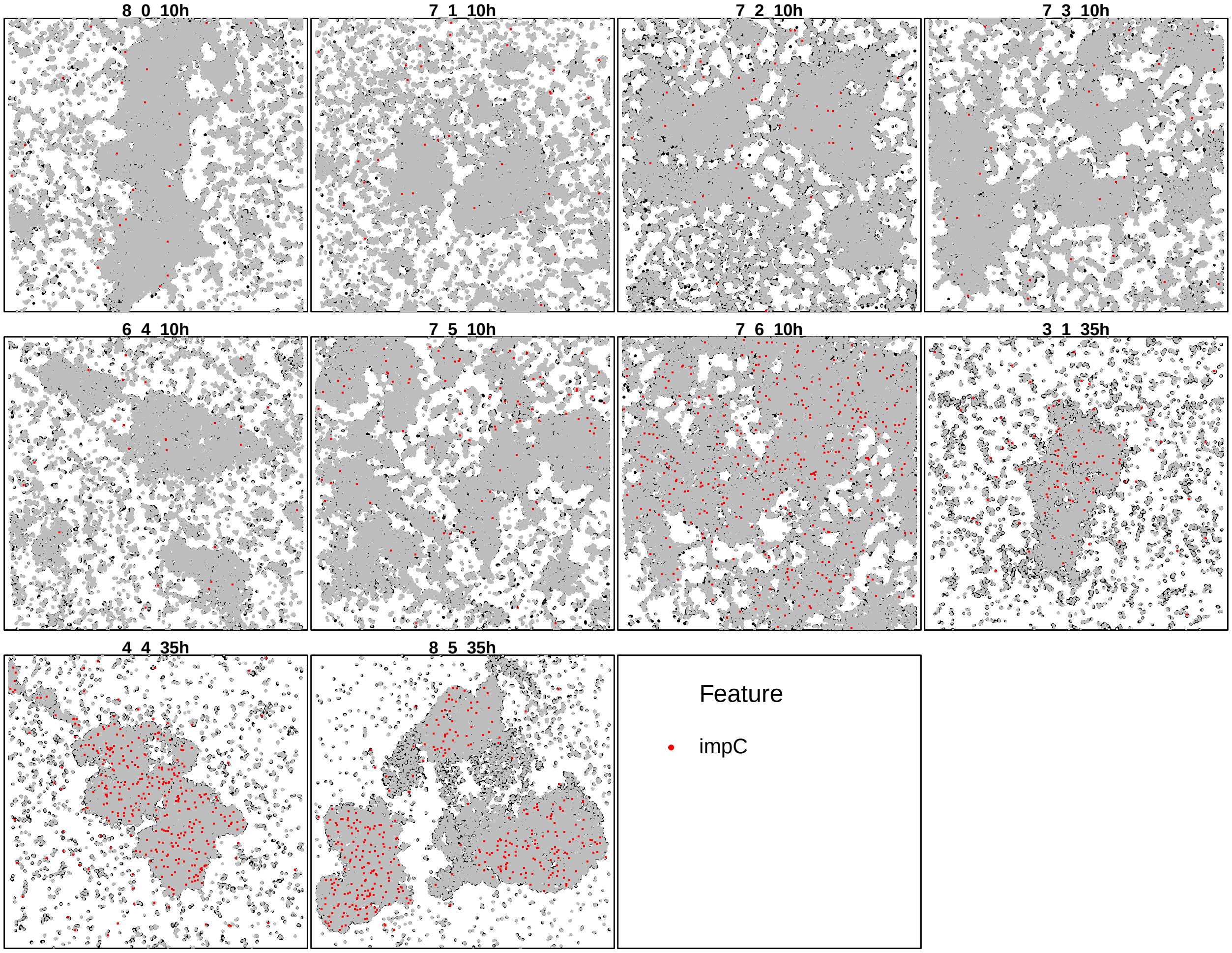

Figure S23: Most significantly aggregated gene according to *spicyR* (also significant in the *smoppix* analysis).

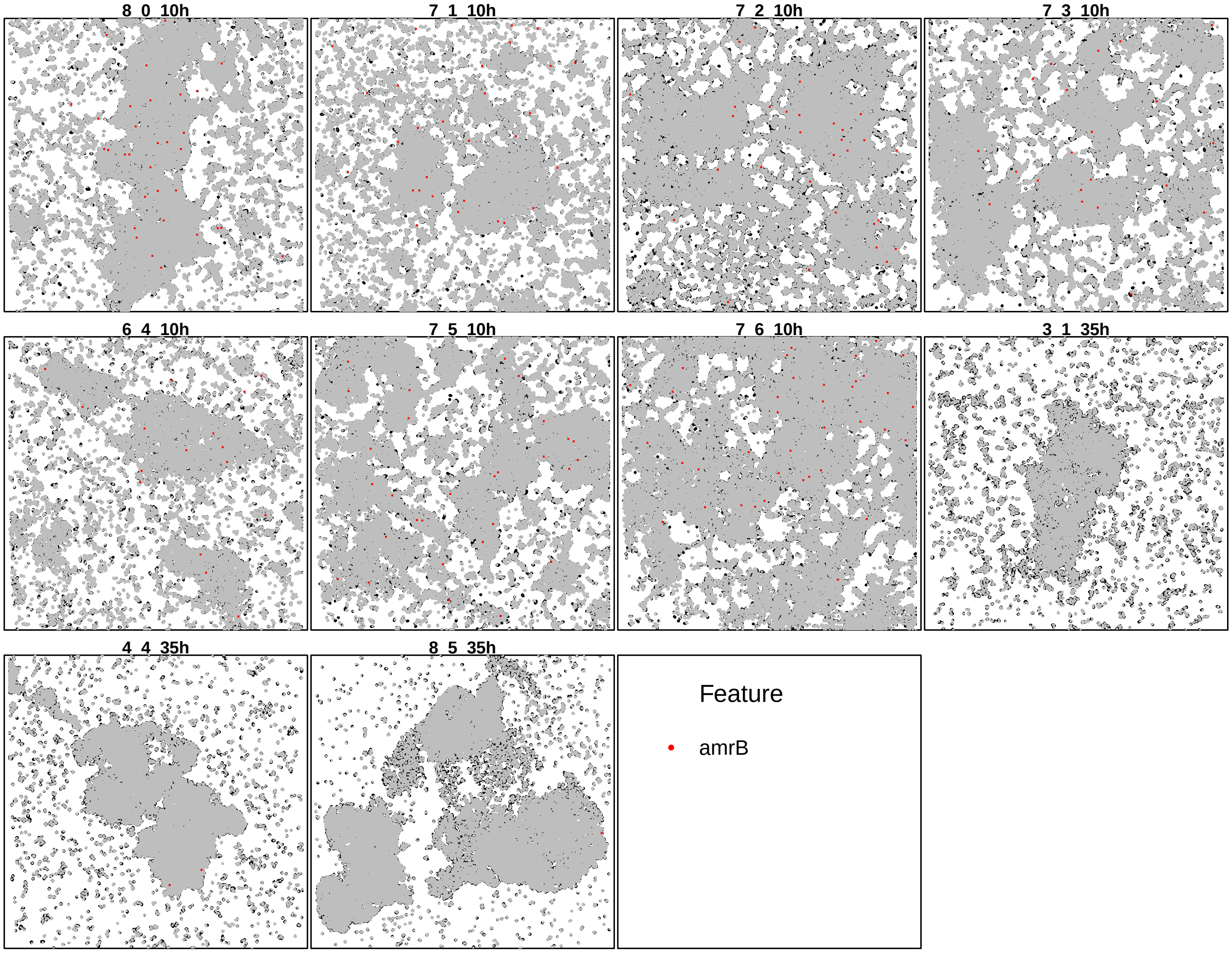

Figure S24: Most significantly regularly spaced gene according to *spicyR* (not significant for *smoppix*).

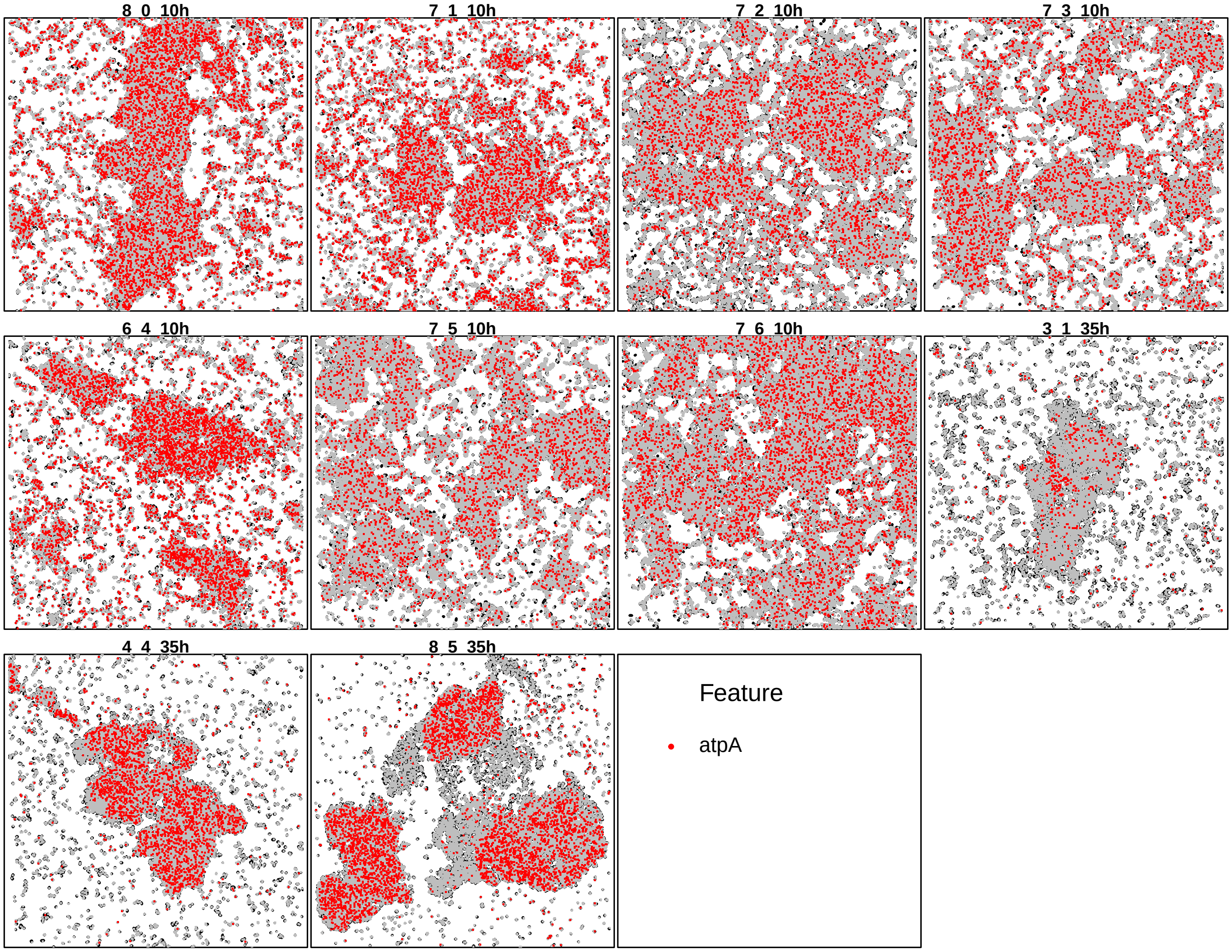

Figure S25: Gene found aggregated by *spicyR* but regularly spaced by *smoppix*

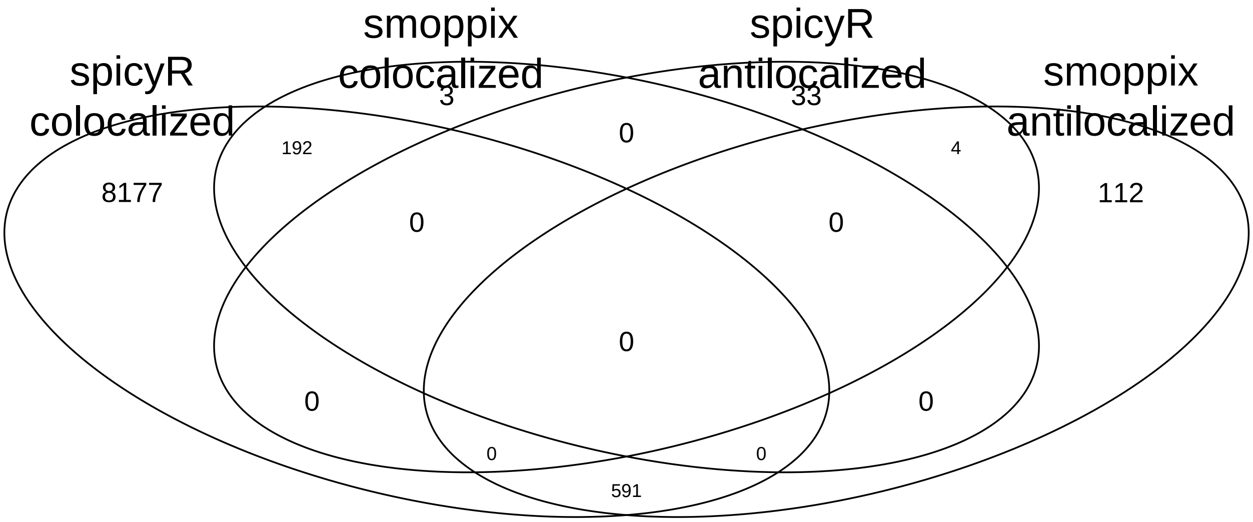

Figure S26: Venn diagram of overlap between *smoppix* and *spicyR* results for bivariate analysis.

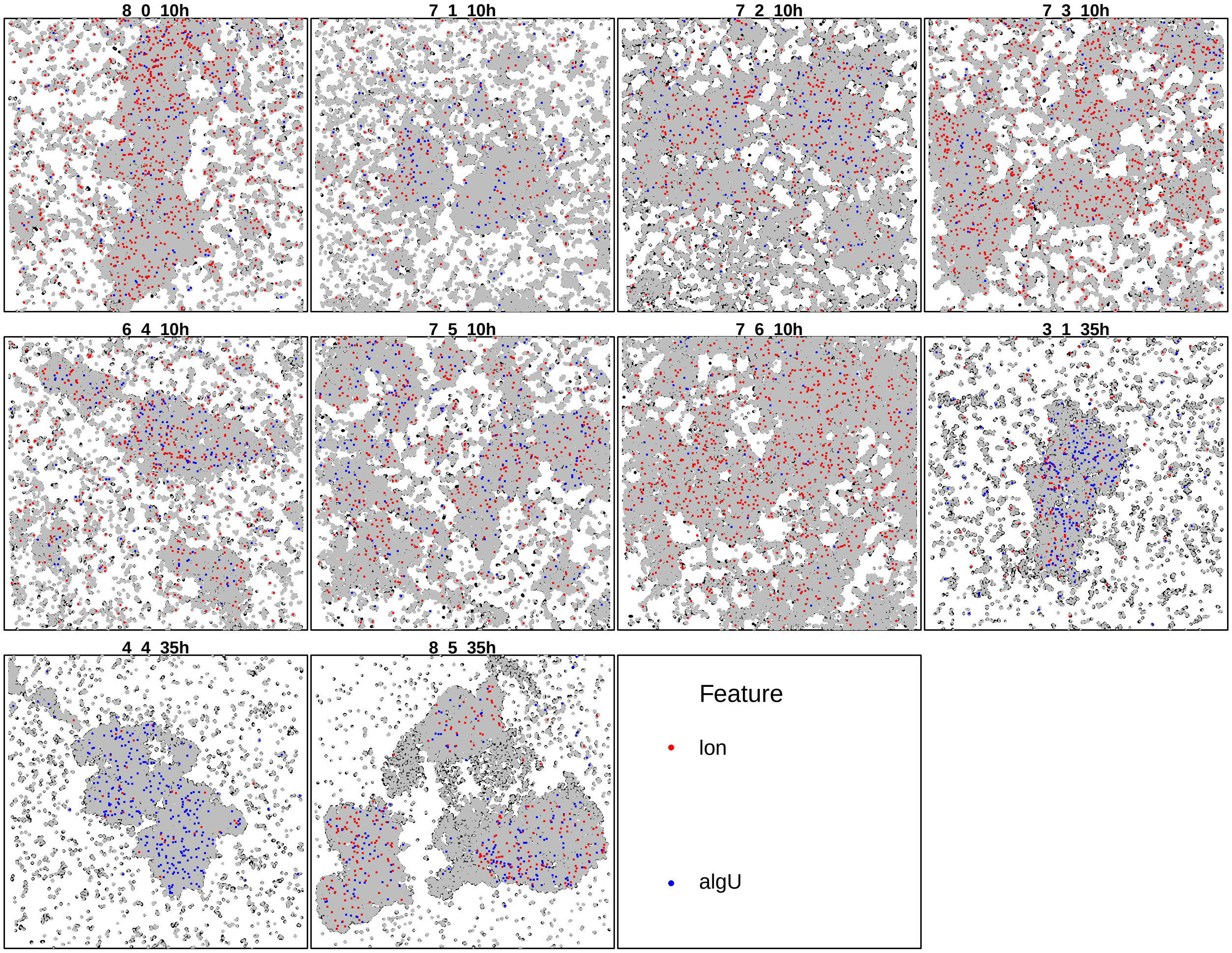

Figure S27: Most significantly colocalized gene pair according to *spicyR* (not significant for *smoppix*).

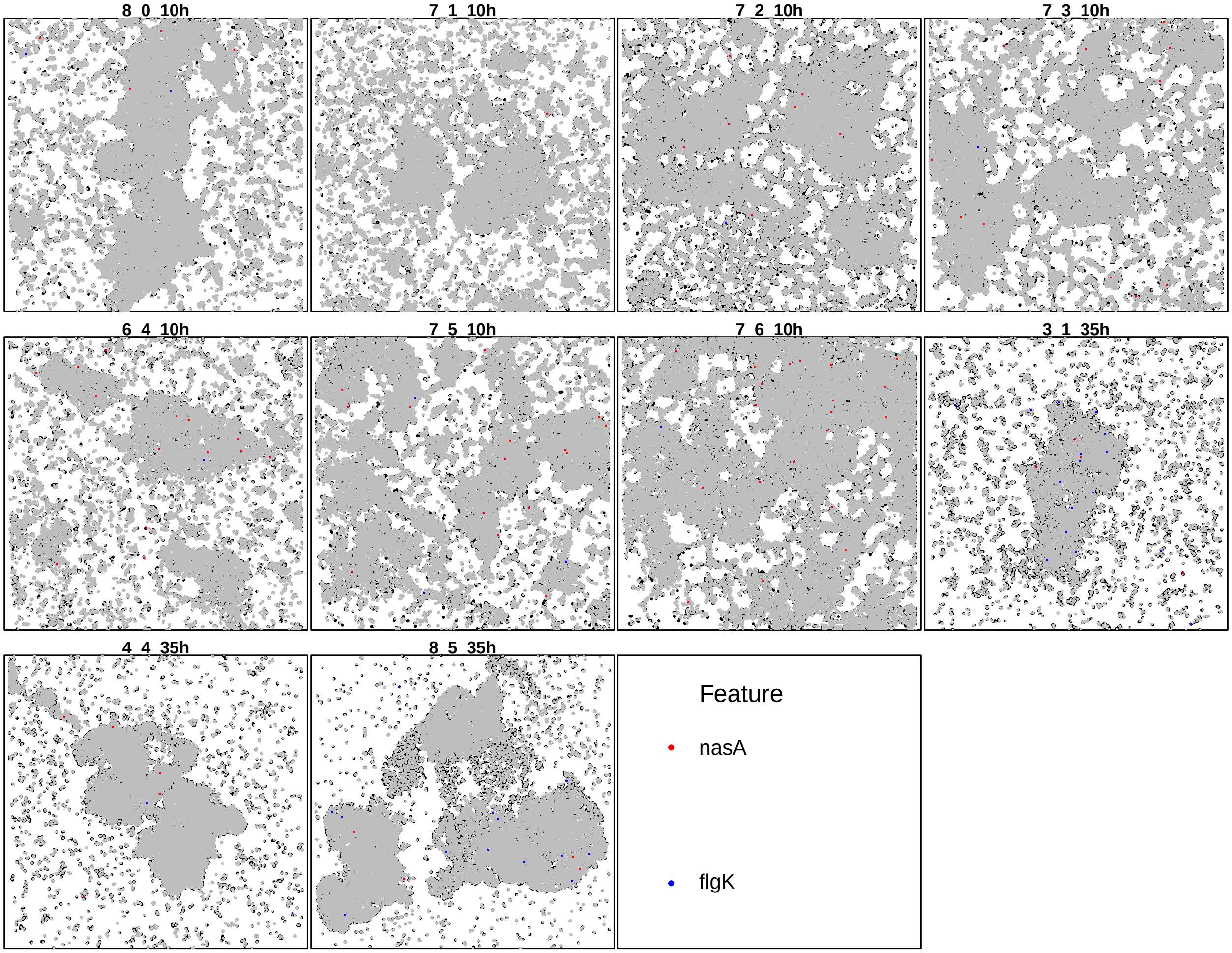

Figure S28: Most significantly antilocalized gene according to *spicyR* (not significant for *smoppix*).

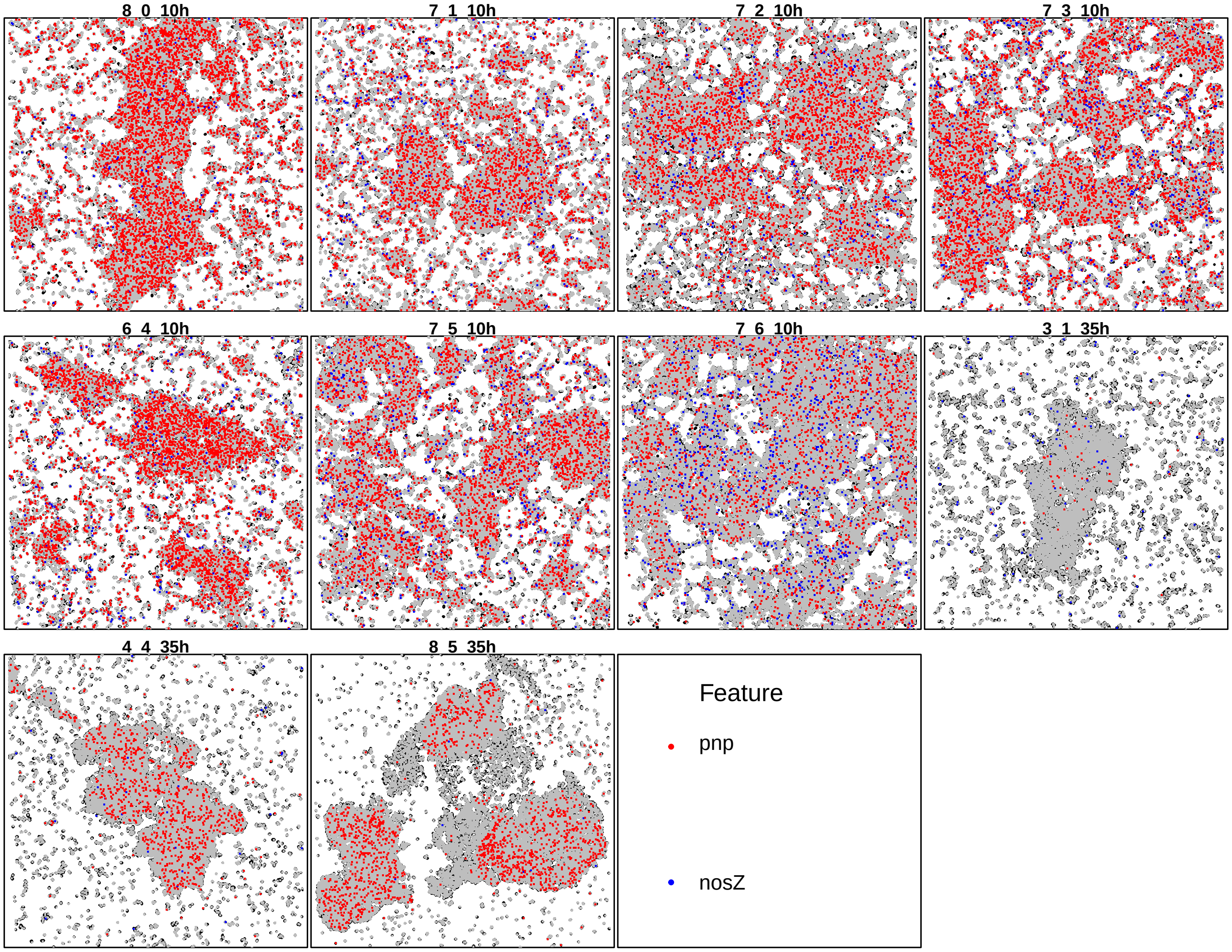

Figure S29: Most significantly antilocalized gene pair according to *smoppix* among the gene pairs colocalized according to *spicyR*.

#### 1.4 Probabilistic indices serve as interpretable predictors in tumor classification

We reanalysed a dataset on breast cancer sections from 33 patients [13], available through the *funkycells* R-package [14]. Locations of 16 different cell types were measured in a single section per patient. The tumors were categorized into two classes, “compartimentalised” and “mixed” by the authors of the original article [13]. A plot highlighting the tumor cells is shown in Figure 6 in the main text. In addition, the age of the patients was recorded, revealing an association between age and tumor type (Figure S30). There are also differences in cell type counts across the tumor types (Figure S31). Here we test whether PIs can distinguish between tumor types, and whether they can serve as interpretable predictors.

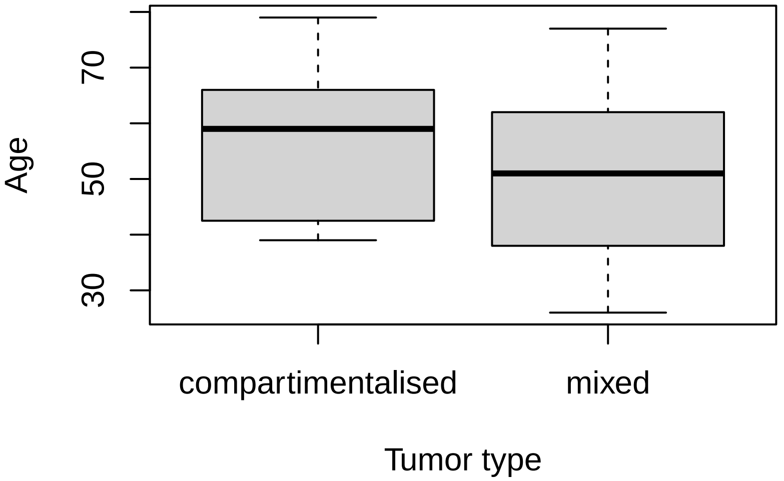

Figure S30: Boxplot of age (y-axis) as a function of tumor type (x-axis).

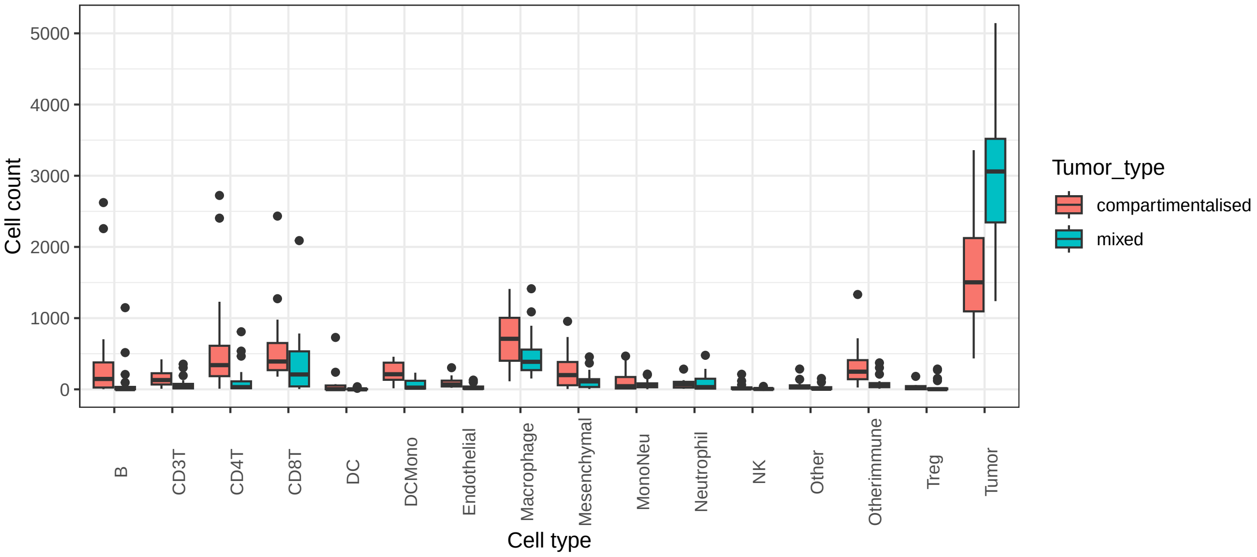

Figure S31: Boxplots of cell counts (y-axis) as a function of tumor type (colour) for different cell types (x-axis).

##### 1.4.1 Hypothesis tests

Significant differences between tumor types according to *smoppix* are shown in Table S3. The cell type pair most significantly more antilocalized in the "compartimentalised" type according to *smoppix* (CD4T–tumor) is shown in Figure S32. Significant differences between tumor types in neighbourhood enrichment tests are found in Table S4.

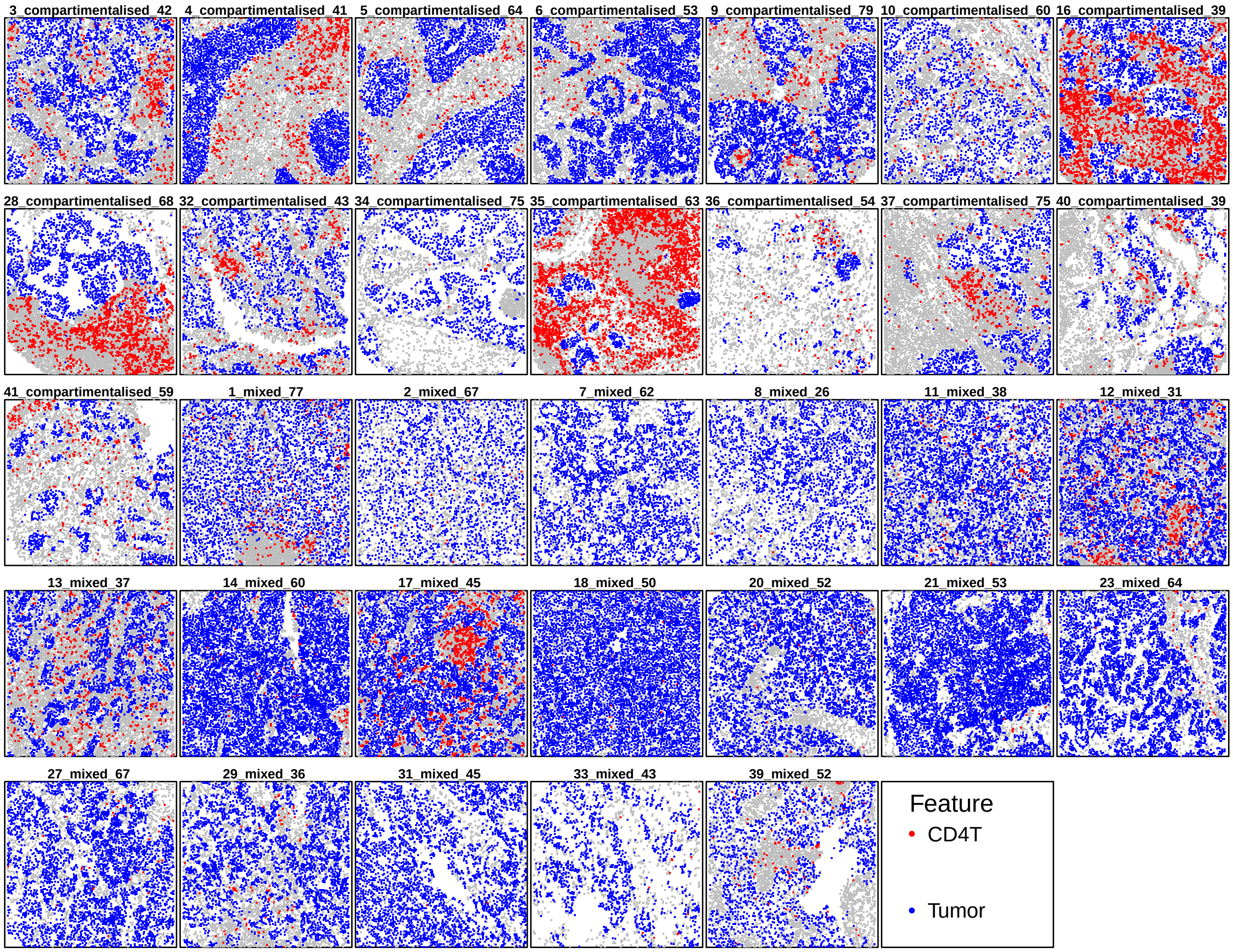

Figure S32: Cell type pair most significantly more antilocalized in type 'compartimentalised' than in type 'mixed' according to *smoppix*.

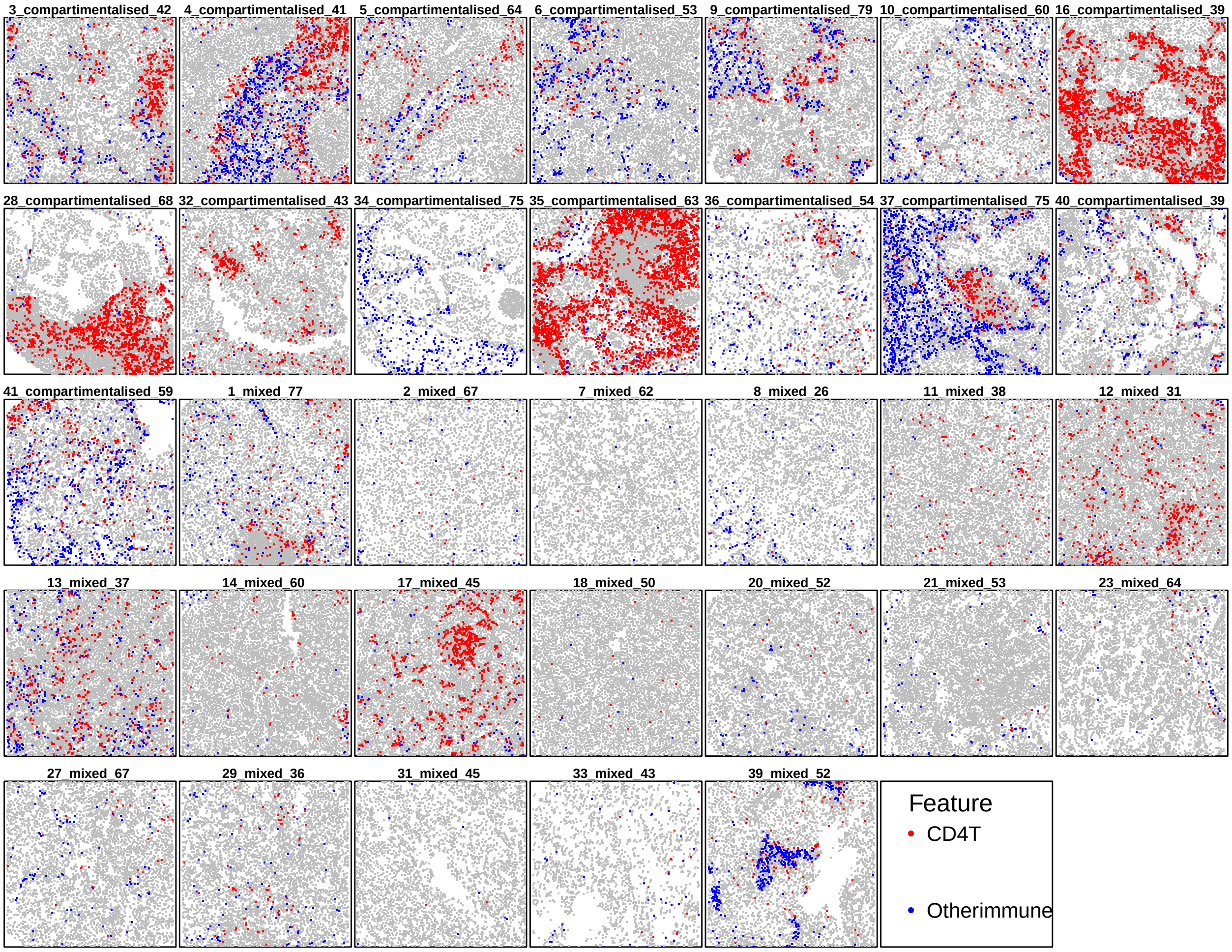

Figure S33: Scatterplot of the only cell type pair not involving tumor cells that is significantly more antilocalized in type ‘compartimentalised’ than in type ‘mixed’ according to *smoppix*.

| | Tumor type: compartimentalised | Tumor type: mixed | P-value | Adjusted p-value |
|---|---|---|---|---|
| Tumor | -0.100 | 0.100 | 1.0e-09 | 1.7e-08 |
| CD4T–Tumor | 0.088 | -0.088 | 1.7e-05 | 1.0e-03 |
| DCMono–Tumor | 0.067 | -0.067 | 1.7e-05 | 1.0e-03 |
| Otherimmune–Tumor | 0.077 | -0.077 | 2.6e-04 | 9.2e-03 |
| CD8T–Tumor | 0.070 | -0.070 | 3.1e-04 | 9.2e-03 |
| CD4T–Otherimmune | 0.048 | -0.048 | 1.1e-03 | 2.6e-02 |
| Macrophage–Tumor | 0.053 | -0.053 | 1.8e-03 | 3.6e-02 |
| B–Tumor | 0.083 | -0.083 | 2.1e-03 | 3.6e-02 |

Table S3: Table of univariate and bivariate localization patterns significantly different between tumor types according to *smoppix*. The first two columns show the coefficients of the tumor types in the sum coding scheme. For the bivariate patterns, the cell types are separated by hyphens.

| | Estimate | Standard error | P-value | Adjusted p-value |
|---|---|---|---|---|
| Tumor | -0.79 | 0.13 | 1.71e-06 | 2.74e-05 |
| Tumor–CD4T | 0.67 | 0.16 | 3.27e-04 | 3.92e-02 |
| Tumor–Otherimmune | 0.65 | 0.16 | 5.03e-04 | 4.03e-02 |
| CD4T–Tumor | 0.76 | 0.16 | 3.64e-05 | 8.73e-03 |
| Otherimmune–Tumor | 0.64 | 0.17 | 7.82e-04 | 4.69e-02 |

Table S4: Table of significant differences in univariate and bivariate neighbourhood enrichments between tumor types. The estimates shown are the average difference (the 'mixed' tumors minus the 'compartimentalised' tumors) in log-ratio of the proportion of molecules in the neighbourhood over overall in the image. For the bivariate results (cell type pairs separated by hyphens), the estimate refers to enrichment of the second cell type in the vicinity of the first. Positive estimates indicate more aggregation or colocalization in the mixed group, and vice versa for negative estimates.

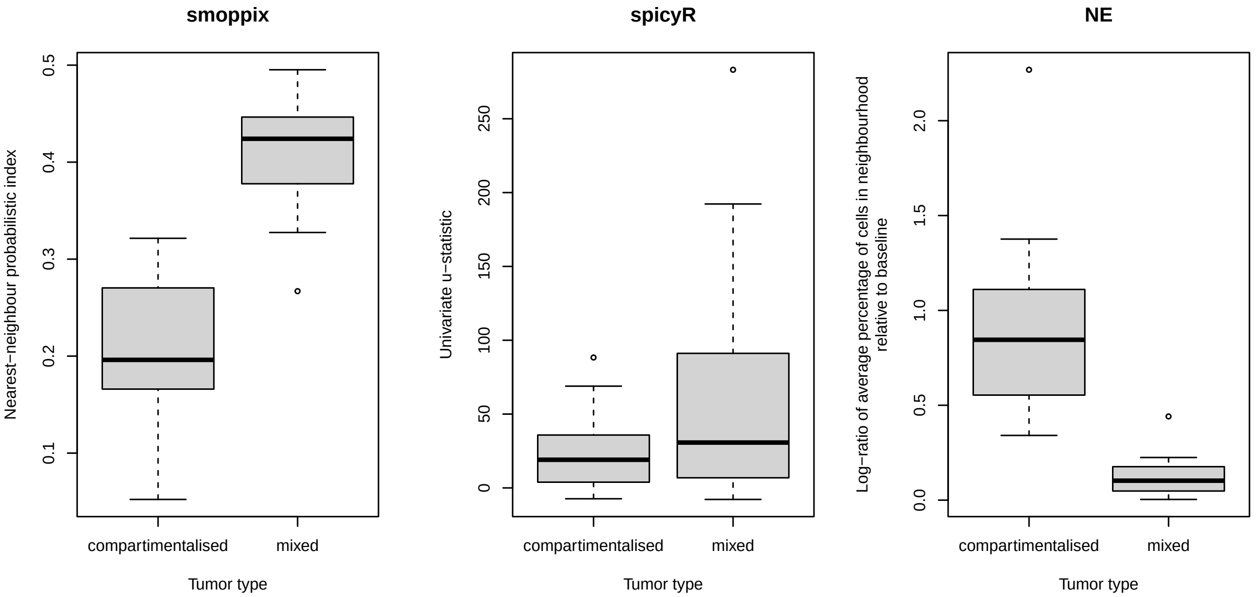

Figure S34: Boxplots of univariate nearest-neighbour distance PI by *smoppix* (left), u-statistic by *spicyR* (center) and NE log-ratio (right) (y-axis) as a function of tumor type (x-axis) for tumor cells. The difference is pronounced for *smoppix* and NE, which capture the aggregation of tumor cells as seen from Figure 6 in the main text. The difference in *spicyR*'s test statistic is less pronounced and even suggests stronger aggregation of tumor cells in the mixed class.

##### 1.4.2 Prediction models

Next we investigated whether the inclusion of PI's as predictor of the tumor type, on top of age and cell type counts, improves predictive accuracy. We use a penalized linear regression model (LASSO) [15] implemented in the *glmnet* package [16] as prediction model, and assess its predictive accuracy (defined as the proportion of correctly classified tumors) through 20-fold cross-validation. The penalty parameter $\lambda$ is tuned through an inner cross-validation loop. The following sets of predictors are used in the model: patient age is always included as predictor, and cell type counts or PIs or both are included in additional models. When only age is included, a regular generalized linear model is used for prediction, without regularization. As feature selection procedure for the PIs, *smoppix* is applied to the observations in the training folds with tumor type and age as fixed effects, and PIs significantly associated with tumor type are retained as predictors for the LASSO model; their missing values are set to 0.5. The splitting into folds was repeated 200 times and out-of-sample accuracy and its standard error were estimated following Bates, Hastie, and Tibshirani [17]. The results are shown in Table 2 in the main text, and confirm that spatial localization patterns add information on tumor type on top of cell counts and patient age. A similar analysis was carried out using NE log-ratios as predictors, yielding similar results to the PIs as shown in Table S5. Since *spicyR* does not declare any cell types significant, no cell type (pairs) were filtered out and all u-statistics are fed as predictors to the LASSO model. Yet inclusion of these u-statistics deteriorates the predictive performance (see Table S5).

| | Age + NE log-ratio | Age + cell count + NE log-ratio | Age + spicyR u | Age + cell count + spicyR u |
|---|---|---|---|---|
| Accuracy | 0.92 | 0.91 | 0.36 | 0.71 |
| SE | 0.08 | 0.06 | 0.09 | 0.08 |

Table S5: Average predictive accuracy over 200 repeats of the split into cross-validation folds for LASSO regression models with different sets of predictors (column names) derived from NE and *spicyR*. Standard errors (SE) of the accuracy estimates are shown in the second row. For *spicyR*, no preliminary feature filtering through hypothesis testing was applied.

### 2 Benchmarking

#### 2.1 Calibration: the p-value distribution under gene label permutation

To assess calibration of the statistical tests, we repeatedly permute the gene labels in the *S. moellendorfii* root [1] and bacterial biofilm [12] datasets and then apply the regular *smoppix*, *spicyR* and NE analyses (NE could not be applied to the biofilm dataset for computational reasons). The resulting p-value distributions are shown in Figures S35-S36 and suggest *smoppix* to be a well-calibrated test. The slight peak of low p-values for the univariate nearest-neighbour distances in Figure S36 is mainly caused by highly expressed genes and is likely a result of the compositional effect discussed in Supplementary Section 3.3. This effect decreases as the dimensionality of the data decreases. The qq-plots for *spicyR* and NE reveal strong and moderate departures from uniformity, respectively.

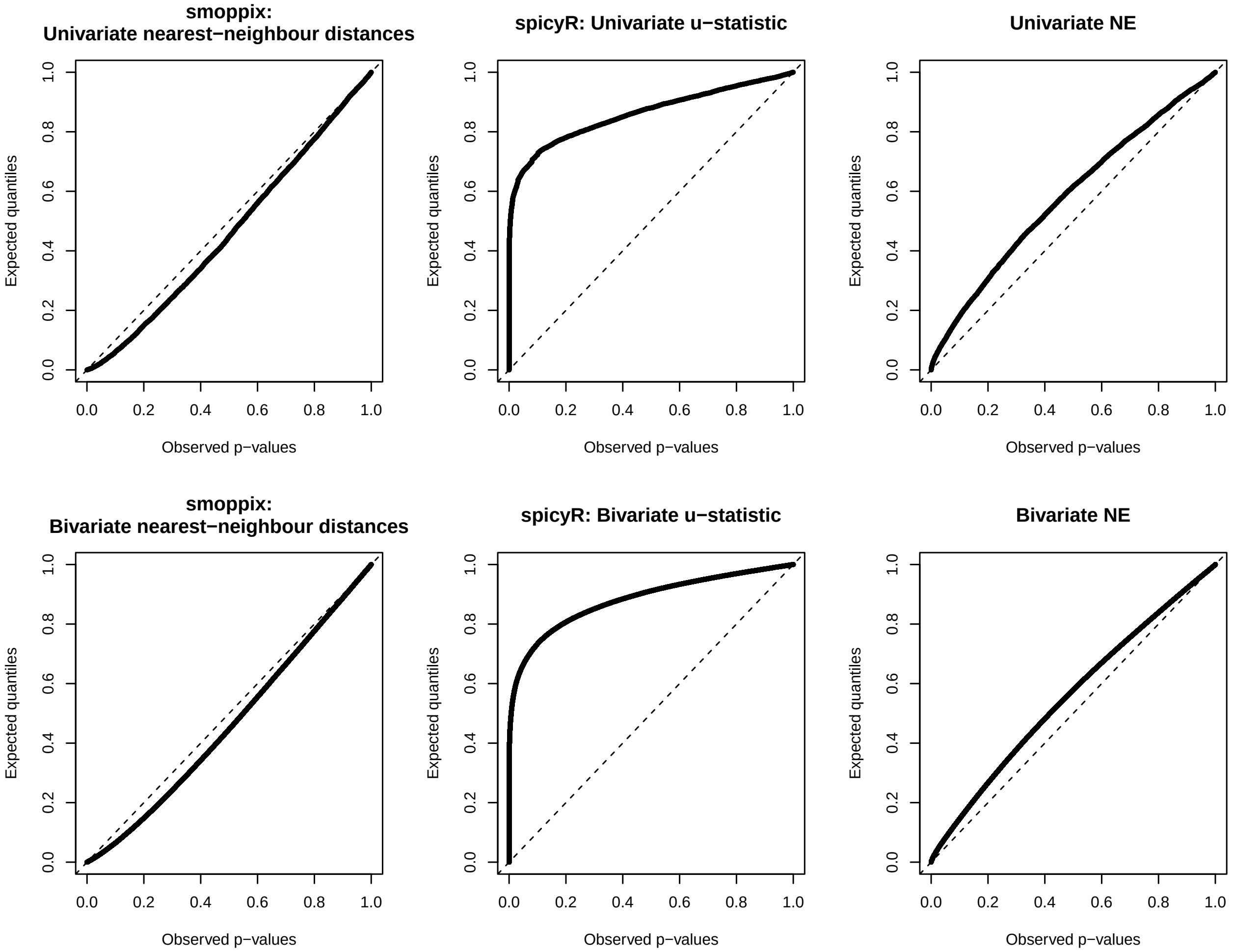

Figure S35: qq-plots versus standard uniform quantiles of *smoppix*, *spicyR* and NE p-values after permutation of the *S. moellendorfii* root data of Yang et al. [1]. The method and test statistic are shown in the title.

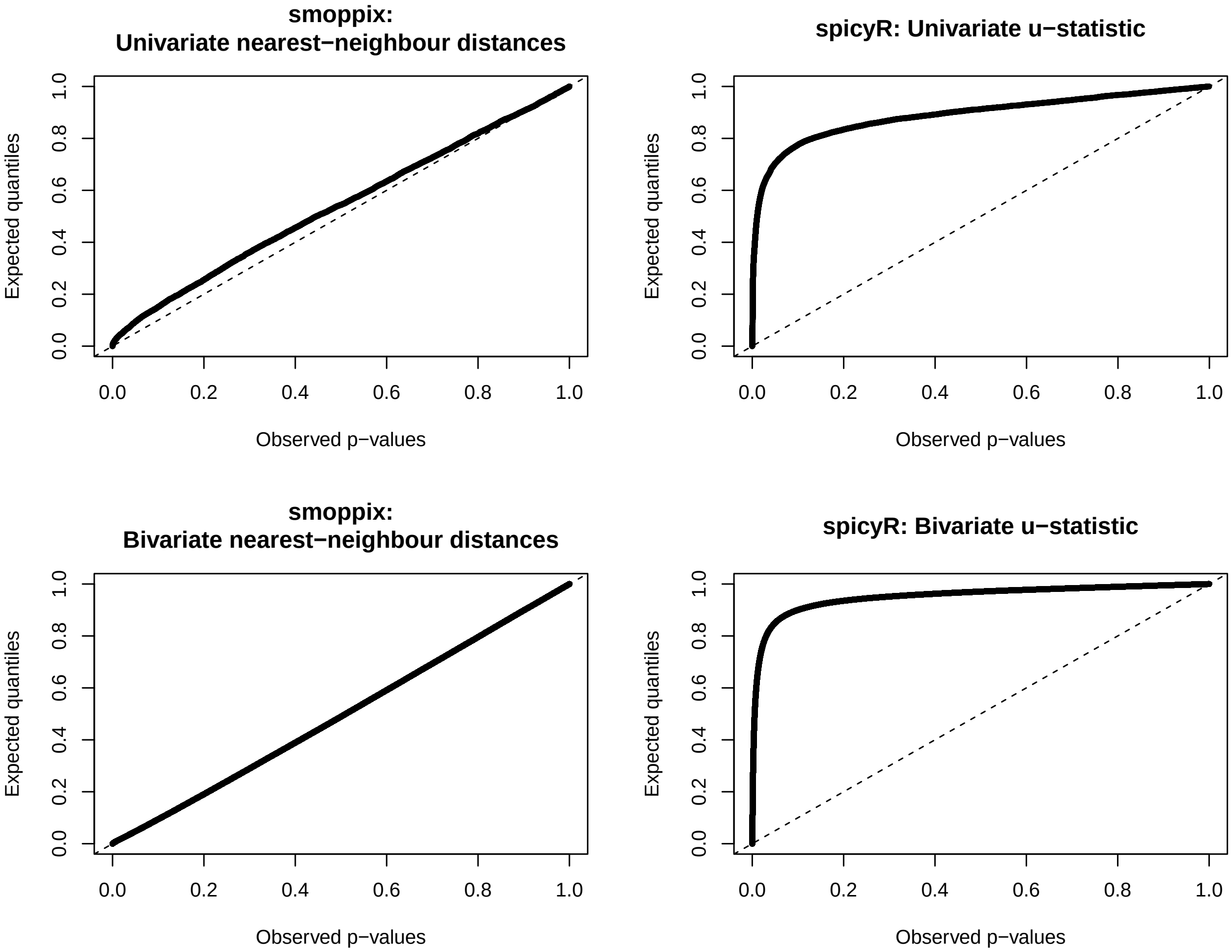

Figure S36: qq-plots versus standard uniform quantiles of *smoppix* and *spicyR* p-values after permutation of the gene labels of the bacterial biofilm data by Dar et al. [12]. The NE method was too computationally intensive to be run on this dataset.

#### 2.2 Invariance under subsampling

We investigated the sensitivity to density differences through a series of subsampling experiments for *smoppix*, *spicyR* and NE. The molecules of the point patterns from the *S. moellendorfii* root [1] and mouse fibroblast [10] datasets were randomly subsampled to proportions running from 0.9 to 0.4 by decrements of 0.1. Measures of aggregation and colocalization were calculated for the subsampled datasets of the first dataset, and PIs for distance to edge and centroid for the datasets subsampled from the second dataset. For the univariate PIs, the 10 most expressed genes were used, for the bivariate PIs the pairs of 11th-20th most expressed genes were paired with the 21st-30th most expressed genes. Choosing non-overlapping sets of genes guarantees independence in the coming regression analysis. The estimated PIs, *spicyR*'s u-statistics and NE log-ratios are plotted as a function of subsampling proportion in Figures S37-S43 for a subset of genes. To formally test sensitivity to subsampling, we fit linear models with the measure of spatial patterning (either PI, u-statistic or log-ratio) as outcome as a function of subsampling proportion, day and root-gene (pair) combination for the *S. moellendorfii* dataset, or as a function of subsampling proportion and experiment-fov-gene combination for the mouse fibroblast dataset. The estimated slopes and associated p-values are shown in Table S6. The univariate nearest-neighbour PI is slightly sensitive to density changes, as observed and background nearest-neighbour distributions do not change in the same way under subsampling. In addition, both the univariate and bivariate u-statistics are sensitive to density changes. Yet the changes in the statistics with subsampling are rather modest compared to the overall variability of the test statistics in the datasets, as quantified through the interquartile ranges (IQRs). All other measures are invariant to subsampling.

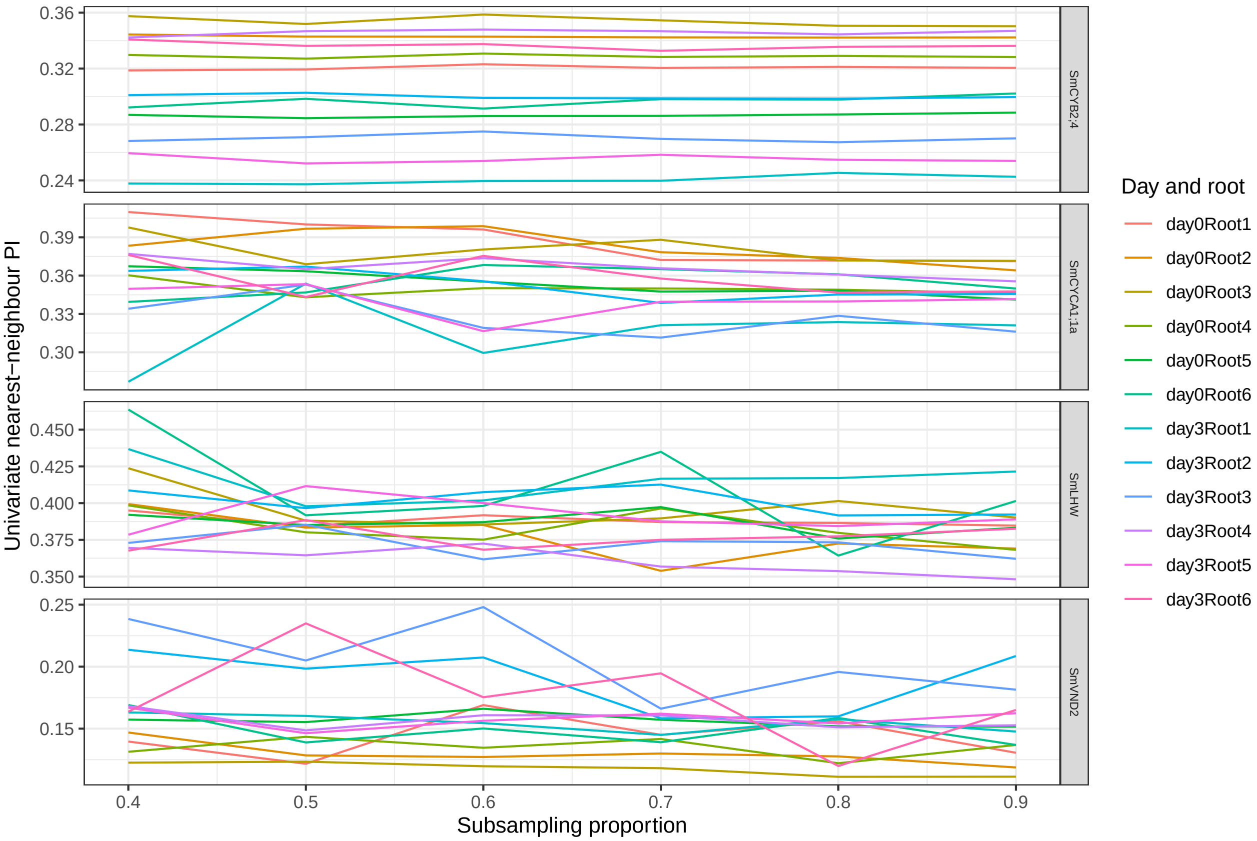

Figure S37: Estimated univariate nearest-neighbour PIs (y-axis) of the 4 most highly expressed genes (rows) as a function of subsampling proportion (x-axis) for the *S. moellendorfii* root data by Yang et al. [1], averaged over sections. Separate lines are drawn per root (colours).

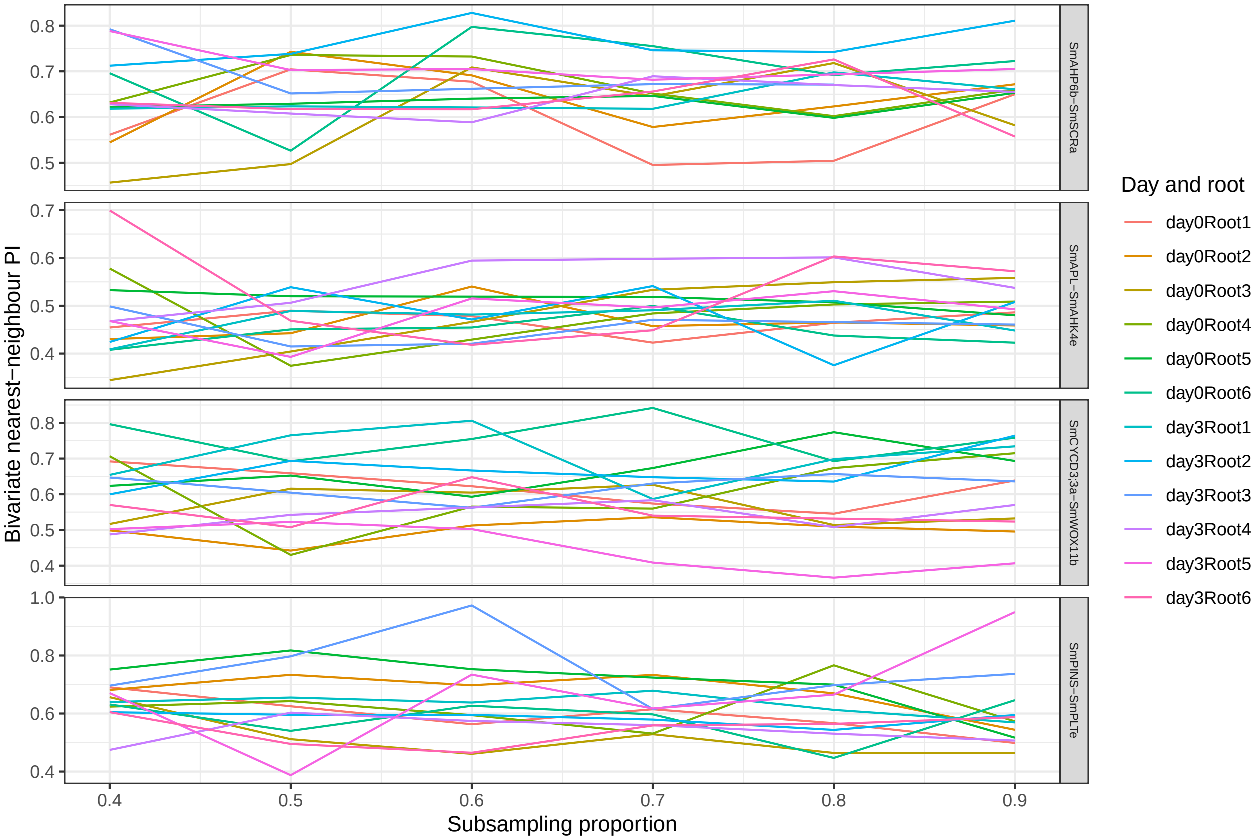

Figure S38: Estimated bivariate nearest-neighbour PIs (y-axis) of the 4 most highly expressed gene pairs (rows) as a function of subsampling proportion (x-axis) for the *S. moellendorfii* root data by Yang et al. [1], averaged over sections. Separate lines are drawn per root (colours).

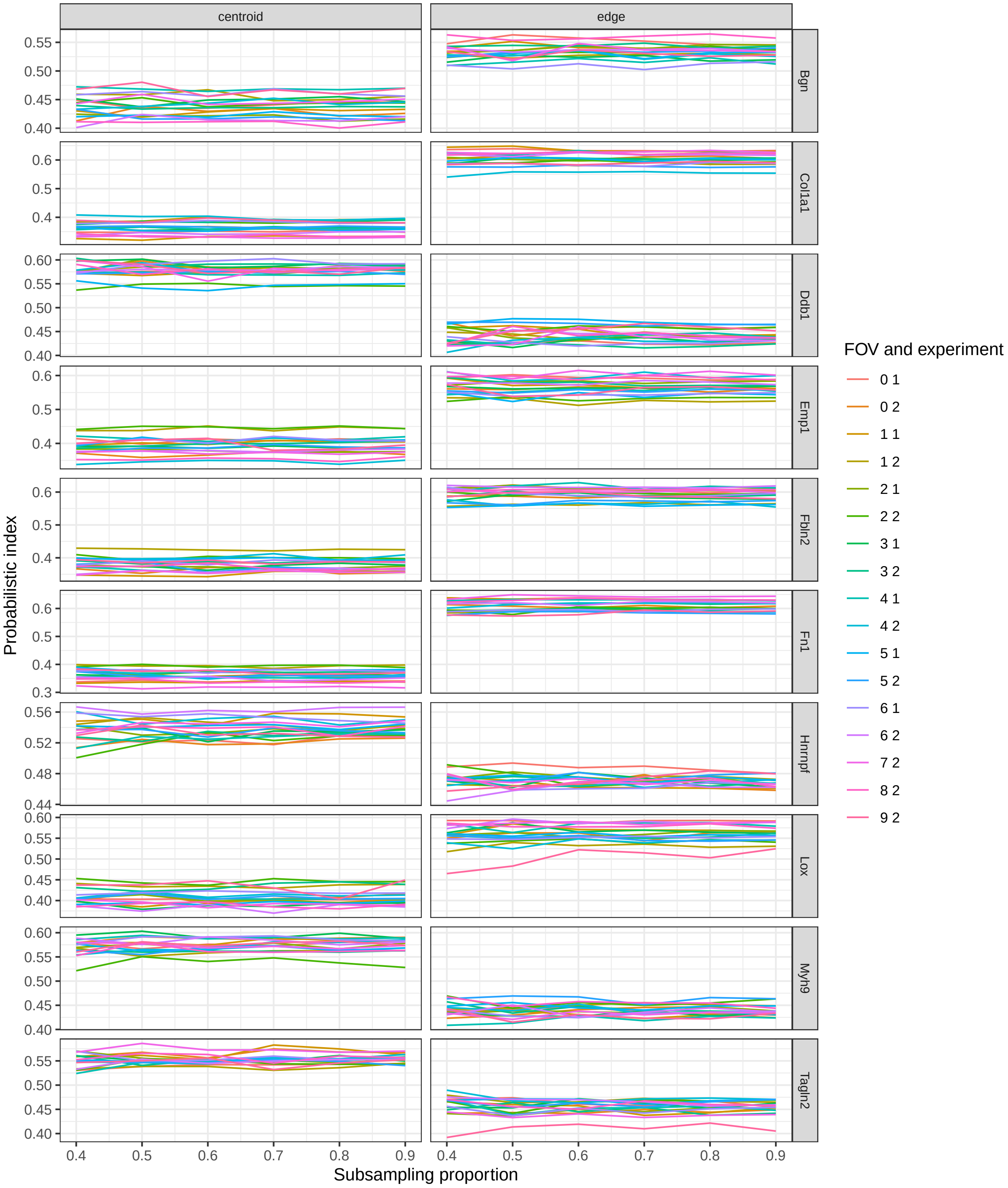

Figure S39: Estimated univariate intracellular PIs (y-axis) for vicinity to centroid or edge (columns) of the 10 most highly expressed genes (rows) as a function of subsampling proportion (x-axis) for the mouse fibroblast data by Eng et al. [10], averaged over cells within fields of view. Separate lines are drawn per field of view and experiment (colours).

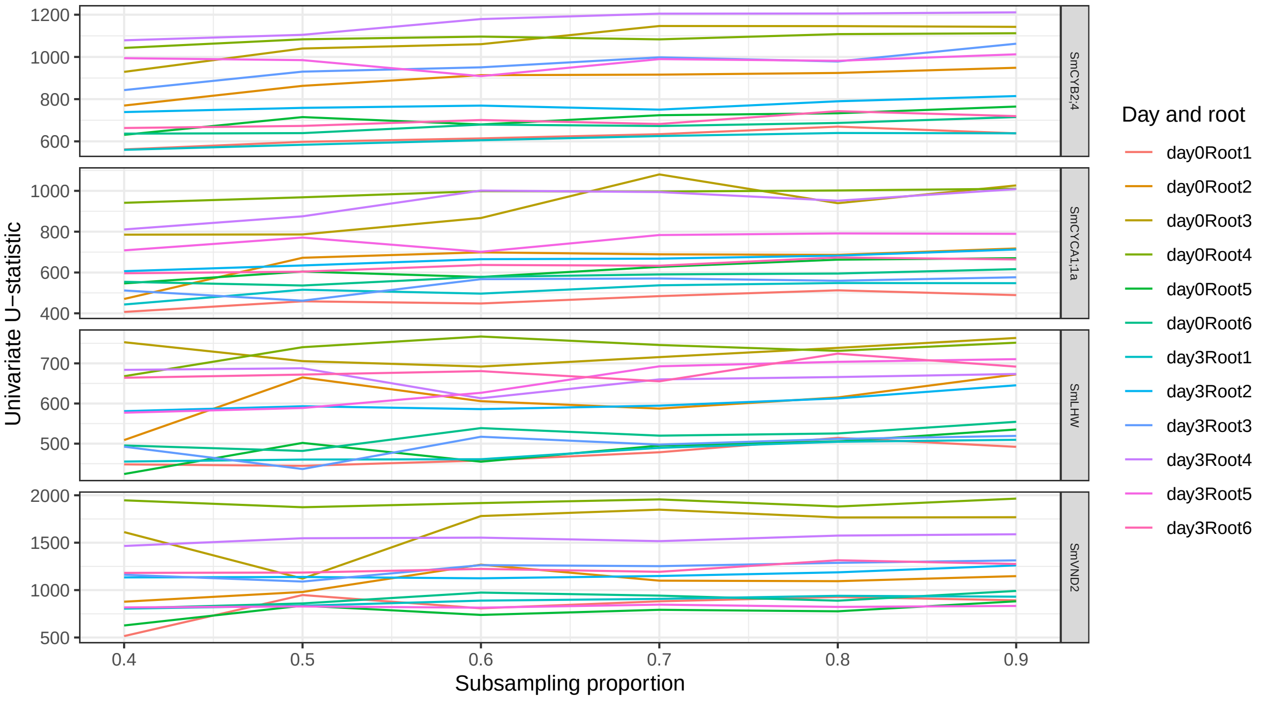

Figure S40: Estimated univariate u-statistic (y-axis) of the4 most highly expressed genes (rows) as a function of subsampling proportion (x-axis) for the *S. moellendorfii* root data by Yang et al. [1], averaged over sections. Separate lines are drawn per root (colours).

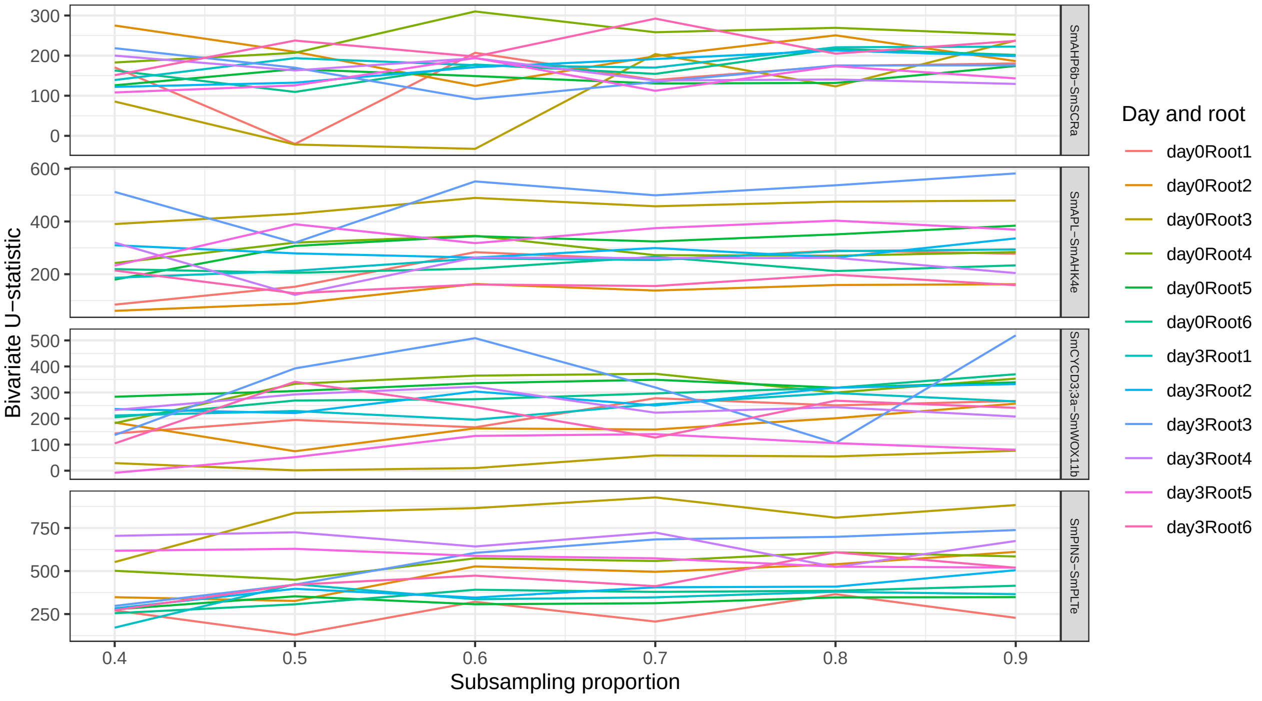

Figure S41: Estimated bivariate u-statistic (y-axis) of the 4 selected gene pairs (rows) as a function of subsampling proportion (x-axis) for the *S. moellendorfii* root data by Yang et al. [1], averaged over sections. Separate lines are drawn per root (colours).

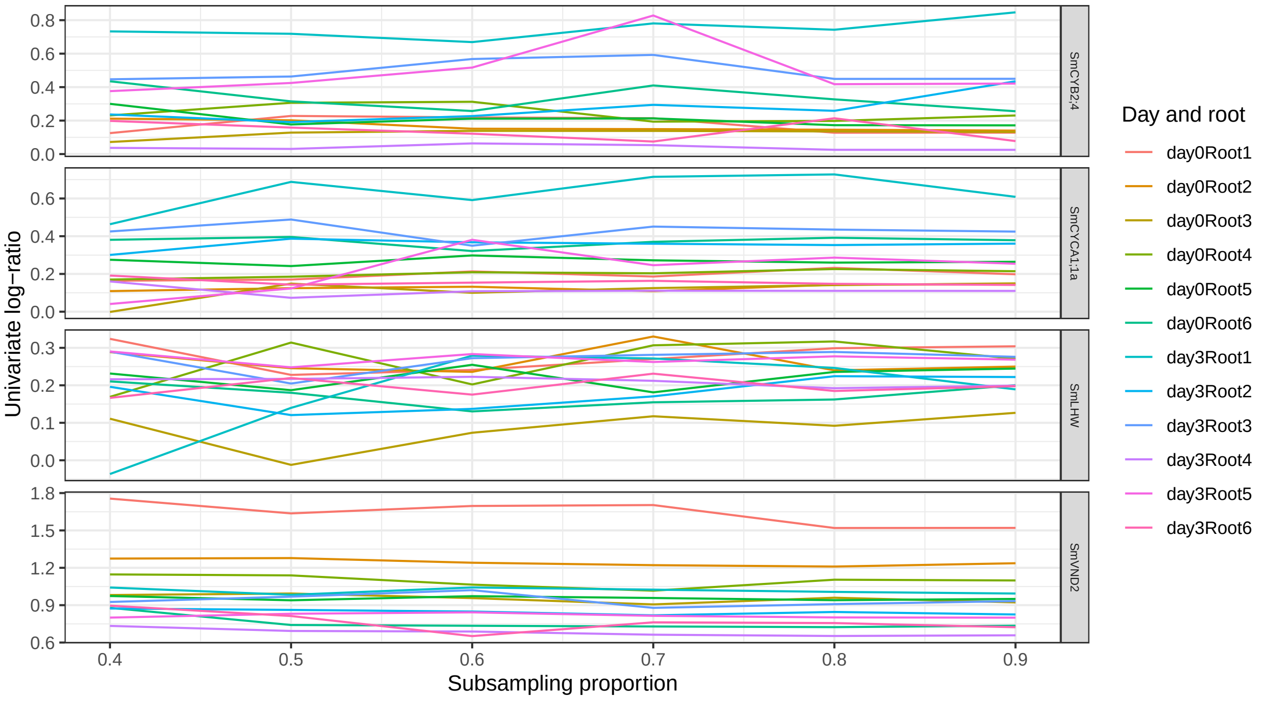

Figure S42: Estimated log-ratio of univariate neighbourhood enrichment by NE (y-axis) of the 4 most highly expressed genes (rows) as a function of subsampling proportion (x-axis) for the *S. moellendorfii* root data by Yang et al. [1], averaged over sections. Separate lines are drawn per root (colours).

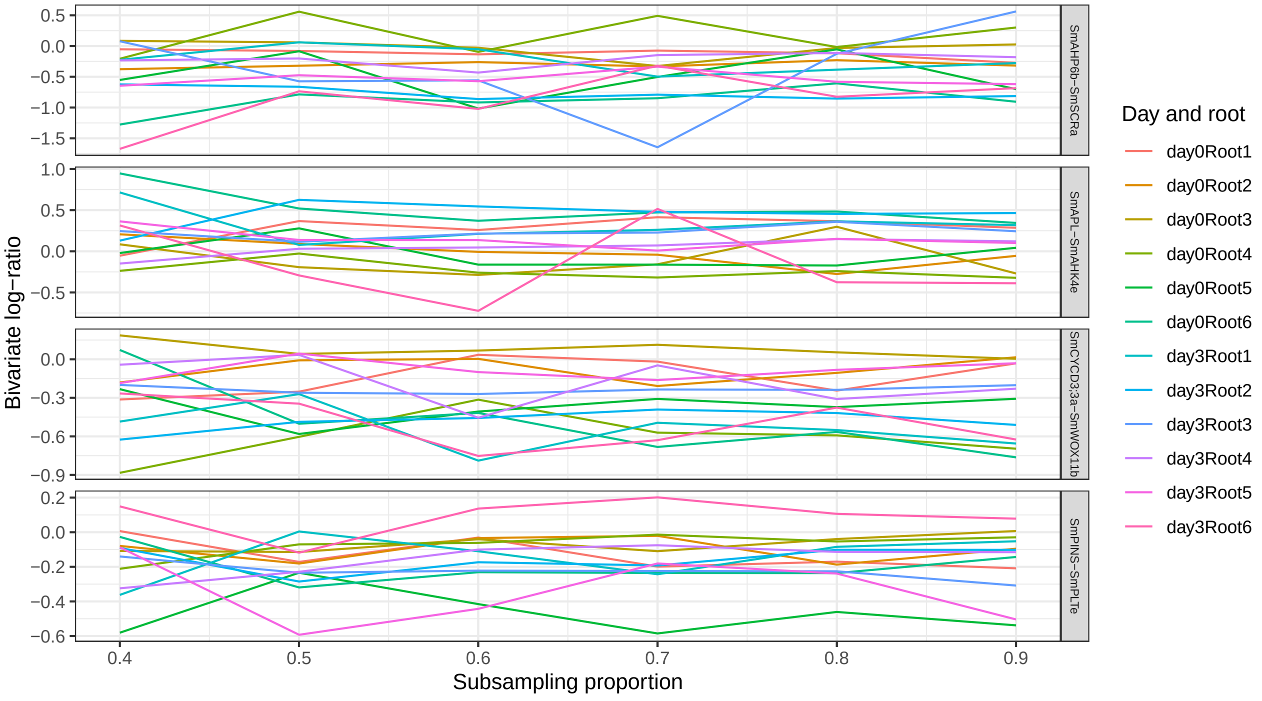

Figure S43: Estimated log-ratio of bivariate neighbourhood enrichment by NE (y-axis) of the 4 selected gene pairs (rows) as a function of subsampling proportion (x-axis) for the *S. moellendorfii* root data by Yang et al. [1], averaged over sections. Separate lines are drawn per root (colours).

| | Slope | P-value | IQR |
|---|---|---|---|
| Univariate PI | -0.00283 | 2.8e-06 | 0.18 |
| Bivariate PI | -0.00010 | 9.5e-01 | 0.27 |
| Centroid PI | 0.00016 | 2.0e-01 | 0.46 |
| Edge PI | 0.00005 | 6.8e-01 | 0.41 |
| Univariate NE | 0.0029 | 4.8e-01 | 1.63 |
| Bivariate NE | 0.0050 | 4.4e-01 | 1.21 |
| Univariate u-statistic | 20.6 | 5.4e-16 | 590.49 |
| Bivariate u-statistic | 18.1 | 3.2e-16 | 585.39 |

Table S6: Estimated slopes of association measure as a function of subsampling proportion and associated p-values. The centroid and edge PI are estimated on the dataset by Eng et al. [10], all others on the dataset by Yang et al. [1]. The slope is the expected change in the spatial measure associated with a 10% increase in subsampling proportion, IQR is the interquartile range of the spatial measure of the entire real dataset. NE: neighbourhood enrichment

#### 2.3 Time and memory benchmark

Timings and RAM usage on an Intel Core i5-11400H 2.70GHz processor were investigated for *smoppix*, *spicyR* and NE for increasing numbers of molecules and genes on synthetic point patterns. The point patterns were generated under CSR on a square area of 10 × 10, the neighbourhood size for NE was set at 1. The total number of molecules was varied between 500, 1,000, 2,000 and 3,000 and the number of features between 10, 60 and 100. Per parameter setting, 10 datasets were generated and runtimes and peak RAM usage were averaged over these repeats. Tests for univariate as well as bivariate patterns were run. No multithreading was used for any method (*smoppix* and *spicyR* natively allow for multithreading, NE is easily adapted for it). NE calculation with *SPIAT* crashed when applied to 100 features, so it was only run for fewer features. Results are shown in Figure S44. NE becomes very slow as th number of features increases, despite calculating the enrichment using SPIAT's *average_percentage_of_cells_within_radius* function, which calls the *frNN* function from the *dbscan* package that relies on a custom C-function.

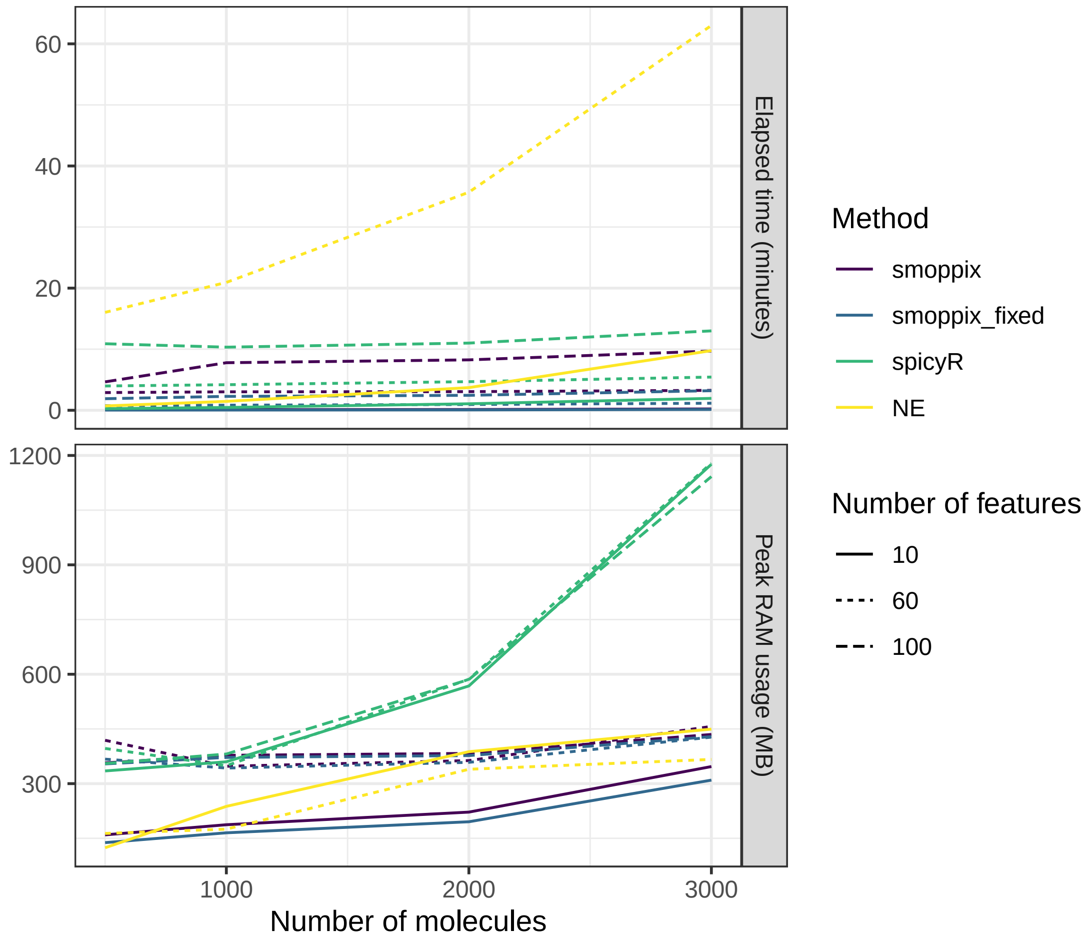

Figure S44: Average timings in minutes and peak RAM usage in megabytes (MB) over 10 repeats for different methods for testing for aggregation and colocalization. smoppix_fixed indicates that fixed-effects models rather than mixed-effects models were used, to illustrate the major computational share of fitting the linear mixed models after the PIs have been estimated.

### 3 Details of the *smoppix* method

#### 3.1 The negative hypergeometric distribution

The probabilistic index is defined as the evaluation of the distribution function under the null in the observed distances. In the main text, we argue that the PI of the nearest-neighbour distances under the background null can be found exactly using the negative hypergeometric distribution. According to this distribution, the number of attempts $x$ needed to reach $q$ successes when sampling without replacement from a set of $w = N - 1$ units with $k = N_g - 1$ successes has probability mass function [18]:

$$Pr(X = x) = \frac{\binom{x+q-1}{x}\binom{w-q-x}{k-x}}{\binom{w}{k}}, \tag{2}$$

and distribution function $H(x) = Pr(X \leq x) = \sum_{k=0}^{x} Pr(X = k)$. Setting $q = 1$, (2) assigns a probability to each distance $d_{(x)}$ (corresponding to the distance to $i$ with rank $x$) of being the smallest among the set of $N_g - 1$ included distances under the background null. In view of the definition of the PI in equation (6) in the main text, and the occurrence of ties, we define the following functions based on the set of all distances to a given molecule:

$$\begin{aligned} F_{all}(D) &= (N-1)^{-1} \sum_{i=1}^{N-1} I(D \leq d_{(i)}) \\ R_{all}(D) &= (N-1)^{-1} \sum_{i=1}^{N-1} I(D < d_{(i)}). \end{aligned} \tag{3}$$

Define $t_i = (N-1)F_{all}(d_{(i)})$ and $r_i = (N-1)R_{all}(d_{(i)})$. The PI can then be estimated as

$$\begin{aligned} \widehat{PI}_g &= \frac{\sum_{i=1}^{N_g-1} \left[H(r_i) + \frac{(H(t_i) - H(r_i))}{2}\right]}{(N_g - 1)} \\ &= \frac{\sum_{i=1}^{N_g-1} [H(r_i) + H(t_i)]}{2(N_g - 1)}, \end{aligned} \tag{4}$$

without having to do permutations. The functions in (3) can be computation and memory intensive to obtain when the number of molecules is large, as $(N - 1)$ distances need to be calculated per molecule $i$. In that case, approximations of $\tilde{F}_{all}$ and $\tilde{R}_{all}$ based on a random subset of $S$ out of $N - 1$ distances are also sufficient. The approximate ranks $\tilde{r}_i = \lfloor S\tilde{R}_{all}(d_i) \rceil$ and $\tilde{t}_i = \lfloor S\tilde{F}_{all}(d_i) \rceil$, with $\lfloor . \rceil$ meaning rounding to the nearest integer, are then plugged into (4). In the spirit of Phipson and Smyth [19], who propose a correction to prevent permutation p-values from becoming exactly zero, all $\tilde{t}_i$'s equal to 0 are set to 1. $\tilde{t}_i = 0$ can occur when the observed nearest-neighbour distance is excluded from $S$. Also when the null distribution and corresponding functions $R_{all}$ and $F_{all}$ are obtained from Monte-Carlo simulation under CSR, these approximate ranks can be used.

We confirm the exact enumeration of the PI null distribution using the negative hypergeometric distribution empirically using Monte-Carlo simulation. For a randomly drawn subset of size 100 out of a set of $10^4$ distances, the distribution of the nearest-neighbour distance (i.e the smallest observation of the subset) is found through permutation, so taking 100,000 other samples of size 100 and finding their minimum. This is compared to our analytical solution using the negative hypergeometric distribution, as implemented in *smoppix*. The resulting densities are similar (see Figure S45), with the exact negative hypergeometric distribution being faster.

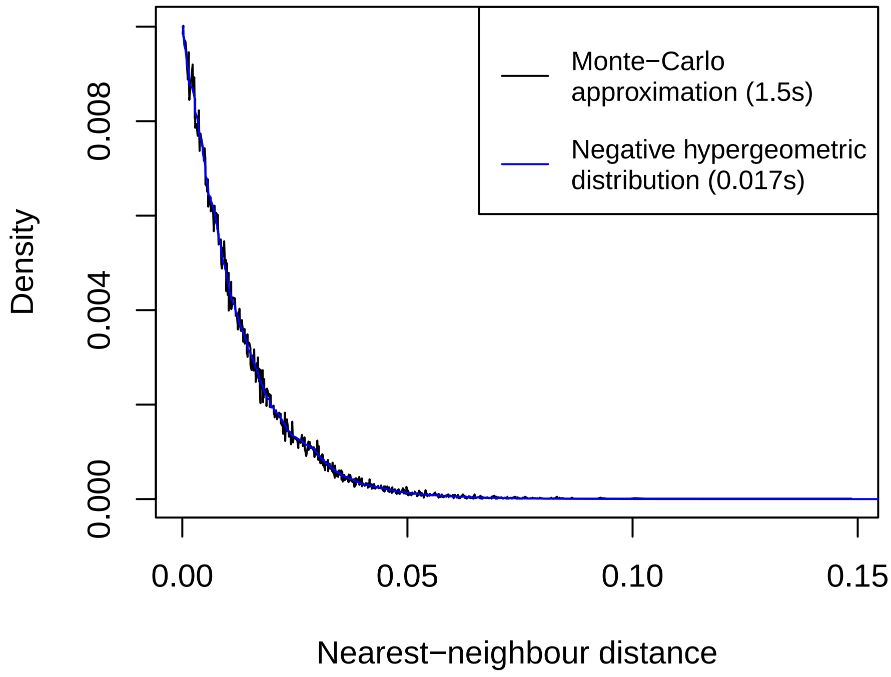

Figure S45: Densities of the nearest-neighbour distance approximated through 100,000 Monte-Carlo simulations (black) and using the analytical approach based on the negative hypergreometric distribution (blue). The computation times in seconds are shown in the legend.

##### 3.2 Gradients

The tests in *smoppix* discussed in the main text are nonparametric, as they do not propose a parametric alternative hypothesis for CSR or the background distribution, but only detect some departure from it through distances. Yet the *smoppix* package also contains a parametric test to detect gradients. The random mechanism spawning point patterns is known as a *point process*, the ones adhering to complete spatial randomness (CSR) are known as a homogeneous Poisson processes. They represent a special case of a wider class of Poisson point processes, whose likelihood function for a set of molecules with $N \times 2$ coordinate matrix $\mathbf{C}$ with elements $\mathbf{c}_i = (x_i, y_i)$ and $i = 1, \dots N$ is given by [11]

$$L(\mathbf{C}|\lambda) = \exp\left(-\int \lambda(\mathbf{C})d\mathbf{C}\right)\prod_{i=1}^{N}\lambda(\mathbf{c}_i), \tag{5}$$

with $\lambda(\mathbf{c}_i)$ an intensity parameter. For a homogeneous Poisson process, the intensity is constant over space: $\lambda(\mathbf{c}_i) = \lambda$. If an region A can be delineated, a parametric approach to detecting departures from CSR is to replace the constant intensity $\lambda$ in (5) by an intensity varying over space. We focus here on gradients as they are known to occur in tissues and cells [20]. Allowing for differences in slope and direction of the gradient across point patterns, the intensity of such an inhomogeneous Poisson process is modeled as:

$$\log\left(\lambda(\mathbf{c}_i)\right) = \gamma + \sum_{t=1}^{n}\left[\alpha_t x_i + \beta_t y_i\right] I(\mathbf{c}_i \in A_t). \tag{6}$$

The slopes $\alpha_t$ and $\beta_t$ are thus allowed to vary across point patterns, e.g. across cells. Fitting such models on multiple point patterns is implemented in the *mppm* function in the *spatstat* package [21]. The omnibus null hypothesis of CSR being valid in all point patterns is $H_0 : \alpha_1 = \alpha_2 = \ldots = \alpha_n = \beta_1 = \beta_2 = \ldots = \beta_n = 0$ and can be tested using *smoppix* with a likelihood-ratio test.

##### 3.3 A note on compositionality

Methods testing for colocalization with respect to the background distribution, such as *smoppix* and other methods that use feature label permutation, but also neighbourhood enrichment methods, are sensitive to compositional effects. When one feature is tightly clustered, it becomes more remote from a second feature. Yet as a result, all other features come nearer to this second feature compared to the background null distribution; this is the compositionality. This compositional effect is strongest in low-dimensional settings but fades as the number of features increases because then the influence of a single feature on the background distribution wanes [22], which is why we observed it in the cell type localization dataset but less in the spatial transcriptomics data. Hence when the number of features is low, we recommend to always test for univariate localization patterns first, and interpret the bivariate patterns in their light. Methods testing for localization patterns departing from CSR are insensitive to compositional effects.

#### 4 Software versions

```
## R version 4.4.0 (2024-04-24)
## Platform: x86_64-pc-linux-gnu
## Running under: Ubuntu 22.04.4 LTS
##
## Matrix products: default
## BLAS:   /usr/lib/x86_64-linux-gnu/blas/libblas.so.3.10.0
## LAPACK: /usr/lib/x86_64-linux-gnu/lapack/liblapack.so.3.10.0
##
## locale:
##  [1] LC_CTYPE=en_US.UTF-8       LC_NUMERIC=C
##  [3] LC_TIME=de_BE.UTF-8        LC_COLLATE=en_US.UTF-8
##  [5] LC_MONETARY=de_BE.UTF-8    LC_MESSAGES=en_US.UTF-8
```

```
## [7] LC_PAPER=de_BE.UTF-8       LC_NAME=C
## [9] LC_ADDRESS=C               LC_TELEPHONE=C
## [11] LC_MEASUREMENT=de_BE.UTF-8 LC_IDENTIFICATION=C
##
## time zone: Europe/Amsterdam
## tzcode source: system (glibc)
##
## attached base packages:
## [1] parallel  stats4    grid      stats     graphics  grDevices utils
## [8] datasets  methods   base
##
## other attached packages:
## [1] extraDistr_1.10.0           peakRAM_1.0.2
## [3] smoppix_0.99.20             spicyR_1.17.1
## [5] lmerTest_3.1-3              lme4_1.1-35.5
## [7] SPIAT_1.7.2                 glmnet_4.1-8
## [9] Matrix_1.7-0                SpatialExperiment_1.15.1
## [11] SingleCellExperiment_1.27.2 SummarizedExperiment_1.35.1
## [13] Biobase_2.65.0              GenomicRanges_1.57.1
## [15] GenomeInfoDb_1.41.1         IRanges_2.39.2
## [17] S4Vectors_0.43.2            BiocGenerics_0.51.0
## [19] MatrixGenerics_1.17.0       matrixStats_1.3.0
## [21] openxlsx_4.2.6.1            gplots_3.1.3.1
## [23] xtable_1.8-4                spatstat_3.1-1
## [25] spatstat.linnet_3.2-1       spatstat.model_3.3-1
## [27] rpart_4.1.23                spatstat.explore_3.3-1
## [29] nlme_3.1-165                spatstat.random_3.3-1
## [31] spatstat.geom_3.3-2         spatstat.univar_3.0-0
## [33] spatstat.data_3.1-2         psych_2.4.6.26
## [35] reshape2_1.4.4              BiocParallel_1.39.0
## [37] ggplot2_3.5.1
##
## loaded via a namespace (and not attached):
## [1] RColorBrewer_1.1-3          rstudioapi_0.16.0
## [3] jsonlite_1.8.8              shape_1.4.6.1
## [5] MultiAssayExperiment_1.31.4 magrittr_2.0.3
## [7] spatstat.utils_3.0-5        magick_2.8.4
## [9] farver_2.1.2                nloptr_2.1.1
## [11] rmarkdown_2.27              zlibbioc_1.51.1
## [13] vctrs_0.6.5                 minqa_1.2.7
## [15] tinytex_0.52                rstatix_0.7.2
## [17] htmltools_0.5.8.1           S4Arrays_1.5.6
## [19] broom_1.0.6                 SparseArray_1.5.31
## [21] KernSmooth_2.23-24          plyr_1.8.9
## [23] lifecycle_1.0.4             iterators_1.0.14
## [25] pkgconfig_2.0.3             R6_2.5.1
## [27] fastmap_1.2.0               rbibutils_2.2.16
## [29] GenomeInfoDbData_1.2.12     digest_0.6.36
## [31] numDeriv_2016.8-1.1         colorspace_2.1-1
## [33] tensor_1.5                  ggpubr_0.6.0
## [35] labeling_0.4.3              fansi_1.0.6
## [37] spatstat.sparse_3.1-0       httr_1.4.7
## [39] polyclip_1.10-7             abind_1.4-5
## [41] mgcv_1.9-1                  compiler_4.4.0
```

```
## [43] withr_3.0.1              backports_1.5.0
## [45] carData_3.0-5            highr_0.11
## [47] ggupset_0.4.0            ggforce_0.4.2
## [49] ggsignif_0.6.4           MASS_7.3-61
## [51] concaveman_1.1.0         DelayedArray_0.31.11
## [53] rjson_0.2.21             gtools_3.9.5
## [55] caTools_1.18.2           tools_4.4.0
## [57] zip_2.3.1                goftest_1.2-3
## [59] glue_1.7.0               ClassifyR_3.9.3
## [61] generics_0.1.3           gtable_0.3.5
## [63] tidyr_1.3.1              data.table_1.15.4
## [65] car_3.1-2                utf8_1.2.4
## [67] XVector_0.45.0           foreach_1.5.2
## [69] pillar_1.9.0             stringr_1.5.1
## [71] splines_4.4.0            dplyr_1.1.4
## [73] tweenr_2.0.3             lattice_0.22-6
## [75] survival_3.7-0           deldir_2.0-4
## [77] tidyselect_1.2.1         knitr_1.48
## [79] gridExtra_2.3            xfun_0.46
## [81] pheatmap_1.0.12          scam_1.2-17
## [83] stringi_1.8.4            UCSC.utils_1.1.0
## [85] yaml_2.3.10              boot_1.3-30
## [87] evaluate_0.24.0          codetools_0.2-20
## [89] tibble_3.2.1             cli_3.6.3
## [91] Rdpack_2.6               munsell_0.5.1
## [93] Rcpp_1.0.13              bitops_1.0-8
## [95] scales_1.3.0             purrr_1.0.2
## [97] crayon_1.5.3             rlang_1.1.4
## [99] mnormt_2.1.1
```
