## Supplementary figures and images for "Unified nonparametric analysis of single-molecule spatial omics data using probabilistic indices"

### Calcofluor.png

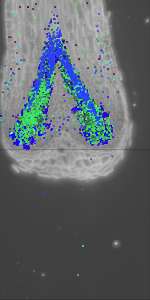
